# Systematic benchmarking of small variant calling pipelines for long-read RNA sequencing data

**DOI:** 10.64898/2026.04.29.721619

**Authors:** Jiayi Wang, Mark D. Robinson

## Abstract

**Background:** Long-read RNA sequencing (lrRNA-seq) enables transcript-resolved variant detection, but systematic and neutral evaluations of small variants calling pipelines remain limited. The performance of existing tools across sequencing technologies, alignment strategies, variant caller choice, genomic contexts and downstream haplotype phasing is not fully understood.

**Results:** Here, we systematically benchmark six lrRNA-seq variant callers (Clair3-RNA, DeepVariant, NanoTS, isoLASER, longcallR and longcallR-nn), along with a widely used short-read RNA-seq variant caller (GATK HaplotypeCaller) as a baseline, using Genome in a Bottle (GIAB) datasets comprising three cell lines sequenced with four ONT and two PacBio library preparation protocols. We further evaluate the impact of upstream alignment strategies, including aligner choice and alignment transformation, on variant-calling performance. Accuracy is assessed across sequencing depths and genomic contexts. Additionally, we compare haplotype phasing tools (WhatsHap, LongPhase, HapCUT2, HiPhase, isoLASER and longcallR) using variant calls generated by different callers to identify optimal pipeline combinations. Finally, we extend our evaluation of variant-calling performance to more recent LongBench datasets.

**Conclusions:** Our benchmark shows that sequencing characteristics are the strongest determinant of lrRNA-seq variant-calling performance, followed by variant caller and alignment strategies, with additional effects from genomic context. Variant caller strengths differed by platform and variant type: Clair3-RNA showed the broadest and most robust applicability across ONT and PacBio chemistries; NanoTS performed best or near-best for SNVs on newer chemistries; longcallR was competitive for SNV calling; and DeepVariant performed strongly on PacBio, particularly for indels. For downstream haplotype phasing, no single method excelled across all metrics. HiPhase and LongPhase achieved the lowest switch error rates on PacBio and ONT data, respectively; WhatsHap maximized coverage and contiguity; and HapCUT2 balanced accuracy and coverage for SNVs. LongcallR offers a strong alternative for joint variant calling and phasing.

## Background

Accurate detection of small variants (single nucleotide variants (SNVs) and insertions/deletions (indels)) underpins our ability to link genetic changes with disease mechanisms, therapeutic responses and clinical outcomes^[1]^. Next generation sequencing is the most widely used approach for variant calling because it enables efficient large scale screening of genetic variants. Often, DNA sequencing (DNA-seq), including whole genome and whole exome sequencing, is used, as it provides comprehensive and relatively unbiased coverage of the genome. Previous studies have also demonstrated the promise of applying RNA sequencing (RNA-seq) for variant calling^[2][3][4]^. However, RNA-seq is limited by expression dependence and biases such as uneven transcript abundance, allele-specific expression, and splicing-related alignment artifacts. Despite these limitations, RNA-seq offers several advantages compared to DNA-seq. It is more cost effective and delivers higher depth for a given read count because the transcriptome spans a much smaller sequence space, while also resolving multiple isoforms of individual genes. In addition, it enables detection of post transcriptional events and simultaneous assessment of genotype and phenotype.

Recent advances in long-read RNA sequencing (lrRNA-seq) offer added power for variant analysis, as it captures full-length transcripts that are not accessible with short-read approaches. This enables direct linkage of detected variants to specific transcript isoforms, thereby facilitating analysis of allele-specific events and splicing. Long reads are also more effective for haplotype phasing since single reads span multiple SNVs^[5]^. Pacific Biosciences (PacBio) developed Iso-Seq to sequence full-length cDNA via SMRT Cell sequencing technology^[6]^, which was later improved by concatenating cDNA molecules into longer fragments, a method known as Mas-Seq (commercialized as Kinnex)^[7]^. The latest Kinnex kits further extend this strategy by concatenating shorter amplicons into larger fragments to increase throughput. Another leading platform, Oxford Nanopore Technology (ONT), supports both cDNA and direct RNA (dRNA) sequencing^[8]^; dRNA examines native RNA without enzymatic conversion and allows detection of RNA modifications and RNA epigenetics analysis^[9]^.

Before lrRNA-seq became widely adopted, several lrDNA-seq variant callers were available, including DeepVariant^[10]^, Clair3^[11]^ and NanoCaller^[12]^. However, these tools were designed for contiguous DNA alignments and did not at the time support processing of spliced RNA-seq reads. A recent study proposed a pipeline that transforms lrRNA-seq alignments by splitting reads at exon-intron joints and generating contiguous alignments^[13]^. With this transformation, lrDNA-seq variant callers are compatible with lrRNA-seq reads and achieve reasonable accuracy to call SNVs and indels. More recently, variant callers have been optimized for lrRNA-seq, including Clair3-RNA^[14]^, the DeepVariant MASSEQ model, isoLASER^[15]^, NanoTS^[16]^, longcallR and longcallR-nn^[17]^. To our knowledge, systematic neutral evaluations comparing their performance and assessing the continued necessity of alignment file transformation across different lrRNA-seq technologies are still lacking.

Following variant calling, heterozygous variants can be phased into haplotypes, defined as sets of variants co-inherited on the same chromosome. Several tools have been developed for this purpose, including WhatsHap^[18]^, HapCUT2^[19]^ and LongPhase^[20]^. A recent study benchmarked phasing tools using the variant calls generated by Clair3-RNA on PacBio Iso-Seq data^[21]^. However, a comprehensive evaluation of phasing performance across diverse lrRNA-seq platforms and variant callers has yet to be conducted.

In this study, we systematically benchmarked six lrRNA-seq variant callers and one short-read RNA-seq variant caller as a baseline using GIAB datasets, comprising three cell lines sequenced with four ONT and two PacBio library kits. We evaluated the impact of upstream alignment strategies, including the choice of aligner and the application of alignment transformation, on variant calling performance. Variant calling accuracy was further assessed across different read depths and genomic contexts. Additionally, we compared haplotype phasing tools using variant calls generated by different callers to identify optimal pipeline combinations. Finally, we extended our variant calling evaluation to more recent LongBench^[22]^ datasets.

## Results

### Benchmark design

Detecting small variants from RNA-seq typically follows the process of alignment, variant calling, filtering and optional haplotype phasing. The most commonly used aligner for lrRNA-seq data is minimap2^[23]^, which provides a splice-aware mode to map reads to reference genome across exon–exon junctions. Another widely used aligner (for PacBio data) is pbmm2, a wrapper around minimap2 that optimizes alignment parameters, generates sorted output files, and performs additional post-processing steps for PacBio reads (Fig. 1a). We included both aligners to evaluate whether the parameter optimization in pbmm2 improves variant calling performance. Other splice-aware aligners for lrRNA-seq include *deSALT* ^[24]^ and *uLTRA*^[25]^. In preliminary testing, we found that *uLTRA* was considerably more computationally demanding, while *deSALT* has not been updated in recent years and is less commonly used. Given the popularity of minimap2 and pbmm2, and the fact that they form the basis of most variant calling pipelines, we focused our analysis on these two aligners. After alignment, transformation of the alignment files had been recommended to mimic contiguously aligned DNA reads, thereby improving variant calling accuracy^[13]^. Building on this, we tested whether this transformation step remains necessary for variant callers specifically designed for lrRNA-seq data.

**Fig. 1:**
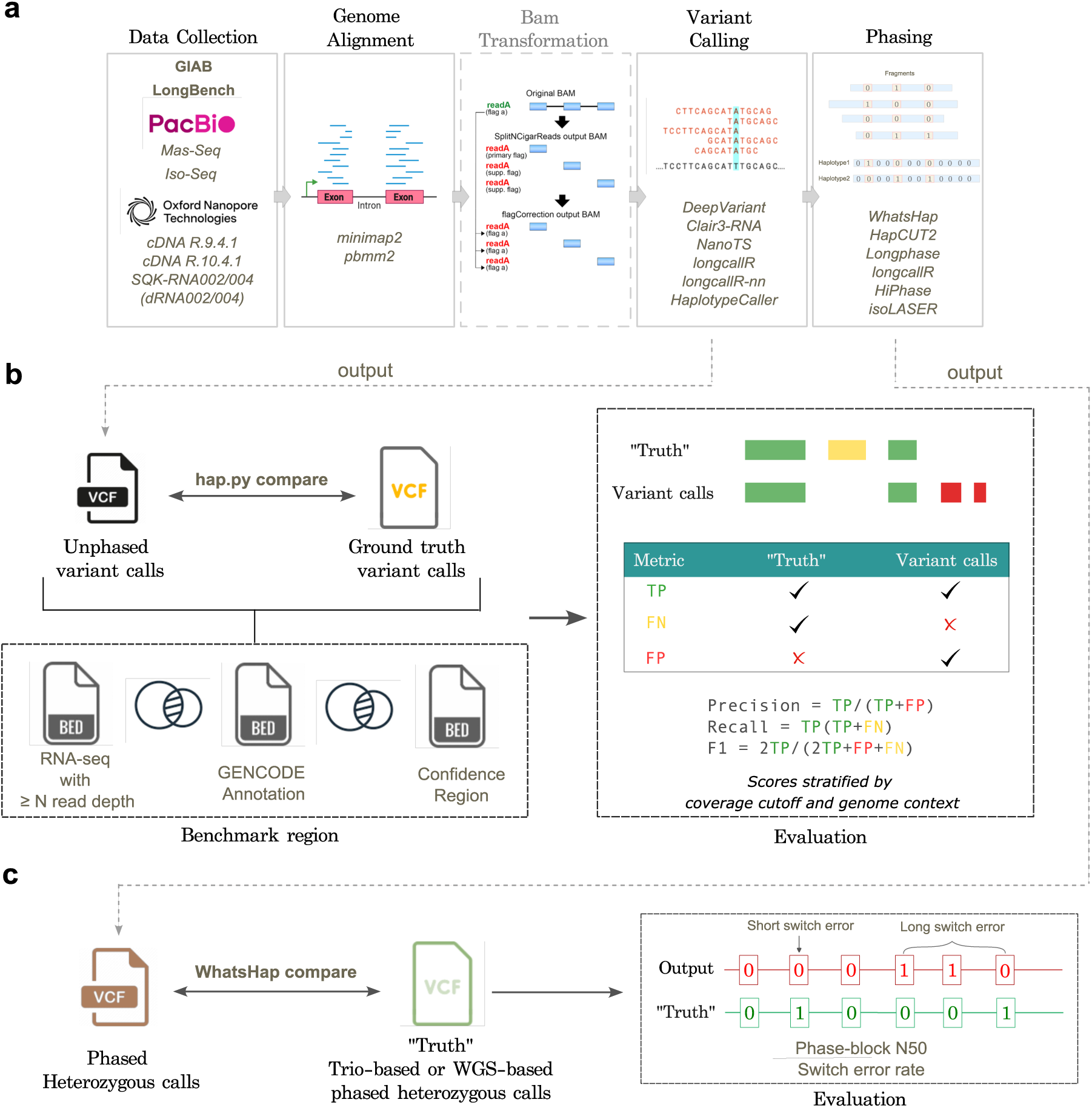
Schematic of the variant calling and phasing benchmarking workflow. (**a**) Overall analysis workflow. (**b**) Evaluation of variant calling results. Variant calls were compared with ground-truth calls within regions overlapping RNA-seq data (above a defined coverage cutoff), GENCODE annotations, and GIAB confidence regions. Performance was quantified using precision, recall, and F1 scores, stratified by sequencing coverage and genomic region. (**c**) Evaluation of phasing results. Phased variant calls were compared with ground-truth phased calls, and performance was assessed using several metrics including phase-block N50 and switch error rate.

Next, the genome-aligned reads, both transformed (where noted) and untransformed, were used as input for variant calling, focusing on callers developed for lrRNA-seq: Clair3-RNA^[14]^, DeepVariant^[10]^, isoLASER^[15]^, NanoTS^[16]^, longcallR and longcallR-nn^[17]^ (Table 1, see Methods). GATK HaplotypeCaller^[26]^ was included as a widely used standard (for short-read RNA-seq) to provide a baseline for comparison (Fig. 1a). To date, Clair3-RNA, isoLASER, NanoTS, longcallR-nn, and longcallR support both PacBio and ONT platforms through trained models or parameter presets, while DeepVariant does not offer a model for ONT RNA-seq. Accordingly, we applied the ONT DNA-seq model with the additional arguments recommended by the DeepVariant developers (see Methods). Each variant caller assigns a quality score and threshold to filter low-confidence calls. This typically depends on multiple factors such as alignment quality, base quality, strand bias, and the presence of RNA-editing sites.

**Table 1:** Feature comparison of variant callers benchmarked in this study. Algorithm abbreviations: CNN = Convolutional Neural Network; Bi-LSTM = Bidirectional Long Short-Term Memory; PairHMM = Pair Hidden Markov Model; MLP = Multilayer Perceptron. *: Optional phasing with WhatsHap or LongPhase, not available for ONT dRNA002 and cDNA R9 libraries. *^†^*: authors recommended indel calling only for PacBio data. *^‡^*:Phasing is performed with WhatsHap; NanoTS uses the haplotype-split reads to re-genotype variants.

| Method | Algorithm | Pretrained<br>model | Variant<br>type | Phased<br>output | lrRNA-seq<br>support |
|---|---|---|---|---|---|
| Clair3-RNA<sup>[14]</sup> | CNN + Bi-LSTM | ✓ | SNV/INDEL | optional* | PacBio & ONT |
| DeepVariant<sup>[10]</sup> | CNN | ✓ | SNV/INDEL | ✗ | PacBio |
| longcallR-nn<sup>[17]</sup> | CNN | ✓ | SNV only | ✗ | PacBio & ONT |
| longcallR<sup>[17]</sup> | Heuristic + Prob. | ✗ | SNV only | ✓ | PacBio & ONT |
| isoLASER<sup>[15]</sup> | Local reassembly + MLP | ✓ | SNV/INDEL<sup>†</sup> | ✓ | PacBio & ONT |
| NanoTS<sup>[16]</sup> | CNN + MLP | ✓ | SNV only | both<sup>‡</sup> | PacBio & ONT |
| GATK HaplotypeCaller<sup>[26]</sup> | PairHMM | ✗ | SNV/INDEL | ✗ | Short-read |

We evaluated variant calling performance using datasets from three Genome in a Bottle (GIAB) Consortium reference cell lines, HG002, HG004 and HG005. Each cell line was generated using six different library preparation kits and sequenced on both Oxford Nanopore Technologies (ONT; cDNA R9.4.1 and R10.4.1 chemistries, and dRNA002 and dRNA004) and PacBio (Mas-Seq and Iso-Seq) platforms (Table S1). Performance was quantified by comparing filtered variant calls to ground-truth calls within high-confidence regions, which represent variants integrated from multiple sequencing technologies for the same cell line. By stratifying performance by sequencing depth or genomic context, we assessed how these factors influence the accuracy and sensitivity of variant detection (Fig. 1b).

Finally, filtered heterozygous variant calls from the output VCF files for the HG002 and HG005 cell lines were phased using WhatsHap^[18]^, LongPhase^[20]^, HapCUT2^[19]^, HiPhase^[27]^ and longcallR. Haplotypes were directly inferred from aligned BAM files, also called read-based phasing (Fig. 1a, Table 2). The phased outputs were compared to high-confidence trio-based or whole-genome sequencing-based phased VCFs as ground truth (Fig. 1c). Performance was assessed using switch error rate and phasing coverage.

**Table 2:** Feature comparison of phasing tools benchmarked in this study. Algorithm abbreviations: wMEC = Weighted Minimum Error Correction; Greedy = graph-based greedy phasing; Max-Cut = Maximum Cut on fragment graphs.

| Method | Algorithm | Support Platform | lrRNA-seq-specific preset |
|---|---|---|---|
| WhatsHap<sup>[18]</sup> | wMEC | PacBio & ONT | ✗ |
| LongPhase<sup>[20]</sup> | Greedy graph-based | PacBio & ONT | ✗ |
| HapCUT2<sup>[19]</sup> | Max-Cut on fragment graph | PacBio & ONT | ✗ |
| HiPhase<sup>[27]</sup> | A* search on read graphs | PacBio | ✓ |
| isoLASER<sup>[15]</sup> | Weighted *k*-means read clustering | PacBio & ONT | ✓ |
| longcallR<sup>[17]</sup> | Iterative probabilistic refinement | PacBio & ONT | ✓ |

### Sequencing characteristics drive variant calling performance

We first examined overall variant-calling performance across platforms and libraries (Fig. 2a, Fig. S1a). All variant callers showed high accuracy on PacBio datasets, where most achieving their highest F1 scores across all sequencing chemistries for each cell line. In contrast, performance was consistently poorest for the ONT dRNA002 chemistry (Fig. 2b-d, Fig. S1b,c). These differences are attributable to variations in data quality arising from the use of different library preparation kits. We observed a clear relationship between sequencing error rate and variant calling accuracy: higher error rates reduce the signal-to-noise ratio of read pileups used by variant callers (Table S3, Fig. S2a). Although variant callers apply base and alignment quality filters, residual base-calling and alignment errors can still introduce spurious variants or obscure true variants, leading to both FPs and FNs. One notable exception was the cDNA R10 datasets, which exhibited higher error rates than the cDNA R9 and dRNA004 datasets, likely reflecting characteristics of the specific public datasets rather than the intrinsic performance of the sequencing chemistry.

**Fig. 2:**
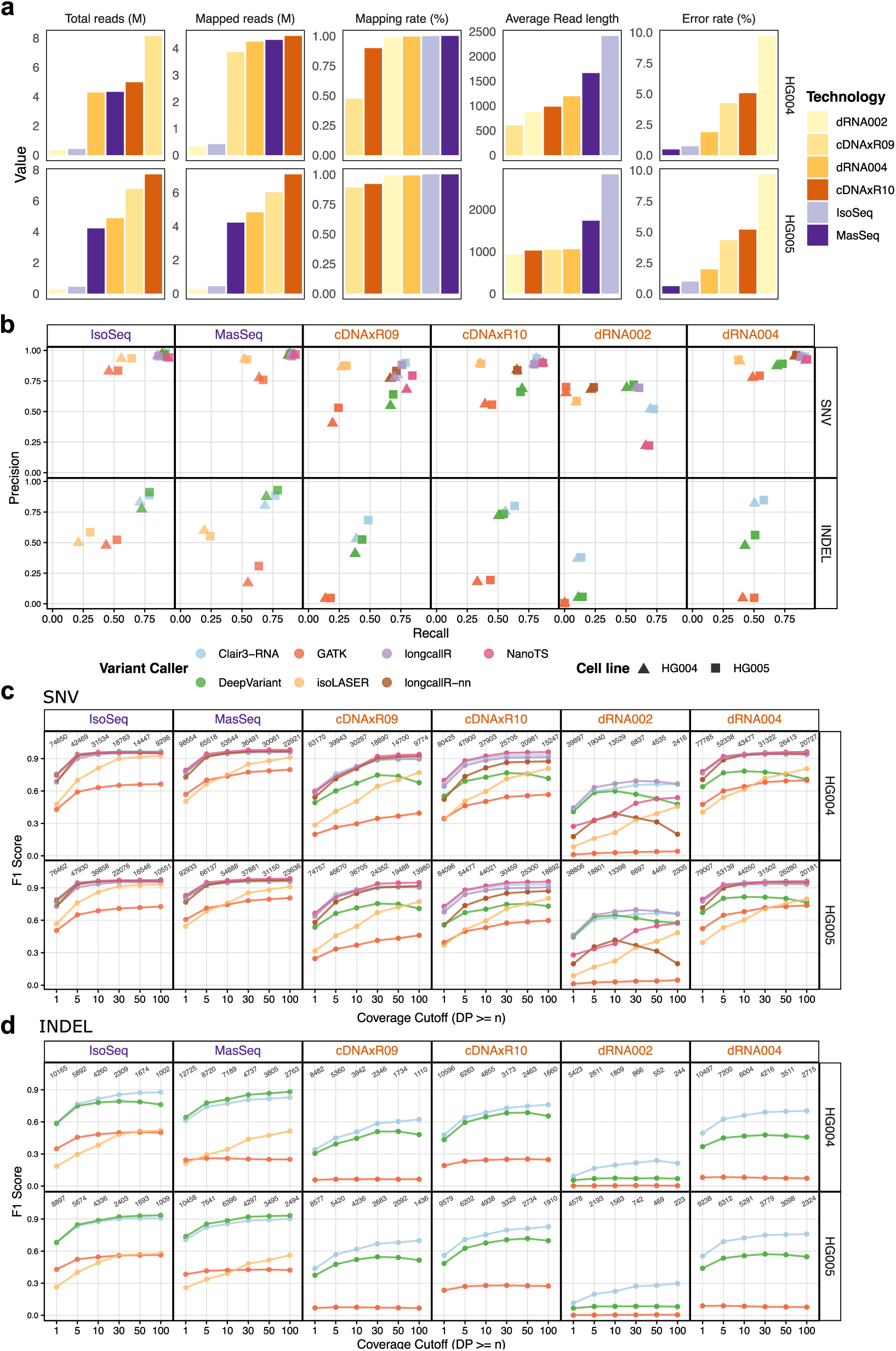
Data quality and variant calling performance. (**a**) Bar plots showing data quality, colored by library preparation kit and stratified by quality metric and cell line. Total sequences and mapped reads are shown in units of millions. Mapping rate is based on minimap2 alignment. Sequencing error rate refers to the ratio between mismatches and bases mapped (cigar). (**b**) Precision and recall performance for each variant caller at 5*×* coverage, stratified by variant type (SNVs and indels) and library preparation kit. (**c, d**) F1 scores for calling SNVs (**c**) and indels (**d**) across different coverage, stratified by cell lines (HG004 and HG005) and library preparation kits. Results for HG002 cell line are shown in Fig. S1. The numbers above the dots indicate the number of benchmark truth variants at each coverage cutoff for each sequencing chemistry. All variant calls were based on minimap2 alignments and without BAM transformation. Panel **b**, **c** and **d** use the same color legend. Purple stands for PacBio technologies and orange stands for ONT technologies.

Together with accuracy, the total number of called variants provides a complementary metric that reflects sequencing throughput, mapping rate, and read length, which together determine the number of callable bases available for variant calling. Sequencing throughput directly determines how much of the transcriptome can be confidently analyzed, and longer reads improve mapping accuracy and expand callable regions. Accordingly, Mas-Seq datasets retained the highest number of callable bases across all coverage thresholds, followed by ONT dRNA004, cDNA R10, Iso-Seq, cDNA R9, and dRNA002 (Fig. S3). As a result, datasets with more callable bases consistently yielded more TP variants, with TP counts strongly correlated with the number of callable bases across all callers and coverage thresholds (Spearman’s *ρ* 0.85–1.0; Table S4, Fig. S2b). Although ONT cDNA sequencing produces higher throughput than PacBio, its lower mappability and shorter reads considerably limit the number of detectable variants (Fig. S4, Fig. S5). Consequently, PacBio datasets yielded *∼*1.6-fold more true variants than ONT cDNA datasets (49,953 vs. 31,299 at 5*×* coverage, averaged across cell lines and lrRNA-seq–specific callers), despite starting from fewer raw reads. Due to its high sequencing error rate, limited throughput, and poor mappability, the dRNA002 dataset failed to recover a substantial proportion of variants. Among ONT library kits, dRNA004 and cDNA R10 showed the strongest performance, with average SNV F1 scores of *∼*0.84 and *∼*0.75, respectively, across all cell lines and lrRNA-seq–specific variant callers at 5× coverage, but remained below PacBio (*∼*0.88 for Mas-Seq and Iso-Seq). In PacBio, Mas-Seq achieves 10.3-fold higher throughput (in reads) than Iso-Seq via fragment concatenation, yielding the highest number of detected true variants (on average 59,367 vs. 40,540).

To further disentangle the effects of callable bases and sequencing accuracy, we sub-sampled the HG004 Mas-Seq reads to match the number of callable bases in the HG004 dRNA004 dataset using five random seeds (Fig. S6a) and evaluated variant calling with Clair3-RNA. At the 5*×* coverage cutoff, the five subsampled Mas-Seq datasets showed similar F1 scores and TP counts for SNV calling compared with the original dRNA004 dataset, whereas Mas-Seq retained higher performance for indel calling. At the 100*×* cutoff, dRNA004 had *>*140 million more callable bases than the subsampled Mas-Seq datasets and consequently yielded more TP calls, but its F1 score remained lower (Fig. S6b-d).

Overall, sequencing characteristics play a central role in variant calling performance, with sequencing depth, mapping rate, read length, and error rate jointly contributing to the consistently stronger performance observed for PacBio compared to ONT. To quantify the relative contributions of different factors, we fitted linear models including sequencing chemistry, variant caller, variant type, coverage cutoff, and cell line. Sequencing chemistry explained the largest proportion of variation in both F1 score (partial *R*^2^ = 0.784) and the number of TP calls (partial *R*^2^ = 0.791), exceeding the contribution of variant caller (partial *R*^2^ = 0.726 and 0.312, respectively; Table S5). In addition, the regression coefficients confirmed PacBio Mas-Seq and Iso-Seq as the highest-performing chemistries, whereas ONT dRNA002 showed the largest reduction in both F1 score and TP calls relative to the ONT cDNA R9 reference (Table S6). Our subsampling analysis further showed that, when the number of callable bases was similar, higher sequencing accuracy improved TP recovery, particularly for indels. Deeper sequencing could largely compensate for lower accuracy in SNV calling, but less so for indels, where sequencing accuracy remained important for F1 scores.

### Variant callers tailored for lrRNA-seq showed strong performance

We next evaluated the performance of the variant callers. Clair3-RNA, DeepVariant, isoLASER, NanoTS and longcallR-nn rely on platform- and/or library-specific pre-trained models. Although longcallR does not use pre-trained models, it depends on preset parameter configurations tailored to the sequencing platform and library. Among the evaluated datasets, the HG002 datasets were used for training by multiple variant callers, whereas HG004 and HG005 Iso-Seq and ONT datasets were training-independent (Table S2). Because DeepVariant was trained on Mas-Seq data from all three GIAB cell lines, with chromosome 20 excluded during training, we additionally report chromosome 20-only results for the HG004 and HG005 Mas-Seq datasets (Fig. S1d).

LrRNA-seq-specific variant callers outperformed the baseline GATK HaplotypeCaller in most cases (Fig. 2b-d, Fig. S1b, c). For SNV calling in HG004 and HG005 ONT datasets, Clair3-RNA, NanoTS and longcallR were generally the best-performing methods, achieving similar F1 scores at 5× coverage across cDNAxR09 (mean F1 0.80, 0.78 and 0.78), cDNAxR10 (0.87, 0.88 and 0.84), and dRNA004 (0.93, 0.92 and 0.91) (Fig. 2c). NanoTS identified the most TPs on these three chemistries, followed by Clair3-RNA, but at the cost of more false positives on cDNAxR09, where its precision was lower (Fig. S4, Fig. S5). On dRNA002 specifically, longcallR slightly outperformed Clair3-RNA in both HG004 and HG005 (F1 0.63–0.65 [95% CI 0.622–0.657] vs. 0.60–0.61 [95% CI 0.587–0.615] at 5×, Table S7). LongcallR-nn also showed competitive performance in several ONT datasets but performed poorly on the dRNA002 datasets. This is because its dRNA model was trained exclusively on dRNA004, as well as NanoTS. DeepVariant consistently performed poorly on ONT datasets, as expected given the lack of an ONT-specific RNA-seq model (Fig. 2c, d, Fig. S1). Despite being designed specifically for lrRNA-seq, isoLASER only outperformed DeepVariant at *≥*50–100*×* on most ONT chemistries. This gap was driven by its low recall (0.59–0.66 vs. 0.64–0.71 at 50*×*), although isoLASER consistently achieved higher precision (0.88–0.92 vs. 0.80–0.92). For indel calling, Clair3-RNA was the only method capable of robust indel detection in ONT datasets and therefore the best-performing tool for this task (Fig. 2d, Fig. S1b, Table S8).

Across HG004 and HG005 PacBio datasets, Clair3-RNA and DeepVariant were the top-performing methods for both SNV and indel calling, joined by NanoTS for SNVs. For SNVs at 5× coverage, NanoTS performed best on Mas-Seq (mean F1 0.947) and Clair3-RNA on Iso-Seq (0.945). They were closely followed by Clair3-RNA (0.935) and DeepVariant (0.930) on Mas-Seq, and by NanoTS (0.941) and DeepVariant (0.938) on Iso-Seq, while longcallR and longcallR-nn trailed by 0.01–0.04 F1, a gap that narrowed substantially at higher coverage (Fig. 2c-d, Table S7). By 10×, all lrRNA-seq callers except isoLASER achieved F1 *≥* 0.93 on PacBio datasets. NanoTS also recovered the most TP SNVs on both Mas-Seq and Iso-Seq (mean 61,154 and 42,580 at 5×), but on Iso-Seq this came with more FPs than Clair3-RNA, explaining its slightly lower F1 (Fig. S4). IsoLASER achieved competitive F1 scores for SNV calling at high coverages but was conservative, resulting in lower recall. For indels at 5× coverage, DeepVariant showed slightly higher mean F1 than Clair3-RNA on Mas-Seq (0.815 vs 0.782). This advantage was driven mainly by fewer FPs (mean 668 vs 1,120), as both callers recovered a similar number of TP indels (6,058 vs 5,935). Because all Mas-Seq datasets were involved in DeepVariant’s training (Table S2), we restricted the comparison to chr20, which is excluded from training, and observed a similar pattern (mean F1 0.790 vs 0.758) (Fig. S1d), indicating that DeepVariant’s indel-calling advantage on PacBio is not solely attributable to training-data overlap.

The relative performance of the callers was robust to preprocessing and variant allele fraction. Restricting the evaluation to loci sufficiently covered after isoLASER preprocessing (see Methods) did not change the ranking of callers (Fig. S7). Users interested in detecting low-allele-fraction variants should carefully consider caller-specific performance in this range, although these variants accounted for only 1–3% of truth SNVs. For the remaining allele fractions, caller differences were consistent with the overall results, and the ranking was also robust to quality-score thresholds, as shown by precision–recall curves (Fig. S8).

Overall, no single caller dominated across all chemistries and variant types. For ONT data, Clair3-RNA offered the most consistent performance, supporting both SNV and indel calling across all chemistries. NanoTS achieved the highest SNV accuracy on several ONT and PacBio datasets but is limited to SNVs and performed poorly on dRNA002, and longcallR was a strong alternative for SNV calling on ONT without reliance on pre-trained models. For PacBio data, NanoTS, Clair3-RNA and DeepVariant performed well across coverages, with DeepVariant showing a modest edge for indel calling. Taken together, Clair3-RNA is the most versatile tool for lrRNA-seq variant calling, while NanoTS, DeepVariant and longcallR(-nn) can offer equal or better alternatives in specific contexts.

### Minimap2 improves variant calling performance compared to pbmm2

Recognizing that read mappability is a key determinant of variant-calling performance, we evaluated the performance of using pbmm2 versus minimap2. Across most caller and dataset combinations, we observed that pbmm2 detected more variants but yielded a slightly lower F1 score than minimap2. Notably, isoLASER was the only exception, with pbmm2 yielding higher F1 despite lower precision, driven by a substantial gain in recall, which may reflect its interaction with isoLASER’s mandatory transcriptome re-alignment step (Fig. S9, Fig. S10). Specifically, pbmm2 tended to introduce more false calls, particularly FPs, which resulted in higher false discovery rates than minimap2 across most datasets (Fig. 3a). This finding is consistent with observations reported by the developers of Clair3-RNA^[14]^.

**Fig. 3:**
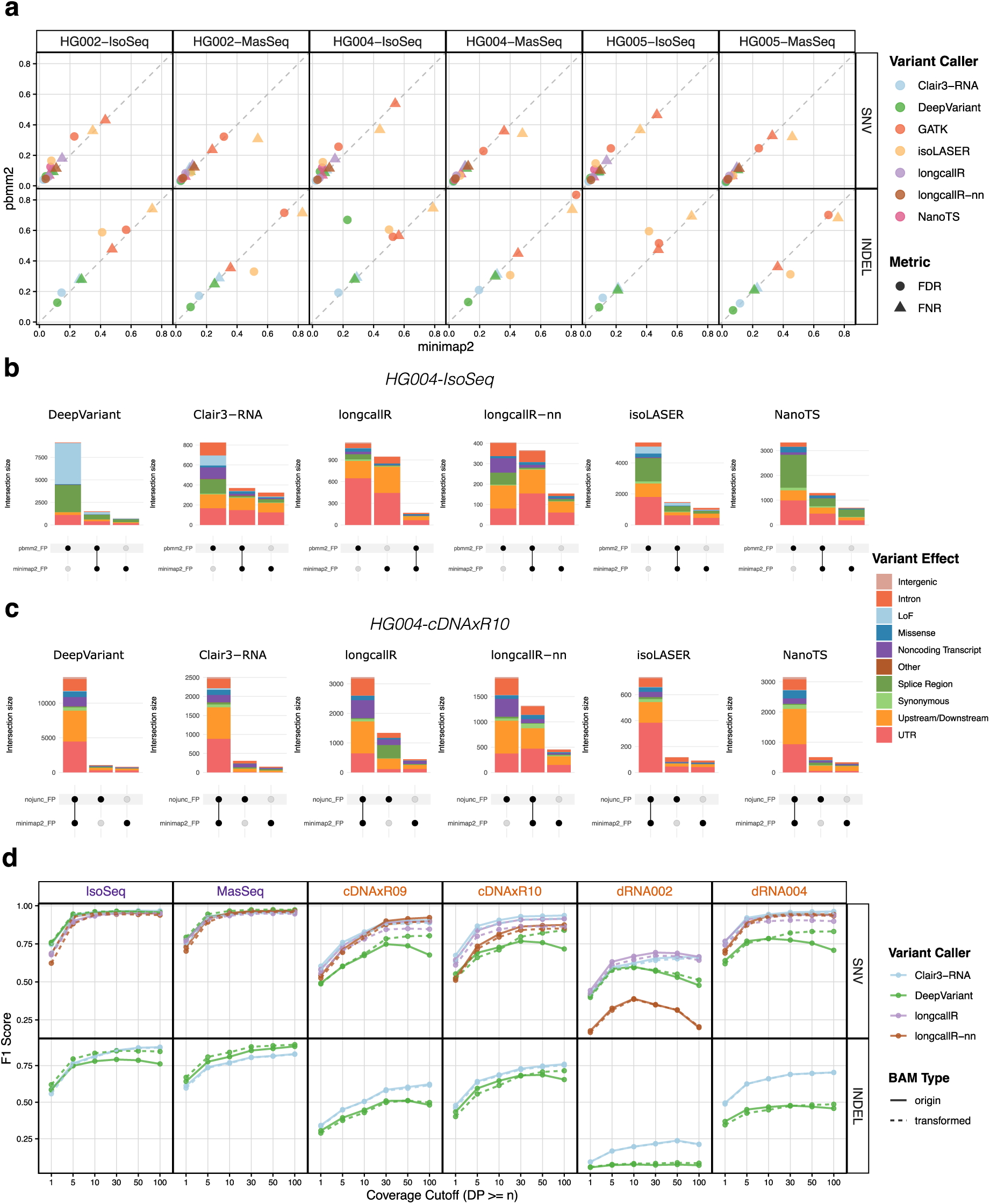
Impact of aligner and BAM file format on variant calling performance. (**a**) Scatter plot comparing the false discovery rate (FDR) and false negative rate (FNR) on variant calling results based between minimap2 and pbmm2 alignments, colored by aligner and stratified by datasets and variant types. The dashed diagonal line indicates equal performance between aligners. (**b**) UpSet plots illustrating the genomic context of aligner-specific and shared FP calls, stratified by variant caller, for HG004 Iso-Seq dataset. (**c**) The same as panel (**b**), but comparing minimap2 with and without the splice-junction annotation file as input for HG004 cDNA R10 dataset. minimap2-nojunc means minimap2 alignment without transcriptome annotation BED file passing for --junc-bed. All alignments in panel **a-c** are without BAM transformation. (**d**) F1 scores for SNV and indel calling in the HG004 datasets across different coverage cutoffs, stratified by library preparation kit and variant type. Solid and dashed lines represent results obtained without (origin) and with (transformed) BAM file transformation, respectively. Purple stands for PacBio technologies and orange stands for ONT technologies.

To understand these differences, we inspected the genomic locations of FP calls that were specific to or shared between minimap2 and pbmm2. The HG004-IsoSeq dataset exhibits the strongest aligner-dependent differences across all variant callers. In this dataset, pbmm2-specific FPs form the largest set (excluding shared TPs) for most variant callers. Among these FPs, the proportion of non-coding transcript and splice-region variants is higher than in the other sets (Fig. 3b). This elevated proportion of splice-region variants in pbmm2-specific FPs was also observed across other caller–dataset combinations (Fig. S11). We also found slightly higher rates of supplementary alignments with pbmm2 (Table S9), but no clear relationship between supplementary alignment rates and the number of FPs. We further selected a pbmm2-specific splice-region FP SNV (chr19:39,457,741 G/A) in HG004-IsoSeq and inspected the alignments on Integrative Genomics Viewer (IGV). pbmm2 appeared less willing to leave read bases unaligned: at splice junctions, it introduced small indels, while at alignment ends, it extended through mismatches rather than soft-clipping (NM = 10 vs. 18 for an identical read) (Fig. S12). Small differences in exon–intron boundary placement can shift a few bases between intronic and exonic alignment, resulting in apparent mismatches or short indels near splice sites. Such local alignment differences can generate consistent mismatch patterns in pbmm2 that are subsequently interpreted as variants by downstream variant callers, thereby producing more FP calls.

Although pbmm2 and minimap2 rely on the same underlying alignment algorithm, pbmm2 applies PacBio-optimized parameter presets that may influence splice-aware alignment behavior (Table S10). The most striking difference is that pbmm2 does not take splice-junction annotations as input and instead identifies introns from the genome sequence alone, whereas minimap2 is provided with a list of known junctions (--junc-bed) and adds a bonus to alignments that use them (--junc-bonus). We therefore compared minimap2 alignments with and without the input annotation file on the HG004 datasets. Removing the annotation file had little effect on variant-calling performance, as measured by F1 score, in most cases, although a small improvement was observed for isoLASER (Fig. S13, Fig. S14). Alignments without splice-junction annotations usually produced slightly more FPs, although these were not clearly enriched at splice sites except for longcallR (Fig. 3c, Fig. S15). Therefore, the lack of splice-junction annotations in pbmm2 may account for a small part of the difference between pbmm2 and minimap2, with the remaining differences likely attributable to parameter settings.

Ultimately, however, overall performance is largely determined by how the variant caller interprets and filters alignment-derived signals. For example, the FP SNV described above was filtered by DeepVariant but retained by longcallR. Overall, pbmm2 yields more mapped bases and consequently more variant calls, but at the cost of increased false positives and a slightly lower F1 score compared with minimap2.

### BAM file transformation does not improve performance for lrRNA-seq variant callers

As mentioned above, transforming RNA-seq alignments into a DNA-like representation improved the performance of lrDNA-seq variant callers on lrRNA-seq data^[13]^. Given the substantial computational cost of this preprocessing step, we evaluated whether it remains beneficial for lrRNA-seq–specific variant callers using HG004 datasets. Of the three supervised methods, DeepVariant MASSEQ model was trained on both original and transformed BAM formats, while Clair3-RNA and longcallR-nn were trained only on original alignments. IsoLASER was not included in this analysis because it performs variant calling directly from transcriptome alignments, rendering the genome-alignment transformation evaluated here inapplicable. NanoTS was also excluded because it requires the NM (edit distance) tag to calculate the per-read mismatch rate for filtering, but this tag is removed during the transformation.

At a read depth of 5, alignment transformation slightly improved the precision and recall of DeepVariant on PacBio datasets, resulting in average F1 score increases of 0.014 for SNVs and 0.025for indels. This improvement was more pronounced for indel calling in Iso-Seq datasets at higher coverage (Fig. S16). However, in other scenarios, alignment transformation did not yield consistent performance gains for DeepVariant across different coverage levels and, in some cases, was associated with reduced performance for Clair3-RNA, longcallR, and longcallR-nn (Fig. 3d). We hypothesize that the reduced performance observed for these lrRNA-seq–specific variant callers may be attributable to the disruption of RNA-specific alignment features, such as splice-aware CIGAR patterns and transcript-context information, which these methods are designed to exploit. In addition, the SplitNCigarReads transformation artificially increases the number of read termini by splitting reads at splice junctions, which may reduce variant calling accuracy for variants located near the newly created read ends. Splitting long reads at introns may also remove haplotype information spanning multiple exons, thereby reducing the long-range evidence available to distinguish true variants from sequencing errors. Together, these factors may contribute to the modest performance decrease observed following alignment transformation.

Therefore, we suggest that alignment transformation can be omitted when using lrRNA-seq–specific variant callers for ONT datasets, while it may still be beneficial when applying DeepVariant to PacBio datasets, particularly for detecting indels.

### LrRNA-seq-specific callers differ in their ability to identify RNA editing sites

Unlike DNA sequencing, RNA-seq is affected by RNA editing, particularly adenosine-to-inosine (A-to-I) editing mediated by ADAR enzymes^[28]^. These A-to-I editing events appear in RNA-seq data as A-to-G or T-to-C substitutions and may lead to an increased FP rate. Supervised methods are trained using DNA-derived labels, allowing the models to implicitly distinguish RNA editing sites from true genomic variants^[4]^. For example, NanoTS labels candidate sites absent from the GIAB truth set as reference during training^[16]^. In addition, longcallR-nn incorporates the REDIportal database^[29]^ during training, treating A-to-I editing as a separate class, and Clair3-RNA also queries REDIportal at inference and tags matching variants as RNA editing sites, leaving their removal to the user. In contrast, longcallR distinguishes true variants from RNA editing events through iterative read phasing and haplotype consistency testing. Here, we overlapped FP A-to-G or T-to-C calls that passed the quality filter and were not annotated as RNA editing sites with REDIportal to evaluate the ability of lrRNA-seq-specific callers to distinguish genomic variants from known RNA editing events.

We observed a lower proportion of known RNA editing sites among A-to-G or T-to-C FP SNVs across all variant callers on the dRNA datasets, consistent with A-to-I editing being more readily represented as A-to-G substitutions after reverse transcription to cDNA^[30]^. By directly tagging RNA editing sites using the built-in REDIportal database, Clair3-RNA had the lowest proportion and absolute number of RNA editing sites among A*>*G/T*>*C false-positive SNVs in most datasets (Fig. 4a, Fig. S17a). As expected, DeepVariant showed the highest number of RNA editing sites in most ONT datasets when using its DNA-seq model. In contrast, DeepVariant showed a lower proportion of RNA editing sites in the PacBio datasets, suggesting that its MASSEQ model has learned to distinguish RNA editing sites from true genomic variants. NanoTS also had a low proportion of known RNA editing sites in ONT cDNA datasets (*∼*9% vs. *∼*6% for Clair3-RNA) without using the REDIportal database, but this increased to *∼*51% in Iso-Seq data with its Mas-Seq model. In ONT cDNA datasets, a notable fraction of A-to-G or T-to-C FP SNVs from isoLASER, longcallR, and longcallR-nn overlapped known RNA editing sites (32–63%), indicating that A-to-I editing contributes to their FP burden both for this substitution class and overall (13–34% of all FP SNVs; Fig. S17b). To assess whether filtering known editing sites would be beneficial in practice, we removed all calls matching a REDIportal site from each call set before benchmarking. This improved precision for DeepVariant in the ONT datasets from 0.706 to 0.745 on average at 5*×* coverage cutoff (Fig. 4b, Fig. S18a). Moderate improvements were also observed for longcallR-nn, isoLASER, and longcallR (0.01–0.02), whereas precision was unchanged for Clair3-RNA, which already filters these sites. However, the filter also removed true variants that coincide with catalogued editing sites, reducing recall by 0.4–1.2% on average for all callers (Fig. 4c, Fig. S18b). The net effect on F1 was therefore small (*−*0.005 to +0.014), and the relative ranking of callers remained largely unchanged (Fig. S18c).

**Fig. 4:**
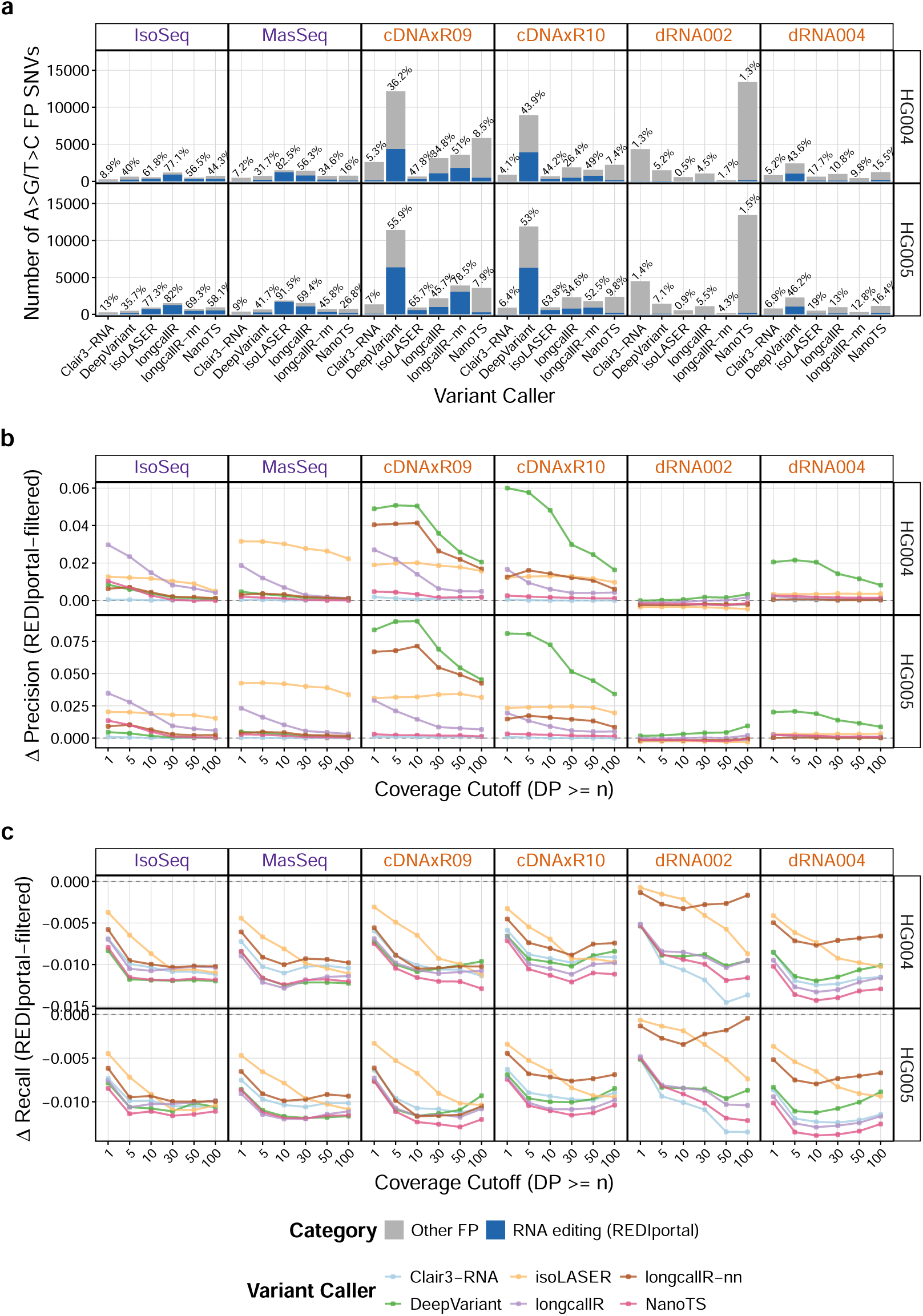
RNA editing site overlap and its impact on variant calling performance. (**a**) Number of A-to-G/T-to-C false-positive SNVs overlapping REDIportal (blue) or not (grey). Percentages of REDIportal-overlapping SNVs are shown above each bar. (**b, c**) Change in precision (**b**) and recall (**c**) after excluding REDIportal-overlapping SNVs.

Overall, RNA editing contributes measurably to the FP burden of several lrRNA-seq-specific callers; known editing sites can be filtered post hoc using databases such as REDIportal, although at the cost of removing some true variants that coincide with catalogued editing sites.

### Genomic context impacts variant-calling performance

Rather than restricting the analysis to high-confidence regions, we examined how challenging genomic features, including homopolymers, tandem repeats and GC content, influence variant calling performance. We approached this by evaluating variant calls within the genomic categories defined by the stratification from GIAB (Fig. 5a).

**Fig. 5:**
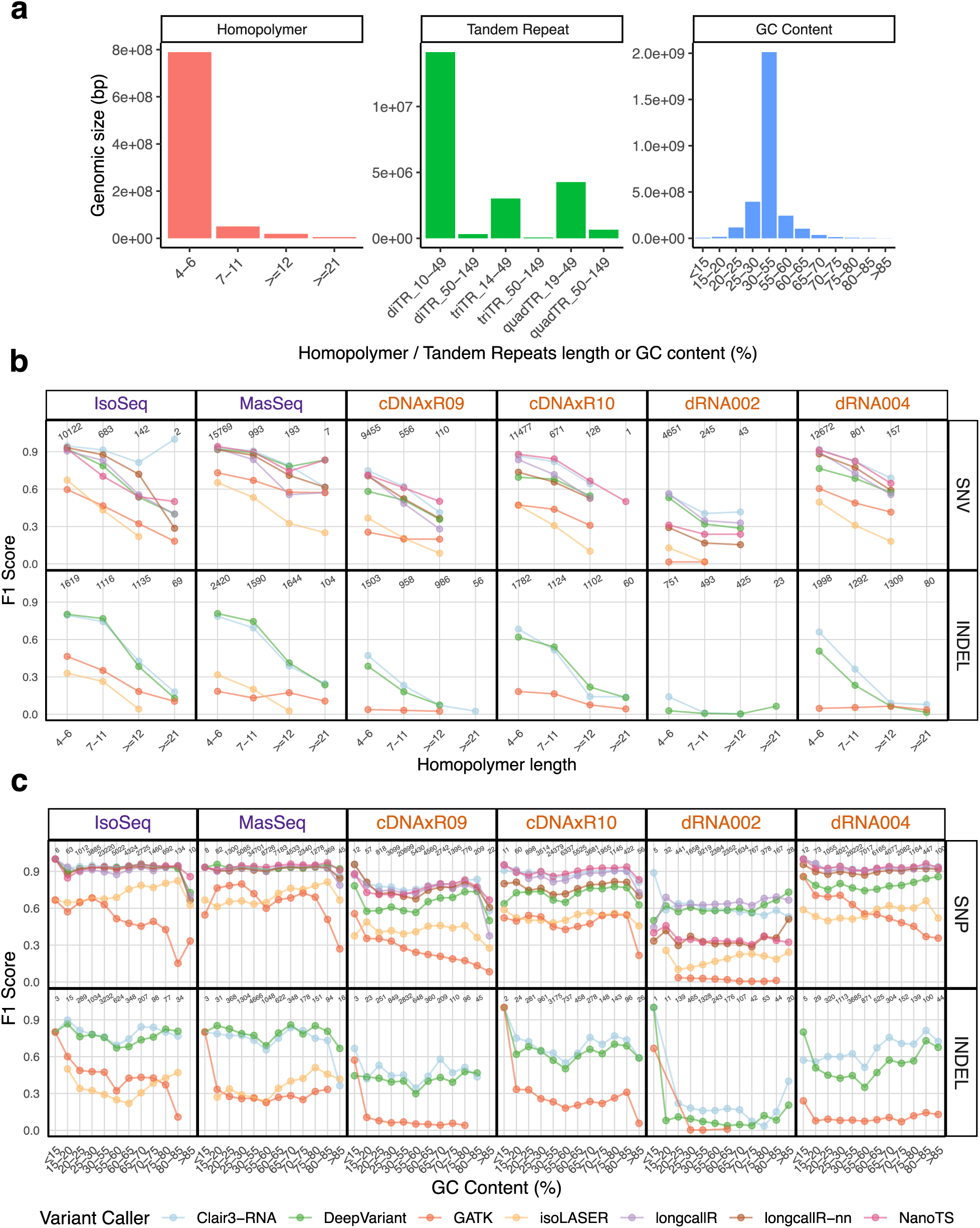
Variant calling performance across genome-context–stratified regions for HG004 datasets. (**a**) Genomic sizes in bp for three genome-context-stratified regions: homopolymer lengths, short tandem repeat subclasses and GC content (%). (**b, c**) SNV and INDEL calling performance (F1 scores) across homopolymer lengths (**b**) and GC content (**c**). GC content was computed in 100-bp sliding windows, with an additional 50-bp flanking extension applied to each region. Results are stratified by cell lines (rows) and library preparation kits (columns) and colored by variant callers. All variant calls are based on minimap2 alignment. Panel **b** and **c** use the same color legend.

Homopolymers are consecutive repeats of the same nucleotide. Such regions often cause sequencing errors, hinder alignment and complicate variant representation^[31]^. In line with this, F1 scores of all variant callers decreased with increasing homopolymer length in most cases (Fig. 5b, Fig. S19). In a small number of cases, an apparent increase in F1 score was observed at homopolymers *≥* 21 bp; however, this is inflated by the low number of true variants in these bins. Such homopolymer-associated performance degradation seems to be more pronounced for indel calling than for SNV calling and more evident in Iso-Seq datasets than in other library preparation kits. This degradation in indel calling was predominantly driven by a loss of recall, indicating the increasing difficulty of placing indels within long homopolymers, whereas the smaller decline observed for SNVs was driven more by a loss of precision (Fig. S20, Fig. S21). In addition, performance declined more rapidly with increasing length in A and T homopolymers (Fig. S22) compared with G and C homopolymers (Fig. S23). This trend could reflect several factors. One possibility is internal priming artifacts, which are enriched in poly(A) regions (or poly(T) regions on the reverse strand) in RNA-seq libraries and may introduce local alignment errors. In addition, the higher prevalence of A*/*T homopolymers in the human genome could increase their overall contribution to performance degradation^[31]^. However, these potential mechanisms were not directly evaluated in this study.

Tandem repeats are repeated regions, but composed of multi-nucleotide motifs. Here, we examined short tandem repeats (STRs), including di-, tri-, and quad-nucleotide repeats across different lengths. Overall performance was reduced compared with genome-wide regions; however, we observed only weak dependence on STR subclass (Fig. S24). The few pronounced decreases or increases occurred in bins with very low true variant counts, likely reflecting their limited genomic space (Fig. 5a) rather than a systematic STR-specific effect.

GC content introduces systematic coverage bias in short-read sequencing technologies by affecting fragmentation and PCR amplification efficiency^[32]^. In particular, regions with extreme GC content tend to exhibit reduced coverage and elevated error rates. Although long-read sequencing often exhibits more uniform coverage across GC content due to single-molecule sequencing, most cDNA-based protocols still involve reverse transcription or PCR amplification. The dRNA library protocol avoids both steps and is expected to exhibit the lowest GC bias. In addition, GC content may affect both transcript representation and base-calling accuracy, shifting the bias from read quantity to per-base error profiles^[33]^. Consistent with these considerations, we observed that variant calling performance declined in extremely GC-rich and GC-poor regions across all callers (Fig. 5c, Fig. S25). This effect was more pronounced for lower-performing callers than for higher-performing ones, for indels compared to SNVs, and in cDNA libraries compared to dRNA libraries. The GC-associated decline showed no consistent driver, with precision and recall contributing comparably across extreme-GC bins (Fig. S26, Fig. S27). Although extreme GC bins contained fewer true variants and may yield less stable F1 estimates, the consistent trends indicate that the decline is unlikely to be explained solely by sampling noise. Interestingly, we noted a localized performance dip in the 55–60% GC range across all callers, particularly for GATK HaplotypeCaller. A similar behavior for GATK HaplotypeCaller has been observed in whole-exome sequencing data^[34]^. As this interval does not represent a classical GC extreme, the non-monotonic pattern suggests that factors beyond GC content alone may contribute to the observed decrease. One possible explanation is an interaction between sequence composition and caller behavior rather than a strictly linear chemical bias, although the underlying mechanism was not investigated in this study. The comparatively smaller decline observed in dRNA libraries also raises the possibility that protocol-specific factors contribute to this effect.

Taken together, our results indicate that homopolymer length, particularly for A/T homopolymers, showed the strongest association with reduced lrRNA-seq variant-calling performance. STRs were also associated with reduced performance, although no consistent subclass-specific patterns were identified. While GC-associated effects were comparatively mild in long-read sequencing technologies, reduced performance was still observed in GC-extreme regions and unexpectedly in the 55–60% GC interval.

### Phasing performance depends on variant caller and phasing method choice

We next evaluated several long-read phasing tools (Table 2) on HG002 and HG005 datasets, the only datasets with available phasing ground truth. In addition, we included the phased output produced directly by longcallR and isoLASER, which performs joint variant calling and phasing. We aimed to identify the combinations of variant caller and phasing approach that achieved high accuracy, coverage and contiguity. Phasing accuracy was assessed using the switch error rate, where lower values indicate more accurate local phasing between consecutive heterozygous variants. Phasing coverage was measured by the number of assessed variant pairs, representing variants that are both phased and present in the ground truth, and the fraction of heterozygous variants phased. Phasing contiguity was assessed using the phase-block N50 (see Methods). Because switch error is calculated only on assessed variant pairs, methods that phase fewer variants may achieve lower switch error by restricting phasing to higher-confidence variants. Therefore, these two metrics should be considered jointly when evaluating phasing performance.

We first evaluated phasing performance using SNVs only. Phasing coverage largely reflected the number of heterozygous variants detected by each caller, with NanoTS-based phasing covering the largest number of variants in most datasets. In PacBio datasets, HiPhase generally achieved the lowest switch error rates but phased a smaller fraction of heterozygous variants and produced shorter phase blocks, suggesting a more conservative strategy. HapCUT2 provided the best balance between accuracy and coverage, whereas WhatsHap phased more variants into longer blocks at the cost of higher switch error rates. In ONT datasets, LongPhase achieved the lowest switch error rates for most caller–dataset combinations, with HapCUT2 occasionally lowest. WhatsHap again phased the largest fraction of variants. Phasing based on longcallR-nn calls generally achieved lower switch error rates than Clair3-RNA, particularly on dRNA004, although longcallR-nn phased very few variants on dRNA002. On dRNA002, NanoTS phased the most variants but also had the highest switch error rates (0.6–4.7%), consistent with its lower variant-calling precision on this chemistry. LongcallR’s native output achieved similarly low switch error rates and phased a comparable absolute number of variants to Clair3-RNA, despite covering a smaller fraction of heterozygous variants because it identified more heterozygous variants overall (Fig. 6a, c).

**Fig. 6:**
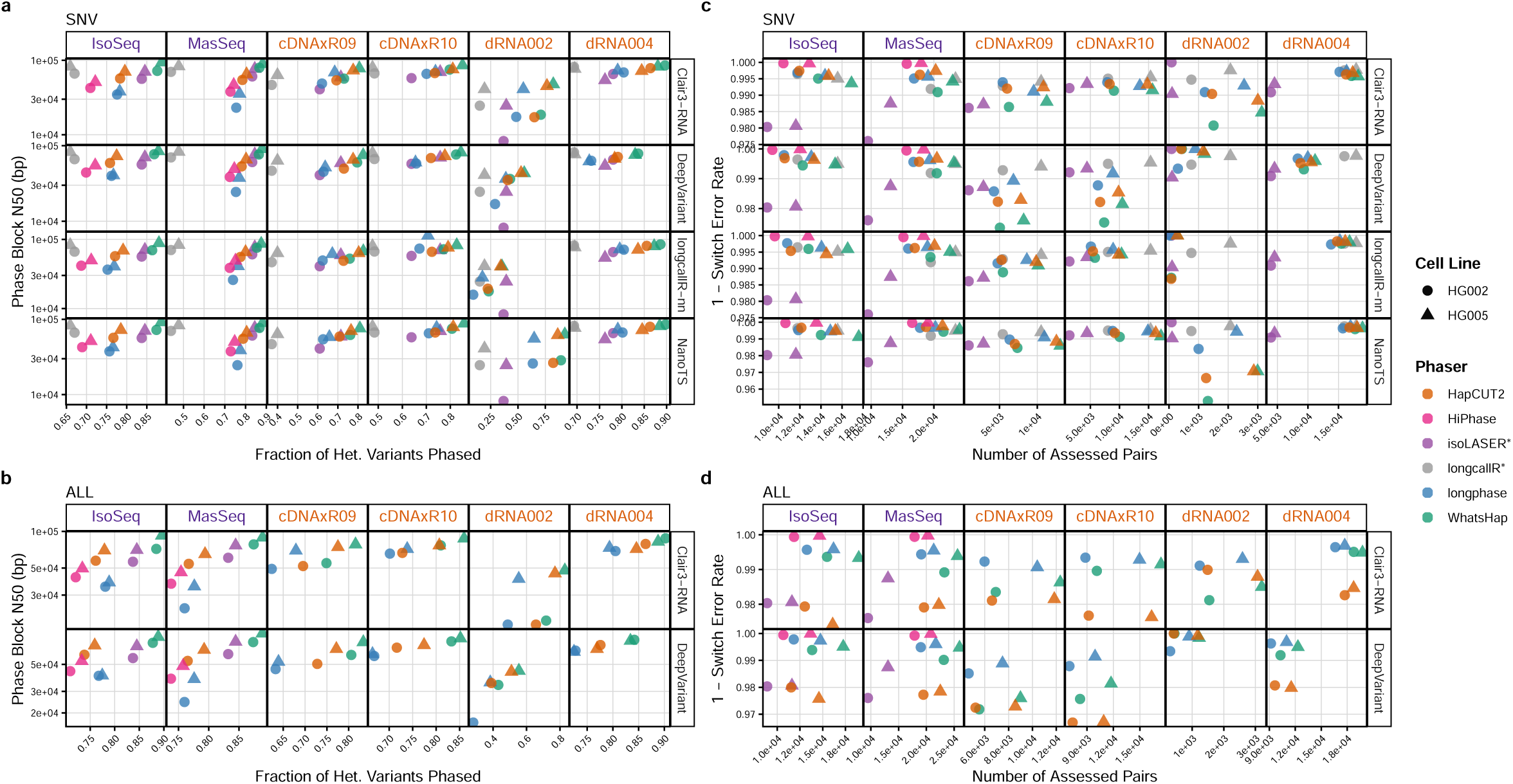
Phasing performance for SNVs and all variants (SNVs and indels), based on the output of different variant callers. Scatter plots showing phasing performance stratified by variant calling tool (rows) and library kit (columns). (**a,b**) The fraction of heterozygous variants phased and phase-block N50, for SNVs and all variants, respectively. (**c,d**) The switch error rate and number of assessed variant pairs, for SNVs and all variants, respectively. Points are shaped by cell line and colored by phasing tool. The number of assessed pairs represents heterozygous variant pairs that are both phased and present in the ground truth. *: these tools jointly call and phase variants. NanoTS provides both unphased and phased outputs, with the latter generated using WhatsHap. Here, we used its unphased VCF as input to different phasing tools.

When phasing both SNVs and indels, we observed similar trends for phasing coverage and contiguity. WhatsHap phased the largest fraction of variants and produced the longest phase blocks, whereas LongPhase yielded the lowest phase-block N50 on PacBio data. Including indels, however, markedly increased the switch error rate of HapCUT2 (1.5–3.3% vs. 0.3–1.8% for SNVs only, excluding dRNA002), making it the least accurate phaser in most datasets. In contrast, switch error rates changed only slightly for the other phasers. HiPhase remained the most accurate on PacBio data (0.01–0.08%), while LongPhase achieved the lowest switch error rates on most ONT datasets (0.3–1.5%) (Fig. 6b, d).

Overall, no single phasing method performed best across all metrics and variant types. HiPhase and LongPhase achieved the lowest switch error rates on PacBio and ONT data, respectively, whereas WhatsHap provided the highest phasing coverage and contiguity. HapCUT2 offered a good balance for SNVs but became error-prone when indels were included. Among variant callers, NanoTS-based phasing covered the most SNVs on most chemistries, while Clair3-RNA provided a robust basis for both SNV and indel phasing. When joint variant calling and phasing is preferred, longcallR offers a strong alternative.

### Variant-calling performance on LongBench datasets

After evaluating the major factors influencing variant calling performance on gold-standard GIAB datasets, we next assessed performance on the more recent LongBench datasets^[22]^. This is particularly important for methods relying on pre-trained models, which are often trained on GIAB data and might therefore exhibit limited generalization. Here, we focused on two small-cell lung cancer cell lines, H211 and H526, sequenced using PacBio Kinnex (Mas-Seq), ONT dRNA004 and cDNA R10 libraries. Similar to GIAB data, PacBio Kinnex gave higher mapped reads, longer average reads and lower sequencing error than ONT datasets (Fig. 7a). Lacking gold-standard ground truth, we generated a high-confidence reference variant set from matched Illumina WGS datasets using the GATK Best Practices pipeline, which served as the benchmark reference (see Methods).

**Fig. 7:**
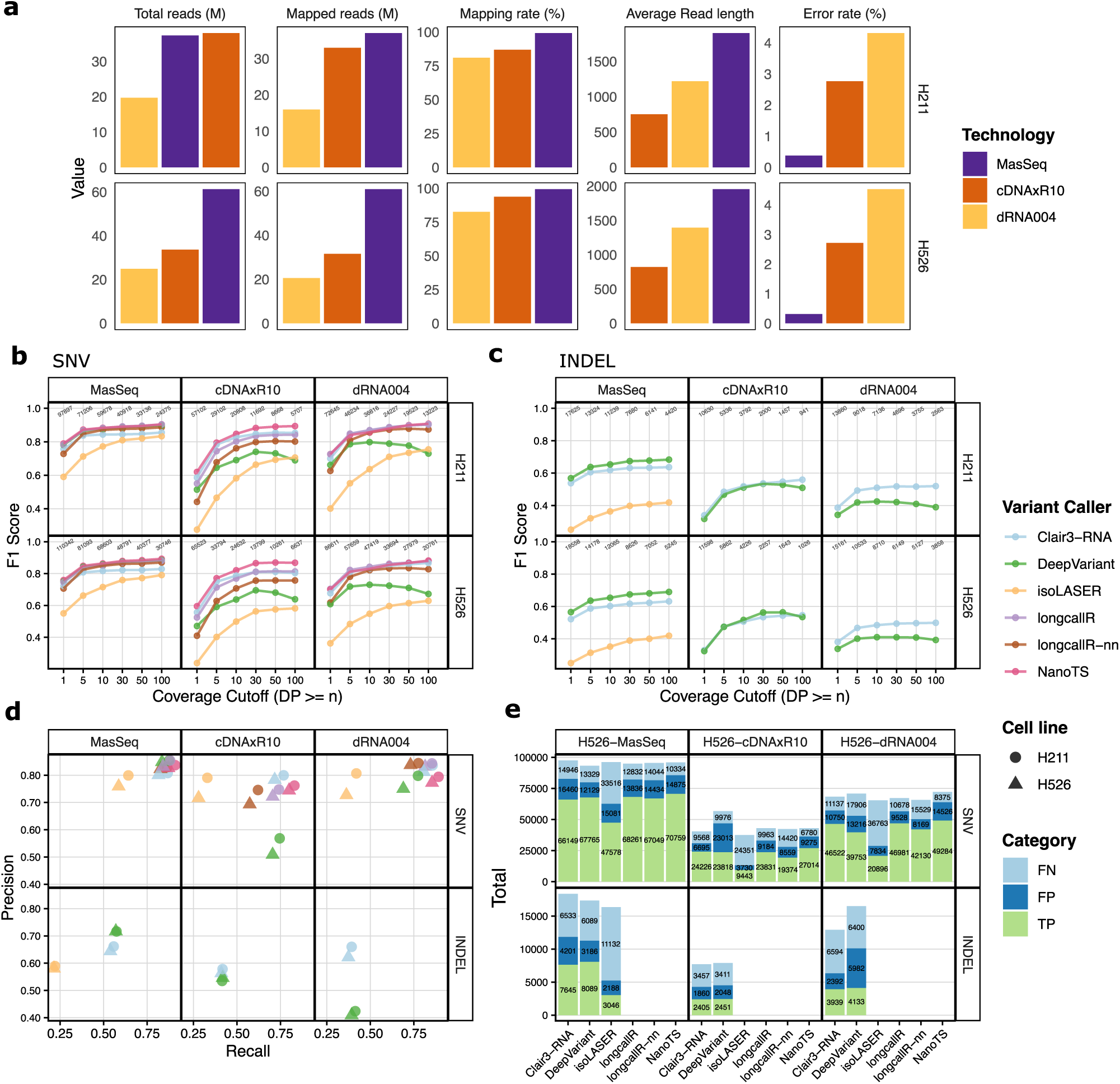
Variant calling performance on LongBench datasets. (**a**) Bar plots showing data quality, colored by library preparation kit and stratified by quality metric and cell line. Total sequences and mapped reads are shown in units of millions. Sequencing error rate refers to the ratio between mismatches and bases mapped (cigar). (**b, c**) F1 scores for calling SNVs (**b**) and indels (**c**) across different coverage, stratified by cell lines (H211 and H526) and library preparation kits. The number on top of each point indicates the number of variants in the truth set. All variant calls were based on minimap2 alignments and without BAM transformation. (**d**) Precision and recall performance for each variant caller at 5*×* coverage, stratified by variant type (SNVs and indels) and library preparation kit. (**e**) Number of TP, FN, and FP calls for each variant caller in the H526 datasets. Results for the H211 cell line are shown in Fig. S30.

First, evaluation on the LongBench datasets confirmed that sequencing characteristics drive variant calling performance (Table S11), with error rate limiting indel calling, while sequencing depth and the number of callable bases shape SNV calling (Fig. S28, Table S12, Table S13). On average, Mas-Seq achieved a low error rate of 0.34%, compared with 2.74% for cDNA R10 and 4.42% for dRNA004, and yielded the highest accuracy, with mean SNV-calling F1 scores across all callers of 0.819 at 5× coverage and 0.869 at 100× coverage (Fig. 7b-d). Between the two ONT chemistries, however, dRNA004 achieved higher SNV F1 scores than cDNA R10 (0.759 vs. 0.664 at 5×; 0.819 vs. 0.771 at 100×) despite its higher error rate. Consistent with our subsampling analysis, deeper sequencing can compensate for sequencing errors in SNV calling: owing to its longer reads, a larger fraction of the dRNA004 callable region was covered at high depth (41-47% vs. 24-25% of callable bases at 30×). In contrast, indel calling followed the error rate, with mean F1 scores of 0.517, 0.475, and 0.445 for Mas-Seq, cDNA R10, and dRNA004, respectively, at 5× coverage. Deeper coverage had little effect on indel calling for dRNA004 (0.445 at 5× vs. 0.450 at 100×), again supporting that systematic indel errors cannot be overcome by increased sequencing depth. Finally, the number of callable bases determined the number of variants that could be detected (Fig. S28b).

Mas-Seq provided, on average, 41.8 and 19.4 million more callable bases than cDNA R10 and dRNA004, respectively (Fig. S29), resulting in 45,928 (*∼*2.8-fold) and 24,684 (*∼*1.5-fold) more variants detected by Clair3-RNA at 5× coverage (Fig. 7e, Fig. S30). Second, the relative performance of the variant callers observed in the GIAB benchmark generalized to the LongBench datasets. NanoTS achieved the highest SNV accuracy overall (mean F1 = 0.837) and was the best SNV caller in most samples and coverage cutoffs, particularly on cDNA R10 (F1 = 0.783 at 5×). Clair3-RNA was strong in both SNV and indel calling, with mean F1 scores of 0.807 and 0.526, respectively. LongcallR was a competitive alternative in SNV calling, as well as longcallR-nn, especially for dRNA004, where it performed best at 5–50× coverage in both cell lines (mean F1 = 0.836 at 5×), followed closely by Clair3-RNA and NanoTS (0.828 and 0.826). In Mas-Seq, NanoTS, DeepVariant and longcallR performed comparably in SNV calling (F1 = 0.861, 0.854 and 0.852 at 5×), and DeepVariant was the best-performing caller for indel calling (mean F1 = 0.637) at 5× coverage, ahead of Clair3-RNA (F1 = 0.596; Fig. 7b, c, Table S14, Table S15). Consistent with the GIAB benchmark, DeepVariant again produced too many false positives in the ONT datasets, whereas isoLASER remained overly conservative (Fig. 7d). We again showed that the relative performance of the callers was robust to differences in allele-fraction and quality-score thresholds (Fig. S32) and to isoLASER preprocessing (Fig. S31).

Together, the LongBench analyses reinforce the dominant role of sequencing characteristics, with Mas-Seq yielding better variant calling results, and support the relative performance of the variant callers observed in the GIAB benchmark. They also provide complementary evidence from biologically relevant cancer transcriptomes.

### Computational cost

Finally, we report runtime, CPU time, and peak memory usage (maximum proportional set size, PSS) (see Methods). These metrics were presented as absolute totals, normalized per mapped gigabase, and normalized per million mapped reads. As expected, absolute runtime and CPU time were higher for PacBio Mas-Seq than for the ONT datasets for most methods, because of its high read depth and substantially longer reads (Fig. 8a); the exception was isoLASER, whose longest runs were on dRNA004. After normalizing by mapped gigabases, computational costs became much more consistent across sequencing chemistries (Fig. 8b), indicating that runtime scales primarily with the total number of mapped bases rather than the number of reads. By contrast, normalization by million mapped reads did not reduce these differences to the same extent (Fig. S33). The exceptions were isoLASER and NanoTS, which remained more expensive per gigabase on dRNA004 data than on the other chemistries (markedly so for isoLASER), and longcallR, which was approximately three times more expensive per gigabase on Mas-Seq than on the ONT datasets. isoLASER and NanoTS were also the most computationally demanding methods overall, with isoLASER requiring substantial runtime and memory, while NanoTS showed high runtime but comparatively moderate memory usage. Among the remaining methods, longcallR and longcallR-nn achieved the shortest runtimes on the ONT datasets, whereas DeepVariant was fastest on Mas-Seq; both longcallR methods, however, required substantially more memory, with longcallR exhibiting the highest memory usage. Clair3-RNA and DeepVariant provided a favorable balance between runtime and memory consumption, combining competitive runtimes with the lowest memory requirements.

**Fig. 8:**
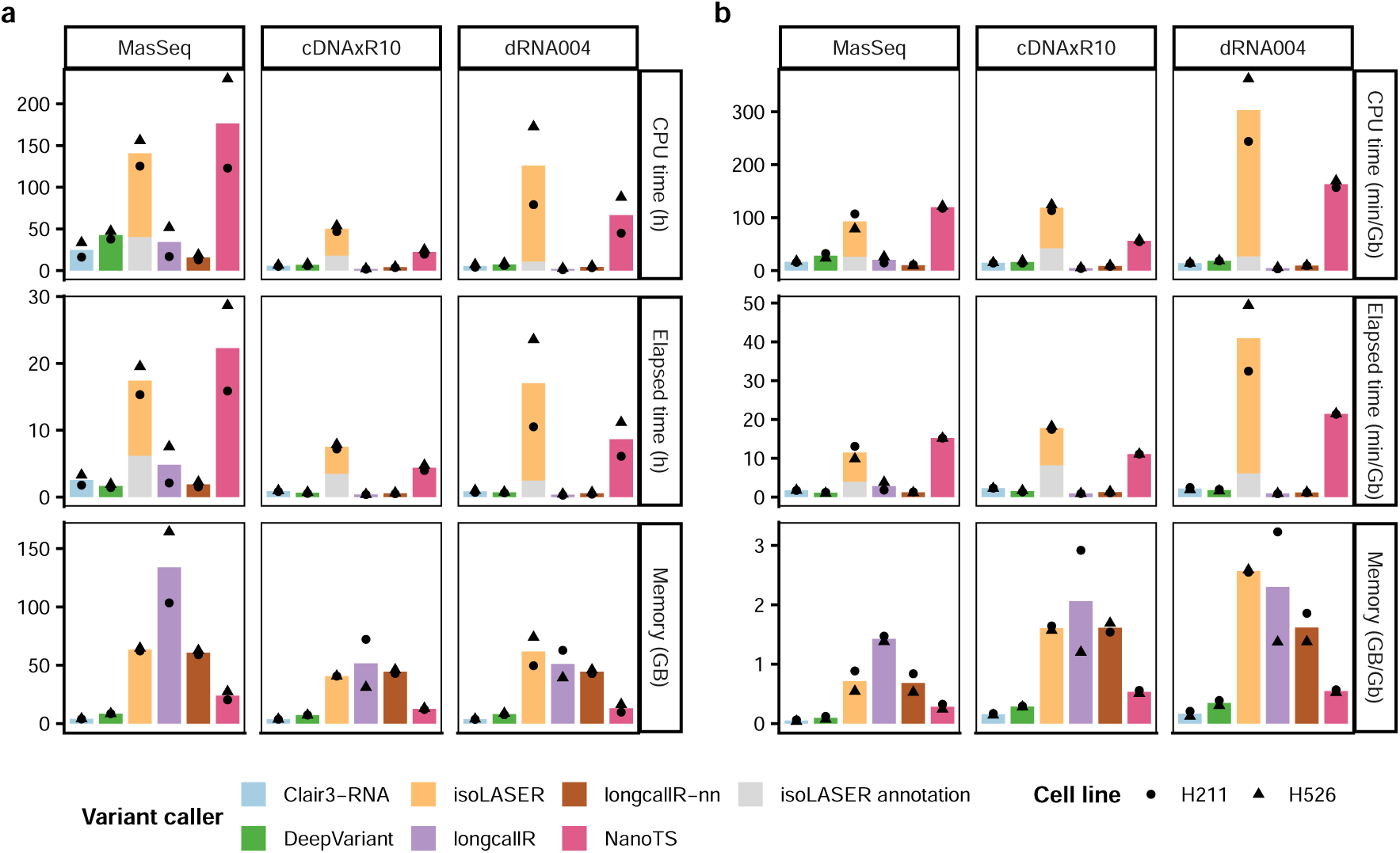
Computational costs of variant calling methods on LongBench datasets. (**a**) Absolute runtime, CPU time, and peak memory usage (maximum proportional set size, PSS) using 10 CPU cores, stratified by metric (rows) and library kit (columns). Mas-Seq was sequenced on the PacBio platform, whereas cDNAxR10 and dRNA004 were sequenced on the ONT platform. (**b**) Runtime, CPU time, and peak memory usage normalized per mapped gigabase (Gb). For isoLASER, variant calling comprises two sequential steps: an initial realignment and annotation step followed by variant calling (isoLASER) (see Methods). Absolute runtime includes both steps, whereas peak memory usage and normalized metrics are reported for the variant-calling step only. Mas-Seq is from PacBio; cDNAxR10 and dRNA004 are from ONT.

## Discussion

This study provides the first systematic and neutral comparison of emerging lrRNA-seq variant calling tools and their key performance determinants, together with an evaluation of phasing methods. Our results show that sequencing characteristics are the strongest determinant of performance, followed by the choice of variant caller, aligner, and BAM file transformation, with additional effects from genomic context. Using gold-standard GIAB datasets, variant calling with PacBio technologies achieved better overall performance than ONT, due to higher sequencing accuracy and a larger number of callable bases. Overall, all lrRNA-seq-specific callers performed reasonably well. Clair3-RNA consistently ranked among the top-performing callers across all chemistries, whereas NanoTS achieved the highest SNV performance across most chemistries matching its trained models. DeepVariant showed strong performance on PacBio datasets, especially for indels.

Beyond sequencing characteristics and variant caller choice, we find that even subtle differences in alignment can increase false calls. Although the PacBio-optimized aligner pbmm2 shares the same underlying algorithm as minimap2, differences in parameter settings led to more FPs around splice regions, consistent with observations from the Clair3-RNA developers. In addition, BAM transformation does not provide significant improvements for lrRNA-seq-specific variant callers, although it may slightly improve the performance of DeepVariant. Furthermore, lrRNA-seq remains prone to errors in challenging and repetitive genomic contexts, such as homopolymers and extreme GC regions, while RNA editing represents an additional source of false positives for some callers. Known RNA editing sites can, however, be annotated and filtered post hoc using databases such as REDIportal, albeit at a small risk of removing true variants. Phasing performance largely reflected the quality of the input variants: callers detecting more heterozygous variants generally enabled higher phasing coverage, while higher variant-calling precision was associated with lower switch error rates. We also observed a trade-off between phasing coverage and accuracy, with WhatsHap favoring coverage and contiguity, and HiPhase and LongPhase achieving the highest accuracy on PacBio and ONT data, respectively. HapCUT2 offered a good balance for SNVs, but its accuracy dropped markedly when indels were included, highlighting the challenges of indel-based phasing in lrRNA-seq data. The choice of phasing tool should therefore depend on the downstream application: accuracy-critical analyses, such as haplotype-resolved allele-specific expression, may benefit from HiPhase or LongPhase, whereas WhatsHap may be preferable when maximizing the number of phased variants is the priority. Notably, phase blocks derived from lrRNA-seq data are inherently confined to expressed genes, limiting long-range phasing compared with DNA-based approaches.

Using the more recent LongBench datasets, our results further support sequencing characteristics as a key determinant of variant-calling performance, with Mas-Seq showing the best overall performance, consistent with our GIAB-based analyses. The results also support the relative performance of variant callers observed in the GIAB benchmark, which was robust to caller-specific preprocessing and allele fractions, although users interested in low-allele-fraction variants should consider caller-specific performance in this range.

Our findings establish a foundation for evaluating variant-calling methods while highlighting several open questions for future research. First, we did not investigate how haplotype phasing improves variant calling. Among the variant callers we compared, longcallR and isoLASER are haplotype-aware, and Clair3-RNA provides an option to incorporate phasing. Future work could assess how incorporating phasing affects the performance of individual tools, for example by enabling phasing in Clair3-RNA and NanoTS or comparing end-to-end pipelines such as PEPPER-Margin-DeepVariant^[35]^. Second, our benchmarking was designed to evaluate variant-calling performance under realistic sequencing conditions. Future studies could perform more controlled comparisons by restricting analyses to a common set of highly expressed loci and sub-sampling libraries to matched read depths, thereby isolating the effects of sequencing accuracy and platform-specific sequencing errors from differences in transcript abundance and coverage. Third, while lrRNA-seq enables variant-to-isoform analyses, the present benchmark focused on variant detection accuracy. Benchmarking transcript-level variant assignment remains an important direction for future work. Finally, we did not evaluate variant-calling performance in long-read single-cell RNA-seq data. Long reads offer additional advantages for variant calling in single-cell data, as short-read scRNA-seq is typically constrained by 5’ or 3’ biases. It has been suggested to perform variant calling at the pseudobulk level^[36]^, and our findings are therefore likely broadly applicable. However, data sparsity and, more generally, shorter read lengths^[37][22][38]^ in single-cell modalities may introduce additional challenges.

More broadly, beyond the methodological considerations discussed above, variant calling in lrRNA-seq remains fundamentally constrained by gene expression, as variants can only be detected in expressed transcripts. This intrinsic limitation restricts sensitivity, particularly for lowly expressed genes or rare cell populations. Recent studies^[39][40]^ have proposed utilizing targeted lrRNA-seq approaches to mitigate this limitation by enriching for regions of interest, thereby increasing effective coverage and improving variant detection. Integrating such targeted strategies with robust variant-calling and phasing frameworks may represent a promising direction for improving sensitivity and enabling more comprehensive characterization of mutational landscapes in lrRNA-seq data.

## Conclusion

Our benchmark results show that sequencing characteristics are the strongest determinant of lrRNA-seq variant-calling performance, followed by variant caller, alignment strategies, and genomic context. Across GIAB and LongBench datasets, Clair3-RNA emerged as the most versatile tool, offering models across multiple sequencing chemistries and robust performance for both SNV and indel calling. NanoTS performed best or near-best for SNV detection on newer sequencing chemistries. DeepVariant is also a strong choice for PacBio data, particularly for indel calling. LongcallR is competitive for SNV calling on ONT data and additionally provides direct phased output. For downstream haplotype phasing, HiPhase and LongPhase achieved the lowest switch error rates on PacBio and ONT data, respectively, whereas WhatsHap maximized phasing coverage and contiguity; HapCUT2 offered a good balance for SNVs only.

## Methods

### Genome alignment

#### minimap2

Both ONT and PacBio reads in FASTQ format were mapped to the human reference genome GRCh38 using minimap2 v2.28^[23]^ with the parameters -ax splice:hq -uf for PacBio, -ax splice -uf -k14 for ONT dRNA, and -ax splice for ONT cDNA. Splice junctions from the GENCODE v47 annotation were provided via the --junc-bed argument.

#### pbmm2

PacBio reads were mapped to the same reference genome using pbmm2 (v26.2.99, https://github.com/PacificBiosciences/pbmm2), a wrapper of minimap2 optimized for PacBio data, with the --preset ISOSEQ option.

### BAM transformation

For each genome-aligned BAM file, we used *GATK* v.4.6.1 *SplitNCigarReads*^[26]^ to split reads at exon-intron junctions, thereby dividing each long read into multiple exon-specific segments. Subsequently, we used *flagCorrection*^[13]^ to ensure that every exon was labeled with primary alignment flag.

### Variant calling methods

All variant callers were provided with genome-aligned BAM files and the reference genome FASTA file (GRCh38.p14) as input, except for isoLASER, which requires re-alignment of genome-aligned reads to the transcriptome. Clair3-RNA is a deep learning–based variant caller that integrates pileup- and full-alignment–based representations for variant detection. DeepVariant represents aligned sequencing reads as image-like pileups and applies a deep convolutional neural network (CNN) to classify genomic loci as true variants or sequencing errors; although originally developed for DNA sequencing, it has been extended to PacBio lrRNA-seq by training a dedicated MASSEQ model. NanoTS combines convolutional and fully connected layers to predict SNV genotypes from read-level features, including base composition, strand-specific allele coverage and distance to read ends, and optionally re-genotypes variants from haplotype-resolved reads obtained with WhatsHap. IsoLASER identifies small variants through local re-assembly of reads into a de Bruijn graph and uses a multi-layer perceptron classifier to filter false-positive calls. LongcallR-nn also applies a CNN to pileup images to predict SNV genotypes and zygosity, while longcallR uses a probabilistic model to jointly refine SNV calls and perform haplotype phasing from lrRNA-seq data. Variant calling was performed using DeepVariant v1.9.0^[10]^, isoLASER v0.03^[15]^, Clair3-RNA v0.2.2^[14]^, NanoTS v1.0.0^[16]^, longcallR v1.12.0^[17]^, longcallR-nn v0.0.2^[17]^ and GATK Haplotype-Caller v.4.6.1^[26]^. Variants were annotated using snpEff v5.2c^[41]^.

#### Clair3-RNA

The --platform parameter was set according to the sequencing protocol and aligner. For PacBio Mas-Seq and Iso-Seq data, hifi mas pbmm2/minimap2 and hifi sequel2 pbmm2/minimap2 (depends on the aligner) were used, respectively. For ONT dRNA002/004 data, ont dorado drna002/drna004 was used depending on the library kit, and for ONT cDNA R09/R10 data, ont r9 guppy cdna/ont r10 dorado cdna was applied. RNA editing sites were tagged with the argument --tag variant using readiportal, and --min coverage was set to 0 (see Table 3).

**Table 3:**
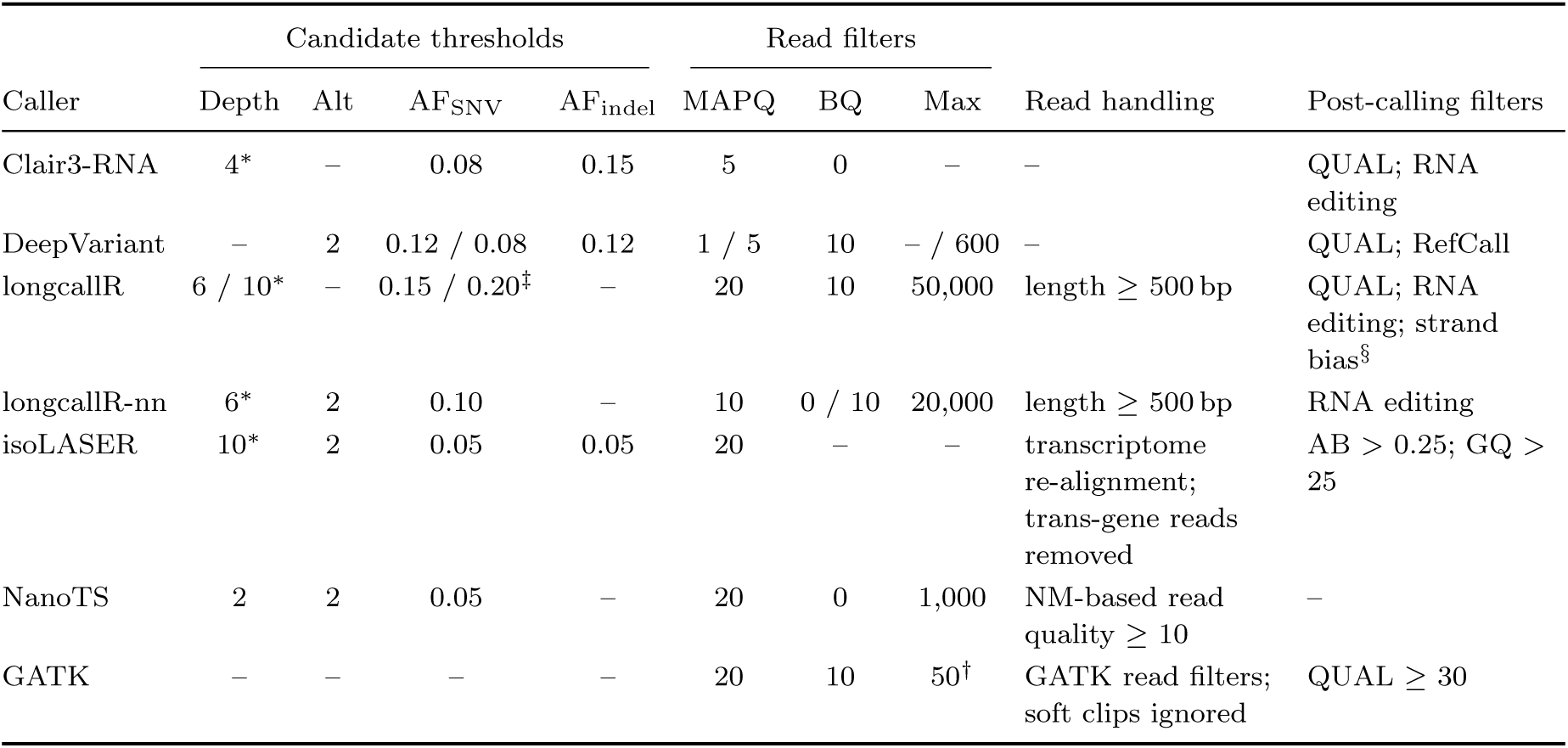
Candidate-generation and filtering parameters of each variant caller. Values shown are the default parameters of each caller, which were used in this study unless marked *^∗^*. Depth: minimum read depth; Alt: minimum number of reads supporting the variant; AF: minimum allele fraction; BQ: minimum base quality; Max: maximum number of reads per site. Values are given as PacBio / ONT where they differ; –: no such criterion. *^∗^*Set to 0 in this study, so no depth limit was applied during variant calling. Depth was instead restricted uniformly across all callers during the coverage-cutoff evaluation. NanoTS was kept at its default value of 2, which is equivalent to 0 because it already requires two supporting alternative reads. Minimum numbers of supporting reads were left unchanged. *^†^*At most 50 reads starting at the same position are kept. *^‡^*Or *≥* 0.05 with *≥* 10 supporting reads. *^§^*Iso-Seq and ONT cDNA presets only. For benchmarking, only calls with FILTER = PASS were retained for all callers (GATK: QUAL *≥* 30).

|  | Candidate thresholds |  |  |  | Read filters |  |  |  |  |
|---|---|---|---|---|---|---|---|---|---|
| Caller | Depth | Alt | AF<sub>SNV</sub> | AF<sub>indel</sub> | MAPQ | BQ | Max | Read handling | Post-calling filters |
| Clair3-RNA | 4* | – | 0.08 | 0.15 | 5 | 0 | – | – | QUAL; RNA editing |
| DeepVariant | – | 2 | 0.12 / 0.08 | 0.12 | 1 / 5 | 10 | – / 600 | – | QUAL; RefCall |
| longcallR | 6 / 10* | – | 0.15 / 0.20<sup>‡</sup> | – | 20 | 10 | 50,000 | length ≥ 500 bp | QUAL; RNA editing; strand bias<sup>§</sup> |
| longcallR-nn | 6* | 2 | 0.10 | – | 10 | 0 / 10 | 20,000 | length ≥ 500 bp | RNA editing |
| isoLASER | 10* | 2 | 0.05 | 0.05 | 20 | – | – | transcriptome re-alignment; trans-gene reads removed | AB > 0.25; GQ > 25 |
| NanoTS | 2 | 2 | 0.05 | – | 20 | 0 | 1,000 | NM-based read quality ≥ 10 | – |
| GATK | – | – | – | – | 20 | 10 | 50<sup>†</sup> | GATK read filters; soft clips ignored | QUAL ≥ 30 |

#### DeepVariant

DeepVariant was run using run deepvariant with --num shards 10. For PacBio Iso-Seq and Mas-Seq data, we used --model type MASSEQ; for ONT data, we used --model type ONT R104, as no ONT RNA-seq model is available. All other parameters were left at their defaults.

#### longcallR

The argument --preset depends on the platform and library kit: hifi-masseq for PacBio Mas-Seq/Iso-Seq data, and ont-cdna/drna for ONT cDNA or dRNA data. The minimum depth for variant filtering (--minimum-depth) was set to 0 (default: 10) to allow manual coverage cutoff in subsequent analyses.

#### longcallR-nn

Variant calling with longcallR-nn includes two steps. First, we used longcallR-dp --mode predict to extract training datasets depending on the sequencing platform and library kit of the input reads, with --min-depth 0 (see Table 3). The minimum base quality (--min-baseq) was set to 0 for ONT datasets and 10 for PacBio datasets, as suggested by the developers. The second step was to call variants using longcallR nn call, with the library-kit-specific trained model. The maximum depth threshold --max depth was set to 200.

#### isoLASER

Variant calling using isoLASER consists of three steps. First, we generated a transcriptome reference from the GENCODE v47 annotation using isolaser convert gtf to fasta and extracted exons using isolaser extract exon parts. Second, reads were re-aligned to the transcriptome using minimap2 and annotated with transcript IDs using isolaser annotate. Third, isolaser was run on the annotated BAM file with the transcriptome database and the reference genome, using --platform PacBio or --platform Nanopore, --DP 0 (see Table 3) and --include-continuous. Following the authors’ recommendations, PASS calls were further filtered for allelic balance (AB *>* 0.25) and genotype quality (GQ *>* 25), and at multi-allelic sites only the two alleles with the highest quality were retained.

#### NanoTS

NanoTS was run with nanoTS full pipeline with default parameters (--ALT 2 --total 2 --ratio 0.05 --depth 1000 --hap qual 10, set explicitly). Pretrained models are available only for the most recent sequencing chemistries (ONT cDNA R10.4.1, ONT dRNA004 and PacBio MAS-Seq). We therefore applied the ONT cDNA model (cDNA nanopore R104 HG002) to all ONT cDNA datasets, including R9.4.1, the ONT dRNA model (dRNA nanopore R104 HG002) to all ONT dRNA datasets, including dRNA002, and the MAS-Seq model (MASseq PacBio Revio HG002) to both MAS-Seq and Iso-Seq datasets. We evaluated the unphased variant calls (unphased predict.pass.vcf).

#### GATK HaplotypeCaller

The --dont-use-soft-clipped-bases argument was set to true, and --pcr-INDEL-model was set to none for ONT dRNA reads and conservative for ONT cDNA and PacBio reads.

### Phasing methods

All phasing tools are read-based and use minimap2-aligned BAM files, a reference genome (FASTA), and filtered heterozygous PASS variants (VCF) as input.

#### WhatsHap

Each VCF file (at the sample level) was phased using WhatsHap v2.8^[18]^, with the reference genome provided as input and the --ignore-read-groups argument specified.

#### HapCUT2

Phasing with HapCUT2 v1.2 was performed in two steps. First, read fragments supporting each variant were extracted from the alignment BAM files using *extractHAIRS*, with the corresponding reference genome and filtered VCF as input, and the --pacbio or --ont argument specifying the sequencing platform. Second, the resulting fragment files were provided to HapCUT2^[19]^ together with the same VCF to assemble haplotypes and generate phased variant calls.

#### LongPhase

LongPhase v1.7.3^[20]^ was run with variant calls in VCF format, the reference genome, and a genome-aligned BAM file. The --INDELs option was used to enable co-phasing of SNVs and indels, and --ont or --pb was specified to indicate the sequencing platform.

#### HiPhase

In addition to standard input files, HiPhase v1.6.0^[27]^ was run with --rna, which is configured for RNA-seq (Iso-Seq) datasets, and --phase-singletons to enable phasing of singleton blocks.

### Benchmarking

#### Sequencing quality

Sequencing quality metrics (total reads, mapped reads, mapping rate, average read length, average base quality, and sequencing error rate) were computed using samtools stats (v1.21)^[42]^ on minimap2-aligned reads.

#### Variant calling performance

To ensure a reliable comparison, we first defined a benchmark region by intersecting sequencing regions that met a specified coverage cutoff with GENCODE v47 annotated exons and, for the GIAB datasets, the corresponding high-confidence regions, using bedtools v2.26.0^[43]^. Per-base sequencing depth was computed with mosdepth v0.3.3^[44]^. The number of callable bases for a given sample and coverage threshold was calculated as the summed length of all intervals in the resulting BED file. Variant calls were then compared to the ground truth using *hap.py* ^[45]^, with VCF files from both the variant caller and the truth set as input, together with a BED file defining the benchmark regions, and the option --engine=vcfeval from RTG Tools^[46]^. Genome-context–stratified comparisons were performed using *qfy .py*, with additional genome stratification BED files (v3.6) from GIAB as input. Variant calling performance was mainly evaluated using F1 score, recall and precision.

To assess the effect of variant allele fraction (VAF), we computed the observed VAF of each truth SNV from the RNA reads as the fraction of non-reference reads at that position, using samtools mpileup (-q 20 -Q 0 -B -d 2000). Truth SNVs within the benchmark region at a coverage cutoff of 5*×* were binned by VAF (0.05–0.10, 0.10–0.15, 0.15–0.20, 0.20–0.50 and *≥*0.50), and recall in each bin was calculated from the hap.py truth labels as TP/(TP+FN). Truth SNVs with VAF *<* 0.05 were excluded. To assess robustness to caller-specific calling thresholds, we further constructed precision–recall curves by sweeping each caller’s variant quality score (QUAL), using the ROC output of hap.py.

#### Credible intervals for performance metrics

To quantify uncertainty in F1 score, we computed 95% credible intervals using a Beta-Binomial model with a Jeffreys prior (*α* = *β* = 0.5), equivalent to the approach implemented by happyCompare^[45]^ (https://github.com/Illumina/happyCompare) for hap.py output. Using the identity

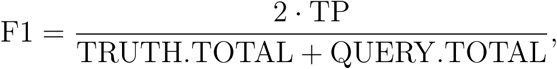

we defined *N* = TRUTH.TOTAL + QUERY.TOTAL and treated TP as the number of successes out of *N* trials, yielding the posterior

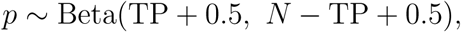

where *p* = TP*/N*. The 95% credible interval for F1 was then obtained by multiplying the 2.5th and 97.5th percentiles of this posterior by two.

#### Subsample analysis

The aligned HG004-MasSeq BAM (minimap2) was downsampled with samtools view-s seed.fraction to match the callable bases (DP*≥*5) of HG004-dRNA004, with the fraction of 0.4375 determined by binary search and applied across five random seeds (1-5) to generate independent replicates.

#### RNA editing site overlap

FP calls were defined as variants classified as BD=FP by *hap.py /vcfeval* that passed the caller’s quality filter and were not annotated as RNA editing sites by the corresponding caller. FP SNVs were intersected with REDIportal v3 (GRCh38) ^[29]^ using bcftools intersect, requiring an exact match of chromosome, position, reference, and alternate allele.

#### Phasing performance

Phasing performance was evaluated using the switch error rate (see below), the number of phased variants overlapping the truth set, the fraction of heterozygous variants phased, and the phase-block N50. Phased VCF files were compared with ground-truth phased VCFs from GIAB, derived using either trio-based or WGS-based approaches, using whatshap compare within the benchmark regions at a coverage cutoff of 5x. The phase-block N50 was calculated using whatshap stats.

In a phased VCF file, each heterozygous genotype represents the assignment of alleles to the two haplotypes. A switch error occurs when the predicted phase flips between consecutive heterozygous variants relative to the ground truth. The switch error rate is defined as the number of switch errors divided by the total number of possible phase transitions between heterozygous sites. The fraction of phased heterozygous variants was calculated as the number of phased heterozygous variants divided by the total number of heterozygous variants in the input VCF. A phase block is a set of heterozygous variants that are phased relative to one another. The phase-block N50 is the phase-block length such that 50% of the total phased sequence is contained in phase blocks of this length or longer, and measures the contiguity of phasing.

#### Ground truth VCFs for LongBench datasets

We generated a high-confidence reference variant set from the publicly available Illumina whole-genome sequencing (WGS) datasets for H211 and H526 using the GATK Best Practices germline variant calling pipeline. Paired-end reads in FASTQ format were aligned to the same GRCh38 reference genome used for the lrRNA-seq analyses using BWA-MEM2^[47]^. Only primary alignments were retained, duplicate reads were marked using GATK MarkDuplicates, and base quality score recalibration (BQSR) was performed using GATK with dbSNP build 138 and the Mills and 1000G gold standard indel resource as known sites. Variants were then called in GVCF mode using GATK HaplotypeCaller, jointly genotyped using GenotypeGVCFs, and filtered using GATK hard-filtering criteria. Only PASS variants were retained in the reference variant set for downstream benchmarking.

#### Computational cost

Clair3-RNA, DeepVariant, longcallR and isoLASER process the entire BAM file, whereas longcallR-nn operates on chromosome-wise BAM files. To ensure a fair comparison, we used 10 threads for Clair3-RNA, DeepVariant, isoLASER, longcallR. For longcallR-nn and NanoTS, we ran it per chromosome and likewise limited to 10 concurrent threads, followed by merging per-chromosome VCF files into a single VCF.

#### F1 score, Precision, and Recall

The F1 score provides a balanced measure of variant calling accuracy by combining precision and recall into a single metric, defined as the harmonic mean of the two:

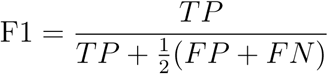

where TP, FP, and FN represent the numbers of true positives, false positives, and false negatives, respectively. Precision and recall were also computed individually as

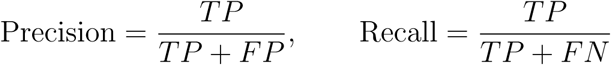

#### False discovery rate and false negative rate

The false discovery rate (FDR) and false negative rate (FNR) were calculated as

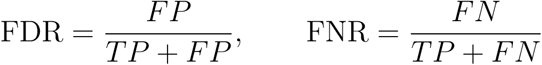

where TP, FP, and FN denote true positives, false positives, and false negatives, respectively.

## Availability of data and materials

The Iso-Seq, Mas-Seq (PacBio) and dRNA002 datasets for HG002 (NA24385), HG004 (NA24143) and HG005 (NA24631) are available at GIAB RNA-seq data repository. ONT R9 datasets are available for HG002, HG004, and HG005. The ONT dRNA004 and cDNA R10 datasets for these cell lines were generated by the developers of Clair3-RNA^[14]^. The dRNA004 datasets are available at NCBI under accession PRJNA1169852, and the cDNA R10 datasets, generated by University of Hong Kong, can be accessed at ONT cDNA R10 data repository. The ground truth variants and high confidence regions are available at GIAB ground truth and confidence region repository. The predefined genome stratification v3.6 is available at GIAB genome stratification repository. Longbench datasets are available at LongBench data repository.

The benchmark workflows and figure-generation code are available on GitHub: variant-comparison (GIAB, via Snakemake v9.20.0^[48]^) and VarCallBench (LongBench, via Omnibenchmark v0.5.1^[49]^). All code and output (raw and phased) variants are stored in VCF format and evaluation results are available on Zenodo^[50]^.

## Ethics, Consent to Participate, and Consent to Publish

Not applicable

## Authors’ contributions

J.W. conceived the study design, developed the computational workflow, and implemented all methods, with assistance from M.D.R. J.W. wrote the original manuscript, with edits by M.D.R. All authors reviewed, edited, and approved the final manuscript.

## Acknowledgments

We thank the members of the Robinson Lab at University of Zürich for their valuable feedback on this project, especially David Wissel for insightful discussions and feedback on the manuscript. We also thank the developers of Clair3-RNA, longcallR(-nn), DeepVariant and isoLASER for their help on implementing their methods.

