## Supplementary Information for "Systematic benchmarking of small variant calling pipelines for long-read RNA sequencing data"

### 1 Supplementary Tables

#### 1.1 Dataset information

| Short name | Source | Platform | Library Kit |
|---|---|---|---|
| HG002-IsoSeq | Ashkenazim Trio son | PacBio Sequel II | Iso-Seq |
| HG002-MasSeq | Ashkenazim Trio son | PacBio Revio | Mas-Seq |
| HG002-cDNAxR09 | Ashkenazim Trio son | ONT MinION | SQK-PCS111 |
| HG002-cDNAxR10 | Ashkenazim Trio son | ONT MinION | SQK-PCS114 |
| HG002-dRNA002 | Ashkenazim Trio son | ONT PromethION | SQK-RNA002 |
| HG002-dRNA004 | Ashkenazim Trio son | ONT PromethION | SQK-RNA004 |
| HG004-IsoSeq | Ashkenazim Trio mother | PacBio Sequel II | Iso-Seq |
| HG004-MasSeq | Ashkenazim Trio mother | PacBio Revio | Mas-Seq |
| HG004-cDNAxR09 | Ashkenazim Trio mother | ONT MinION | SQK-PCS111 |
| HG004-cDNAxR10 | Ashkenazim Trio mother | ONT MinION | SQK-PCS114 |
| HG004-dRNA002 | Ashkenazim Trio mother | ONT PromethION | SQK-RNA002 |
| HG004-dRNA004 | Ashkenazim Trio mother | ONT PromethION | SQK-RNA004 |
| HG005-IsoSeq | Chinese Trio son | PacBio Sequel II | Iso-Seq |
| HG005-MasSeq | Chinese Trio son | PacBio Revio | Mas-Seq |
| HG005-cDNAxR09 | Chinese Trio son | ONT MinION | SQK-PCS111 |
| HG005-cDNAxR10 | Chinese Trio son | ONT MinION | SQK-PCS114 |
| HG005-dRNA002 | Chinese Trio son | ONT PromethION | SQK-RNA002 |
| HG005-dRNA004 | Chinese Trio son | ONT PromethION | SQK-RNA004 |

**Table S1: GIAB datasets information.** HG002 cell line has three replicates: NA24385, NA26105 and NA27730, while only the canonical HG002 NA24385 was used for benchmarking. The Iso-Seq datasets analysed in this benchmark study were sequenced by Baylor College of Medicine.

| Dataset | Clair3-RNA | longcallR-nn | DeepVariant[*] | isoLASER | NanoTS[§] |
|---|---|---|---|---|---|
| HG001-IsoSeq[†] | | | | ✓ | |
| HG001-dRNA002[†] | | | | ✓ | |
| HG002-IsoSeq | ✓ | ✓ | | | |
| HG002-MasSeq | ✓ | NA24385 | ✓ | | NA24385 |
| HG002-cDNAxR09 | ✓ | | | | |
| HG002-cDNAxR10 | ✓ | | | | NA24385 |
| HG002-dRNA002 | ✓ | | | | |
| HG002-dRNA004 | NA24385 | NA24385 | | | NA24385 |
| HG004-MasSeq | | | ✓ | | |
| HG005-MasSeq | | | ✓ | | |
| WTC11-cDNAxR09[†] | | ✓ | | | |

**Table S2: Training datasets used by deep learning-based variant callers.** Sample IDs indicate which HG002 replicates were used for training; ✓ indicates all three available replicates were used. [†]these were not used in the benchmark study. [*]DeepVariant excludes chr20 from training. [§]NanoTS excludes chr20 from training but uses it as the validation set for model selection; its cDNA model was trained on an in-house HG002 dataset (SRR33022232).

#### 1.2 Correlation between data characteristics and variant calling performance

| Type | Variant Caller | 1 | 5 | 10 | 30 | 50 | 100 |
|---|---|---|---|---|---|---|---|
| SNV | Clair3-RNA | 0.86 | 0.84 | 0.85 | 0.84 | 0.83 | 0.82 |
| | DeepVariant | 0.91 | 0.85 | 0.86 | 0.87 | 0.88 | 0.82 |
| | GATK | 0.87 | 0.88 | 0.88 | 0.86 | 0.86 | 0.87 |
| | isoLASER | 0.89 | 0.88 | 0.85 | 0.83 | 0.83 | 0.81 |
| | longcallR | 0.84 | 0.85 | 0.93 | 0.94 | 0.94 | 0.94 |
| | longcallR-nn | 0.91 | 0.88 | 0.93 | 0.95 | 0.96 | 0.95 |
| | NanoTS | 0.84 | 0.91 | 0.92 | 0.89 | 0.90 | 0.82 |
| INDEL | Clair3-RNA | 0.85 | 0.80 | 0.79 | 0.77 | 0.77 | 0.76 |
| | DeepVariant | 0.79 | 0.75 | 0.74 | 0.71 | 0.71 | 0.71 |
| | GATK | 0.74 | 0.70 | 0.70 | 0.68 | 0.66 | 0.69 |

**Table S3: Spearman correlation between (1 − error rate) and F1 score.** Correlations are computed per variant caller, variant type, and coverage cutoff, across all samples, cell lines and sequencing platforms/chemistries ($n = 18$ per row). isoLASER INDEL calls are restricted to PacBio; longcallR, longcallR-nn and NanoTS do not call INDELs. Therefore, these were omitted from INDEL block.

| Type | Variant Caller | 1 | 5 | 10 | 30 | 50 | 100 |
|---|---|---|---|---|---|---|---|
| SNV | Clair3-RNA | 0.93 | 0.99 | 1.00 | 0.99 | 0.99 | 0.99 |
| | DeepVariant | 0.85 | 0.94 | 0.97 | 0.98 | 0.98 | 0.98 |
| | GATK | 0.91 | 0.95 | 0.97 | 0.97 | 0.97 | 0.94 |
| | isoLASER | 0.85 | 0.87 | 0.87 | 0.90 | 0.98 | 0.99 |
| | longcallR | 0.96 | 0.99 | 1.00 | 1.00 | 1.00 | 0.99 |
| | longcallR-nn | 0.89 | 0.95 | 0.98 | 0.99 | 0.99 | 1.00 |
| | NanoTS | 0.95 | 0.99 | 1.00 | 0.99 | 1.00 | 0.99 |
| INDEL | Clair3-RNA | 0.86 | 0.90 | 0.91 | 0.96 | 0.96 | 0.97 |
| | DeepVariant | 0.85 | 0.87 | 0.88 | 0.94 | 0.96 | 0.95 |
| | GATK | 0.88 | 0.93 | 0.94 | 0.93 | 0.95 | 0.94 |

**Table S4: Spearman correlation between callable bases and number of TPs.** Correlations are computed per variant caller, variant type, and coverage cutoff, across all samples, cell lines and sequencing platforms/chemistries ($n = 18$ per row). isoLASER INDEL calls are restricted to PacBio; longcallR, longcallR-nn and NanoTS do not call INDELs. Therefore, these were omitted from INDEL block.

To quantify the relative contribution of sequencing chemistry, variant caller, variant type, and coverage cutoff to performance, we fit linear models with F1 score and $\log_{10}$(number of true positives +1) as outcomes, using hap.py results merged with per-sample quality metrics (error rate, mapping rate, read length) and per-sample, per-coverage callable-bases counts. Each model included chemistry, caller, variant type, coverage cutoff, and cell line as additive predictors:

$$\text{F1 or } \log_{10}(\text{TP}+1) \sim \text{chemistry} + \text{caller} + \text{variant type} + \text{coverage cutoff} + \text{cell line}$$

Error rate, mapping rate, and read length were not included as separate terms, as they did not explain additional variance once chemistry was included and were represented by chemistry identity instead. GATK was set as the reference level for caller. The unique variance explained by each term (partial $R^2$) was computed by term deletion: $(RSS_{\text{reduced}} - RSS_{\text{full}})/RSS_{\text{reduced}}$. Both models were fit on $n = 1{,}008$ observations across 18 samples ($R^2 = 0.89$ and 0.90, respectively).

| Term | F1 score | $\log_{10}$(TP count) |
|---|---|---|
| Chemistry | 0.784 | 0.791 |
| Variant caller | 0.726 | 0.312 |
| Variant type (SNV/INDEL) | 0.573 | 0.766 |
| Coverage cutoff | 0.404 | 0.378 |
| Cell line | 0.023 | 0.010 |

**Table S5: Partial $\mathbf{R^2}$ of factors contributing to variant-calling performance.** Partial $R^2$ was computed by term deletion from linear models relating F1 score and $\log_{10}$(number of TPs + 1) to sequencing chemistry, variant caller, variant type, coverage cutoff, and cell line. Values indicate the unique variance explained by each factor after accounting for all others; both models achieved $R^2 = 0.90$.

| | F1 score | | $\log_{10}$(TP count) | |
|---|---|---|---|---|
| Term | Estimate | $p$ | Estimate | $p$ |
| caller: Clair3-RNA | 0.388 | <2e-16 | 0.428 | <2e-16 |
| caller: DeepVariant | 0.302 | <2e-16 | 0.357 | <2e-16 |
| caller: isoLASER | 0.093 | <2e-16 | 0.104 | 0.0002 |
| caller: longcallR | 0.357 | <2e-16 | 0.412 | <2e-16 |
| caller: longcallR-nn | 0.288 | <2e-16 | 0.312 | <2e-16 |
| caller: NanoTS | 0.347 | <2e-16 | 0.448 | <2e-16 |
| chemistry: cDNAxR10 | 0.085 | <2e-16 | 0.179 | 2.9e-11 |
| chemistry: dRNA002 | −0.289 | <2e-16 | −1.053 | <2e-16 |
| chemistry: dRNA004 | 0.099 | <2e-16 | 0.302 | <2e-16 |
| chemistry: IsoSeq | 0.220 | <2e-16 | 0.178 | 1.9e-11 |
| chemistry: MasSeq | 0.212 | <2e-16 | 0.447 | <2e-16 |
| Type: INDEL | −0.253 | <2e-16 | −1.106 | <2e-16 |

**Table S6: Linear model coefficients for F1 score and variant count.** Reference levels: variant caller = GATK, chemistry = cDNAxR09, variant type = SNV. Coefficients for coverage cutoff and cell line are omitted here for brevity (both models control for these terms; see Table S 5 for their partial $R^2$). All models: $n = 1116$ observations across 18 samples.

#### 1.3 Variant calling performance

**Table S7:** F1 scores with 95% Jeffreys credible intervals for SNV calling on GIAB HG004 and HG005 datasets.

| Dataset | Coverage | Clair3-RNA | DeepVariant | longcallR | longcallR-nn | NanoTS | isoLASER | GATK |
|---|---|---|---|---|---|---|---|---|
| HG004-IsoSeq | 1 | 0.745 [0.740, 0.751] | 0.754 [0.748, 0.759] | 0.688 [0.682, 0.693] | 0.684 [0.679, 0.690] | 0.744 [0.739, 0.750] | 0.477 [0.471, 0.482] | 0.428 [0.423, 0.434] |
| | 5 | 0.942 [0.935, 0.949] | 0.936 [0.929, 0.943] | 0.901 [0.894, 0.908] | 0.927 [0.920, 0.934] | 0.937 [0.930, 0.943] | 0.699 [0.692, 0.707] | 0.591 [0.584, 0.598] |
| | 10 | 0.960 [0.952, 0.967] | 0.954 [0.946, 0.962] | 0.939 [0.931, 0.947] | 0.951 [0.943, 0.959] | 0.956 [0.948, 0.964] | 0.812 [0.804, 0.820] | 0.632 [0.624, 0.640] |
| | 30 | 0.965 [0.955, 0.975] | 0.963 [0.952, 0.973] | 0.950 [0.939, 0.960] | 0.954 [0.944, 0.965] | 0.960 [0.950, 0.970] | 0.900 [0.890, 0.910] | 0.653 [0.643, 0.664] |
| | 50 | 0.968 [0.956, 0.979] | 0.962 [0.950, 0.974] | 0.951 [0.940, 0.963] | 0.953 [0.941, 0.964] | 0.961 [0.949, 0.972] | 0.914 [0.903, 0.926] | 0.661 [0.649, 0.673] |
| | 100 | 0.967 [0.952, 0.982] | 0.958 [0.943, 0.973] | 0.949 [0.934, 0.964] | 0.949 [0.934, 0.963] | 0.959 [0.945, 0.973] | 0.924 [0.910, 0.939] | 0.664 [0.650, 0.680] |
| HG004-MasSeq | 1 | 0.779 [0.774, 0.784] | 0.776 [0.771, 0.781] | 0.773 [0.768, 0.778] | 0.728 [0.724, 0.733] | 0.793 [0.788, 0.797] | 0.504 [0.499, 0.509] | 0.569 [0.564, 0.573] |
| | 5 | 0.930 [0.925, 0.936] | 0.926 [0.921, 0.932] | 0.923 [0.917, 0.928] | 0.914 [0.909, 0.920] | 0.943 [0.937, 0.948] | 0.666 [0.660, 0.672] | 0.701 [0.696, 0.706] |
| | 10 | 0.949 [0.943, 0.955] | 0.948 [0.942, 0.954] | 0.946 [0.939, 0.952] | 0.952 [0.946, 0.958] | 0.965 [0.958, 0.971] | 0.747 [0.741, 0.754] | 0.737 [0.731, 0.743] |
| | 30 | 0.957 [0.950, 0.965] | 0.963 [0.955, 0.970] | 0.959 [0.952, 0.967] | 0.965 [0.958, 0.973] | 0.979 [0.971, 0.986] | 0.848 [0.840, 0.856] | 0.775 [0.768, 0.782] |
| | 50 | 0.959 [0.951, 0.967] | 0.967 [0.959, 0.975] | 0.962 [0.954, 0.970] | 0.967 [0.959, 0.975] | 0.981 [0.973, 0.989] | 0.879 [0.871, 0.888] | 0.786 [0.779, 0.795] |
| | 100 | 0.959 [0.950, 0.969] | 0.970 [0.961, 0.979] | 0.961 [0.952, 0.970] | 0.966 [0.957, 0.976] | 0.982 [0.973, 0.991] | 0.910 [0.901, 0.920] | 0.798 [0.789, 0.807] |
| HG004-cDNAxR09 | 1 | 0.603 [0.598, 0.609] | 0.492 [0.487, 0.496] | 0.582 [0.577, 0.587] | 0.542 [0.536, 0.548] | 0.597 [0.592, 0.602] | 0.285 [0.280, 0.290] | 0.199 [0.195, 0.203] |
| | 5 | 0.759 [0.753, 0.766] | 0.601 [0.595, 0.607] | 0.737 [0.730, 0.744] | 0.713 [0.706, 0.720] | 0.734 [0.728, 0.740] | 0.411 [0.404, 0.418] | 0.264 [0.259, 0.270] |
| | 10 | 0.828 [0.820, 0.836] | 0.673 [0.666, 0.680] | 0.817 [0.809, 0.825] | 0.808 [0.800, 0.816] | 0.821 [0.814, 0.829] | 0.504 [0.496, 0.513] | 0.296 [0.290, 0.303] |
| | 30 | 0.894 [0.884, 0.904] | 0.748 [0.738, 0.758] | 0.887 [0.876, 0.897] | 0.900 [0.890, 0.910] | 0.919 [0.909, 0.929] | 0.644 [0.633, 0.655] | 0.346 [0.338, 0.355] |
| | 50 | 0.903 [0.891, 0.914] | 0.737 [0.726, 0.749] | 0.896 [0.884, 0.907] | 0.915 [0.904, 0.927] | 0.933 [0.921, 0.944] | 0.705 [0.693, 0.717] | 0.368 [0.359, 0.378] |
| | 100 | 0.906 [0.892, 0.920] | 0.677 [0.662, 0.691] | 0.895 [0.881, 0.910] | 0.923 [0.908, 0.937] | 0.938 [0.924, 0.952] | 0.771 [0.756, 0.785] | 0.396 [0.384, 0.408] |
| HG004-cDNAxR10 | 1 | 0.674 [0.669, 0.679] | 0.552 [0.547, 0.557] | 0.644 [0.639, 0.649] | 0.524 [0.519, 0.529] | 0.698 [0.693, 0.703] | 0.340 [0.335, 0.345] | 0.346 [0.341, 0.350] |
| | 5 | 0.867 [0.861, 0.874] | 0.691 [0.685, 0.697] | 0.838 [0.832, 0.844] | 0.735 [0.728, 0.741] | 0.880 [0.874, 0.886] | 0.507 [0.500, 0.513] | 0.463 [0.457, 0.469] |
| | 10 | 0.906 [0.898, 0.913] | 0.728 [0.721, 0.734] | 0.884 [0.877, 0.891] | 0.814 [0.807, 0.822] | 0.923 [0.915, 0.930] | 0.596 [0.588, 0.604] | 0.503 [0.497, 0.510] |
| | 30 | 0.930 [0.921, 0.939] | 0.768 [0.759, 0.776] | 0.908 [0.900, 0.917] | 0.864 [0.855, 0.873] | 0.952 [0.944, 0.961] | 0.714 [0.705, 0.723] | 0.546 [0.538, 0.554] |
| | 50 | 0.933 [0.923, 0.943] | 0.758 [0.748, 0.768] | 0.911 [0.902, 0.921] | 0.871 [0.861, 0.880] | 0.957 [0.948, 0.967] | 0.759 [0.750, 0.769] | 0.556 [0.548, 0.566] |
| | 100 | 0.936 [0.925, 0.948] | 0.716 [0.705, 0.728] | 0.914 [0.903, 0.926] | 0.874 [0.863, 0.886] | 0.961 [0.950, 0.972] | 0.809 [0.798, 0.821] | 0.567 [0.557, 0.577] |
| HG004-dRNA002 | 1 | 0.426 [0.420, 0.432] | 0.408 [0.402, 0.415] | 0.443 [0.437, 0.450] | 0.178 [0.173, 0.183] | 0.273 [0.269, 0.277] | 0.083 [0.079, 0.087] | 0.012 [0.011, 0.014] |
| | 5 | 0.596 [0.587, 0.604] | 0.585 [0.575, 0.595] | 0.632 [0.622, 0.642] | 0.327 [0.318, 0.336] | 0.330 [0.325, 0.335] | 0.161 [0.154, 0.169] | 0.023 [0.021, 0.027] |
| | 10 | 0.623 [0.613, 0.633] | 0.599 [0.588, 0.611] | 0.668 [0.657, 0.680] | 0.389 [0.378, 0.401] | 0.374 [0.367, 0.381] | 0.215 [0.206, 0.225] | 0.029 [0.025, 0.033] |
| | 30 | 0.654 [0.640, 0.668] | 0.570 [0.554, 0.587] | 0.693 [0.677, 0.709] | 0.351 [0.335, 0.367] | 0.486 [0.475, 0.498] | 0.335 [0.320, 0.350] | 0.035 [0.029, 0.041] |
| | 50 | 0.663 [0.646, 0.680] | 0.530 [0.510, 0.551] | 0.688 [0.669, 0.708] | 0.314 [0.295, 0.333] | 0.527 [0.512, 0.543] | 0.391 [0.373, 0.410] | 0.038 [0.031, 0.047] |
| | 100 | 0.664 [0.640, 0.687] | 0.477 [0.450, 0.504] | 0.665 [0.638, 0.693] | 0.199 [0.178, 0.223] | 0.538 [0.516, 0.561] | 0.456 [0.431, 0.482] | 0.041 [0.031, 0.054] |
| HG004-dRNA004 | 1 | 0.764 [0.759, 0.770] | 0.639 [0.634, 0.645] | 0.768 [0.762, 0.773] | 0.706 [0.700, 0.711] | 0.780 [0.775, 0.785] | 0.403 [0.398, 0.408] | 0.475 [0.470, 0.480] |
| | 5 | 0.921 [0.915, 0.927] | 0.768 [0.762, 0.774] | 0.910 [0.904, 0.916] | 0.888 [0.881, 0.894] | 0.919 [0.913, 0.925] | 0.541 [0.535, 0.548] | 0.600 [0.594, 0.607] |
| | 10 | 0.944 [0.937, 0.951] | 0.784 [0.777, 0.791] | 0.929 [0.922, 0.936] | 0.934 [0.928, 0.941] | 0.939 [0.932, 0.945] | 0.614 [0.607, 0.621] | 0.641 [0.634, 0.648] |
| | 30 | 0.957 [0.949, 0.965] | 0.775 [0.767, 0.783] | 0.939 [0.932, 0.947] | 0.946 [0.938, 0.954] | 0.952 [0.945, 0.960] | 0.717 [0.709, 0.726] | 0.681 [0.674, 0.690] |
| | 50 | 0.961 [0.952, 0.969] | 0.755 [0.746, 0.764] | 0.941 [0.932, 0.949] | 0.946 [0.938, 0.955] | 0.959 [0.950, 0.967] | 0.761 [0.752, 0.770] | 0.693 [0.685, 0.702] |
| | 100 | 0.963 [0.954, 0.973] | 0.708 [0.697, 0.718] | 0.938 [0.928, 0.948] | 0.943 [0.933, 0.953] | 0.962 [0.953, 0.972] | 0.806 [0.796, 0.816] | 0.699 [0.689, 0.709] |
| HG005-IsoSeq | 1 | 0.790 [0.785, 0.795] | 0.774 [0.769, 0.780] | 0.731 [0.726, 0.737] | 0.730 [0.725, 0.735] | 0.790 [0.785, 0.795] | 0.567 [0.562, 0.572] | 0.506 [0.501, 0.511] |
| | 5 | 0.949 [0.942, 0.955] | 0.940 [0.933, 0.946] | 0.905 [0.899, 0.911] | 0.933 [0.926, 0.939] | 0.945 [0.939, 0.951] | 0.762 [0.755, 0.769] | 0.651 [0.644, 0.657] |
| | 10 | 0.965 [0.958, 0.972] | 0.959 [0.951, 0.966] | 0.936 [0.929, 0.943] | 0.958 [0.951, 0.966] | 0.964 [0.956, 0.971] | 0.860 [0.852, 0.867] | 0.688 [0.681, 0.696] |
| | 30 | 0.971 [0.961, 0.980] | 0.965 [0.955, 0.975] | 0.952 [0.943, 0.962] | 0.966 [0.956, 0.975] | 0.971 [0.961, 0.980] | 0.917 [0.908, 0.926] | 0.710 [0.701, 0.720] |
| | 50 | 0.971 [0.960, 0.982] | 0.965 [0.953, 0.976] | 0.952 [0.941, 0.963] | 0.965 [0.954, 0.976] | 0.971 [0.960, 0.981] | 0.924 [0.914, 0.935] | 0.718 [0.707, 0.730] |
| | 100 | 0.971 [0.957, 0.985] | 0.975 [0.960, 0.989] | 0.952 [0.938, 0.966] | 0.963 [0.949, 0.977] | 0.971 [0.957, 0.984] | 0.932 [0.919, 0.946] | 0.727 [0.714, 0.742] |

*Continued on next page*

Table 7 – continued from previous page

| Dataset | Coverage | Clair3-RNA | DeepVariant | longcallR | longcallR-nn | NanoTS | isoLASER | GATK |
|---|---|---|---|---|---|---|---|---|
| HG005-MasSeq | 1 | 0.818 [0.813, 0.823] | 0.815 [0.810, 0.820] | 0.808 [0.803, 0.812] | 0.768 [0.763, 0.773] | 0.830 [0.826, 0.835] | 0.544 [0.539, 0.549] | 0.607 [0.602, 0.611] |
| | 5 | 0.940 [0.935, 0.946] | 0.934 [0.929, 0.940] | 0.928 [0.922, 0.933] | 0.923 [0.918, 0.929] | 0.950 [0.945, 0.956] | 0.683 [0.678, 0.689] | 0.713 [0.708, 0.719] |
| | 10 | 0.956 [0.950, 0.962] | 0.952 [0.946, 0.958] | 0.948 [0.942, 0.954] | 0.958 [0.952, 0.964] | 0.968 [0.962, 0.974] | 0.758 [0.752, 0.765] | 0.745 [0.739, 0.751] |
| | 30 | 0.964 [0.956, 0.971] | 0.965 [0.957, 0.972] | 0.959 [0.952, 0.967] | 0.971 [0.963, 0.978] | 0.980 [0.973, 0.987] | 0.854 [0.847, 0.862] | 0.782 [0.775, 0.789] |
| | 50 | 0.964 [0.956, 0.972] | 0.967 [0.959, 0.975] | 0.960 [0.952, 0.968] | 0.971 [0.963, 0.979] | 0.983 [0.975, 0.991] | 0.882 [0.874, 0.890] | 0.795 [0.787, 0.803] |
| | 100 | 0.963 [0.954, 0.972] | 0.972 [0.963, 0.981] | 0.960 [0.951, 0.969] | 0.972 [0.963, 0.981] | 0.987 [0.978, 0.996] | 0.912 [0.903, 0.921] | 0.806 [0.798, 0.816] |
| HG005-cDNAxR09 | 1 | 0.661 [0.656, 0.667] | 0.536 [0.531, 0.541] | 0.638 [0.633, 0.644] | 0.580 [0.575, 0.586] | 0.665 [0.660, 0.670] | 0.316 [0.311, 0.321] | 0.245 [0.241, 0.250] |
| | 5 | 0.840 [0.833, 0.846] | 0.664 [0.658, 0.670] | 0.816 [0.809, 0.822] | 0.769 [0.763, 0.776] | 0.819 [0.812, 0.825] | 0.457 [0.451, 0.464] | 0.334 [0.329, 0.340] |
| | 10 | 0.885 [0.878, 0.893] | 0.715 [0.708, 0.722] | 0.868 [0.861, 0.875] | 0.849 [0.842, 0.856] | 0.878 [0.871, 0.885] | 0.544 [0.536, 0.551] | 0.370 [0.364, 0.377] |
| | 30 | 0.914 [0.905, 0.923] | 0.755 [0.746, 0.764] | 0.902 [0.893, 0.911] | 0.903 [0.894, 0.911] | 0.937 [0.928, 0.946] | 0.672 [0.663, 0.681] | 0.414 [0.406, 0.422] |
| | 50 | 0.917 [0.907, 0.927] | 0.749 [0.739, 0.759] | 0.907 [0.897, 0.917] | 0.910 [0.900, 0.920] | 0.946 [0.936, 0.956] | 0.722 [0.712, 0.732] | 0.434 [0.425, 0.443] |
| | 100 | 0.914 [0.902, 0.926] | 0.709 [0.697, 0.721] | 0.904 [0.892, 0.916] | 0.917 [0.906, 0.929] | 0.949 [0.937, 0.960] | 0.772 [0.760, 0.784] | 0.461 [0.451, 0.473] |
| HG005-cDNAxR10 | 1 | 0.708 [0.703, 0.713] | 0.560 [0.555, 0.564] | 0.678 [0.673, 0.683] | 0.557 [0.552, 0.562] | 0.728 [0.724, 0.733] | 0.369 [0.365, 0.374] | 0.394 [0.390, 0.399] |
| | 5 | 0.871 [0.865, 0.877] | 0.671 [0.666, 0.677] | 0.839 [0.833, 0.845] | 0.736 [0.730, 0.742] | 0.881 [0.875, 0.886] | 0.514 [0.508, 0.520] | 0.498 [0.493, 0.504] |
| | 10 | 0.904 [0.898, 0.911] | 0.703 [0.697, 0.709] | 0.878 [0.871, 0.884] | 0.803 [0.796, 0.810] | 0.917 [0.911, 0.924] | 0.595 [0.588, 0.602] | 0.532 [0.526, 0.538] |
| | 30 | 0.926 [0.918, 0.934] | 0.747 [0.739, 0.755] | 0.897 [0.889, 0.905] | 0.850 [0.842, 0.859] | 0.947 [0.939, 0.955] | 0.709 [0.701, 0.718] | 0.574 [0.567, 0.581] |
| | 50 | 0.931 [0.922, 0.940] | 0.752 [0.744, 0.761] | 0.901 [0.892, 0.909] | 0.863 [0.854, 0.872] | 0.953 [0.944, 0.962] | 0.754 [0.745, 0.763] | 0.586 [0.579, 0.595] |
| | 100 | 0.933 [0.922, 0.943] | 0.731 [0.721, 0.741] | 0.905 [0.895, 0.915] | 0.872 [0.861, 0.882] | 0.958 [0.948, 0.968] | 0.802 [0.792, 0.813] | 0.599 [0.590, 0.608] |
| HG005-dRNA002 | 1 | 0.443 [0.437, 0.449] | 0.448 [0.441, 0.455] | 0.462 [0.455, 0.469] | 0.197 [0.192, 0.203] | 0.280 [0.276, 0.284] | 0.088 [0.084, 0.092] | 0.013 [0.011, 0.014] |
| | 5 | 0.607 [0.598, 0.615] | 0.631 [0.621, 0.641] | 0.647 [0.637, 0.657] | 0.355 [0.345, 0.364] | 0.336 [0.331, 0.341] | 0.168 [0.161, 0.175] | 0.025 [0.022, 0.028] |
| | 10 | 0.630 [0.620, 0.640] | 0.648 [0.636, 0.659] | 0.679 [0.667, 0.690] | 0.417 [0.406, 0.429] | 0.385 [0.378, 0.392] | 0.223 [0.214, 0.233] | 0.030 [0.026, 0.034] |
| | 30 | 0.655 [0.642, 0.669] | 0.621 [0.605, 0.638] | 0.697 [0.681, 0.714] | 0.366 [0.350, 0.383] | 0.504 [0.492, 0.516] | 0.349 [0.334, 0.365] | 0.037 [0.031, 0.043] |
| | 50 | 0.665 [0.648, 0.682] | 0.588 [0.567, 0.609] | 0.685 [0.665, 0.705] | 0.314 [0.296, 0.334] | 0.549 [0.533, 0.565] | 0.405 [0.386, 0.424] | 0.038 [0.031, 0.047] |
| | 100 | 0.663 [0.639, 0.687] | 0.578 [0.549, 0.607] | 0.658 [0.630, 0.686] | 0.198 [0.176, 0.222] | 0.573 [0.550, 0.597] | 0.485 [0.459, 0.511] | 0.046 [0.035, 0.059] |
| HG005-dRNA004 | 1 | 0.778 [0.773, 0.783] | 0.673 [0.668, 0.678] | 0.776 [0.771, 0.781] | 0.715 [0.710, 0.720] | 0.795 [0.789, 0.800] | 0.394 [0.390, 0.399] | 0.523 [0.518, 0.528] |
| | 5 | 0.932 [0.926, 0.938] | 0.803 [0.797, 0.809] | 0.914 [0.907, 0.920] | 0.898 [0.892, 0.904] | 0.928 [0.922, 0.934] | 0.532 [0.525, 0.538] | 0.646 [0.641, 0.653] |
| | 10 | 0.950 [0.944, 0.957] | 0.817 [0.810, 0.823] | 0.928 [0.922, 0.935] | 0.941 [0.935, 0.948] | 0.944 [0.937, 0.950] | 0.603 [0.596, 0.610] | 0.680 [0.674, 0.687] |
| | 30 | 0.960 [0.952, 0.968] | 0.814 [0.806, 0.822] | 0.935 [0.928, 0.943] | 0.952 [0.944, 0.960] | 0.954 [0.946, 0.962] | 0.709 [0.700, 0.717] | 0.716 [0.709, 0.725] |
| | 50 | 0.963 [0.954, 0.971] | 0.802 [0.793, 0.811] | 0.936 [0.927, 0.945] | 0.953 [0.944, 0.961] | 0.959 [0.951, 0.968] | 0.754 [0.745, 0.763] | 0.728 [0.720, 0.738] |
| | 100 | 0.963 [0.954, 0.973] | 0.765 [0.755, 0.775] | 0.933 [0.923, 0.943] | 0.949 [0.939, 0.959] | 0.961 [0.951, 0.970] | 0.797 [0.787, 0.808] | 0.738 [0.729, 0.749] |

F1 [95% CI] shown for each variant caller, dataset (cell line-chemistry), and coverage cutoff (DP $\geq$ n). Credible intervals were computed using a Beta-Binomial model with a Jeffreys prior ($\alpha = \beta = 0.5$), based on the identity F1 = 2·TP / (TRUTH.TOTAL + QUERY.TOTAL), equivalent to the interval computed by happyCompare (`https://github.com/Illumina/happyCompare`) for hap.py output.

**Table S8:** F1 scores with 95% Jeffreys credible intervals for indel calling on GIAB HG004 and HG005 datasets.

| Dataset | Coverage | Clair3-RNA | DeepVariant | isoLASER | GATK |
|---|---|---|---|---|---|
| HG004-IsoSeq | 1 | 0.587 [0.573, 0.601] | 0.585 [0.567, 0.596] | 0.184 [0.175, 0.194] | 0.348 [0.335, 0.358] |
| | 5 | 0.766 [0.748, 0.784] | 0.749 [0.724, 0.762] | 0.297 [0.281, 0.312] | 0.456 [0.437, 0.468] |
| | 10 | 0.815 [0.794, 0.837] | 0.781 [0.752, 0.796] | 0.382 [0.363, 0.402] | 0.483 [0.462, 0.497] |
| | 30 | 0.855 [0.826, 0.884] | 0.792 [0.753, 0.811] | 0.483 [0.456, 0.511] | 0.499 [0.470, 0.516] |
| | 50 | 0.872 [0.838, 0.906] | 0.786 [0.743, 0.810] | 0.511 [0.479, 0.543] | 0.503 [0.469, 0.522] |
| | 100 | 0.877 [0.833, 0.921] | 0.762 [0.708, 0.792] | 0.520 [0.478, 0.560] | 0.500 [0.456, 0.524] |
| HG004-MasSeq | 1 | 0.613 [0.601, 0.626] | 0.642 [0.626, 0.651] | 0.211 [0.202, 0.221] | 0.243 [0.233, 0.246] |
| | 5 | 0.740 [0.726, 0.755] | 0.777 [0.756, 0.786] | 0.293 [0.280, 0.306] | 0.260 [0.249, 0.263] |
| | 10 | 0.770 [0.754, 0.787] | 0.811 [0.787, 0.821] | 0.342 [0.327, 0.357] | 0.258 [0.247, 0.261] |
| | 30 | 0.806 [0.787, 0.827] | 0.853 [0.825, 0.866] | 0.437 [0.417, 0.457] | 0.253 [0.240, 0.256] |
| | 50 | 0.816 [0.794, 0.839] | 0.867 [0.835, 0.880] | 0.474 [0.452, 0.497] | 0.249 [0.234, 0.252] |
| | 100 | 0.829 [0.803, 0.856] | 0.881 [0.845, 0.898] | 0.513 [0.486, 0.539] | 0.249 [0.232, 0.253] |
| HG004-cDNAxR09 | 1 | 0.342 [0.329, 0.355] | 0.305 [0.292, 0.316] | – | 0.058 [0.053, 0.061] |
| | 5 | 0.449 [0.432, 0.466] | 0.394 [0.377, 0.407] | – | 0.064 [0.059, 0.068] |
| | 10 | 0.506 [0.486, 0.526] | 0.446 [0.425, 0.462] | – | 0.064 [0.059, 0.069] |
| | 30 | 0.585 [0.559, 0.612] | 0.509 [0.481, 0.532] | – | 0.064 [0.059, 0.069] |
| | 50 | 0.603 [0.573, 0.635] | 0.511 [0.477, 0.537] | – | 0.064 [0.057, 0.069] |
| | 100 | 0.623 [0.584, 0.662] | 0.481 [0.442, 0.517] | – | 0.064 [0.057, 0.070] |
| HG004-cDNAxR10 | 1 | 0.478 [0.465, 0.491] | 0.433 [0.419, 0.445] | – | 0.192 [0.183, 0.198] |
| | 5 | 0.642 [0.625, 0.660] | 0.594 [0.574, 0.608] | – | 0.233 [0.221, 0.240] |
| | 10 | 0.687 [0.668, 0.707] | 0.646 [0.623, 0.663] | – | 0.244 [0.230, 0.250] |
| | 30 | 0.729 [0.705, 0.754] | 0.683 [0.654, 0.704] | – | 0.250 [0.234, 0.257] |
| | 50 | 0.747 [0.720, 0.775] | 0.687 [0.654, 0.711] | – | 0.253 [0.236, 0.261] |
| | 100 | 0.760 [0.727, 0.794] | 0.655 [0.616, 0.686] | – | 0.246 [0.228, 0.256] |
| HG004-dRNA002 | 1 | 0.092 [0.082, 0.102] | 0.054 [0.049, 0.060] | – | 0.002 [0.001, 0.004] |
| | 5 | 0.166 [0.148, 0.186] | 0.070 [0.062, 0.078] | – | 0.003 [0.001, 0.007] |
| | 10 | 0.196 [0.173, 0.220] | 0.074 [0.065, 0.083] | – | 0.004 [0.002, 0.009] |
| | 30 | 0.217 [0.185, 0.253] | 0.070 [0.058, 0.084] | – | 0.005 [0.002, 0.013] |
| | 50 | 0.238 [0.196, 0.285] | 0.074 [0.059, 0.091] | – | 0.004 [0.001, 0.012] |
| | 100 | 0.213 [0.154, 0.284] | 0.069 [0.048, 0.095] | – | 0.003 [0.000, 0.015] |
| HG004-dRNA004 | 1 | 0.495 [0.481, 0.509] | 0.369 [0.356, 0.379] | – | 0.081 [0.077, 0.083] |
| | 5 | 0.625 [0.608, 0.642] | 0.450 [0.433, 0.461] | – | 0.083 [0.079, 0.085] |
| | 10 | 0.661 [0.642, 0.679] | 0.467 [0.449, 0.480] | – | 0.081 [0.077, 0.083] |
| | 30 | 0.690 [0.669, 0.713] | 0.477 [0.456, 0.493] | – | 0.076 [0.072, 0.078] |
| | 50 | 0.697 [0.673, 0.721] | 0.469 [0.446, 0.486] | – | 0.074 [0.069, 0.076] |
| | 100 | 0.704 [0.676, 0.731] | 0.457 [0.432, 0.477] | – | 0.073 [0.067, 0.074] |
| HG005-IsoSeq | 1 | 0.680 [0.665, 0.696] | 0.681 [0.663, 0.694] | 0.265 [0.254, 0.277] | 0.429 [0.413, 0.439] |
| | 5 | 0.834 [0.816, 0.853] | 0.846 [0.822, 0.861] | 0.402 [0.385, 0.419] | 0.524 [0.504, 0.536] |
| | 10 | 0.876 [0.855, 0.898] | 0.887 [0.859, 0.904] | 0.492 [0.472, 0.512] | 0.545 [0.523, 0.559] |
| | 30 | 0.901 [0.873, 0.930] | 0.921 [0.884, 0.944] | 0.561 [0.534, 0.589] | 0.558 [0.528, 0.575] |
| | 50 | 0.907 [0.874, 0.942] | 0.930 [0.887, 0.958] | 0.569 [0.537, 0.602] | 0.562 [0.526, 0.580] |
| | 100 | 0.907 [0.863, 0.951] | 0.935 [0.881, 0.974] | 0.577 [0.537, 0.619] | 0.563 [0.519, 0.588] |

*Continued on next page*

Table 8 – continued from previous page

| Dataset | Coverage | Clair3-RNA | DeepVariant | isoLASER | GATK |
|---|---|---|---|---|---|
| HG005-MasSeq | 1 | 0.708 [0.694, 0.722] | 0.737 [0.719, 0.747] | 0.257 [0.246, 0.268] | 0.383 [0.369, 0.388] |
| | 5 | 0.823 [0.807, 0.839] | 0.854 [0.832, 0.864] | 0.336 [0.322, 0.350] | 0.416 [0.400, 0.420] |
| | 10 | 0.855 [0.838, 0.873] | 0.885 [0.861, 0.897] | 0.391 [0.376, 0.408] | 0.422 [0.405, 0.426] |
| | 30 | 0.885 [0.864, 0.906] | 0.920 [0.892, 0.935] | 0.482 [0.462, 0.504] | 0.426 [0.406, 0.431] |
| | 50 | 0.892 [0.868, 0.916] | 0.927 [0.897, 0.944] | 0.516 [0.493, 0.540] | 0.427 [0.405, 0.433] |
| | 100 | 0.902 [0.874, 0.930] | 0.932 [0.897, 0.953] | 0.562 [0.535, 0.591] | 0.422 [0.397, 0.430] |
| HG005-cDNAxR09 | 1 | 0.437 [0.422, 0.451] | 0.374 [0.360, 0.386] | – | 0.069 [0.064, 0.072] |
| | 5 | 0.568 [0.550, 0.586] | 0.477 [0.458, 0.491] | – | 0.075 [0.069, 0.078] |
| | 10 | 0.618 [0.597, 0.639] | 0.521 [0.499, 0.538] | – | 0.075 [0.069, 0.079] |
| | 30 | 0.669 [0.643, 0.695] | 0.546 [0.520, 0.569] | – | 0.072 [0.066, 0.076] |
| | 50 | 0.681 [0.652, 0.711] | 0.541 [0.512, 0.567] | – | 0.070 [0.064, 0.074] |
| | 100 | 0.698 [0.663, 0.735] | 0.515 [0.481, 0.547] | – | 0.067 [0.059, 0.071] |
| HG005-cDNAxR10 | 1 | 0.561 [0.547, 0.576] | 0.484 [0.469, 0.496] | – | 0.234 [0.224, 0.240] |
| | 5 | 0.708 [0.691, 0.726] | 0.626 [0.606, 0.641] | – | 0.270 [0.257, 0.276] |
| | 10 | 0.754 [0.734, 0.774] | 0.676 [0.653, 0.693] | – | 0.278 [0.264, 0.284] |
| | 30 | 0.798 [0.774, 0.822] | 0.706 [0.679, 0.727] | – | 0.280 [0.264, 0.286] |
| | 50 | 0.812 [0.786, 0.839] | 0.718 [0.688, 0.742] | – | 0.276 [0.259, 0.283] |
| | 100 | 0.828 [0.796, 0.860] | 0.698 [0.662, 0.728] | – | 0.273 [0.253, 0.280] |
| HG005-dRNA002 | 1 | 0.114 [0.102, 0.127] | 0.066 [0.059, 0.073] | – | 0.001 [0.000, 0.003] |
| | 5 | 0.198 [0.177, 0.220] | 0.082 [0.074, 0.092] | – | 0.002 [0.001, 0.006] |
| | 10 | 0.224 [0.199, 0.251] | 0.083 [0.073, 0.094] | – | 0.003 [0.001, 0.008] |
| | 30 | 0.272 [0.234, 0.314] | 0.084 [0.070, 0.098] | – | 0.003 [0.001, 0.011] |
| | 50 | 0.280 [0.232, 0.334] | 0.083 [0.067, 0.102] | – | 0.005 [0.001, 0.015] |
| | 100 | 0.297 [0.226, 0.381] | 0.081 [0.058, 0.110] | – | 0.004 [0.000, 0.019] |
| HG005-dRNA004 | 1 | 0.553 [0.538, 0.568] | 0.439 [0.424, 0.450] | – | 0.088 [0.084, 0.089] |
| | 5 | 0.688 [0.670, 0.706] | 0.534 [0.515, 0.547] | – | 0.088 [0.084, 0.090] |
| | 10 | 0.723 [0.703, 0.743] | 0.557 [0.536, 0.571] | – | 0.085 [0.080, 0.086] |
| | 30 | 0.749 [0.726, 0.772] | 0.572 [0.549, 0.590] | – | 0.080 [0.075, 0.081] |
| | 50 | 0.754 [0.728, 0.779] | 0.566 [0.540, 0.586] | – | 0.078 [0.073, 0.079] |
| | 100 | 0.759 [0.730, 0.789] | 0.547 [0.519, 0.571] | – | 0.076 [0.070, 0.078] |

F1 [95% CI] shown for each variant caller, dataset (cell line-chemistry), and coverage cutoff (DP $\geq$ n). Credible intervals were computed using a Beta-Binomial model with a Jeffreys prior ($\alpha = \beta = 0.5$), based on the identity F1 = 2·TP / (TRUTH.TOTAL + QUERY.TOTAL), equivalent to the interval computed by happyCompare (`https://github.com/Illumina/happyCompare`) for hap.py output.

#### 1.4 pbmm2 vs. minimap2

| Dataset | minimap2 (%) | pbmm2 (%) |
|---|---|---|
| HG002-MasSeq | 2.40 | 2.59 |
| HG002-IsoSeq | 4.84 | 5.18 |
| HG004-MasSeq | 1.11 | 1.63 |
| HG004-IsoSeq | 4.33 | 4.33 |
| HG005-MasSeq | 1.34 | 3.31 |
| HG005-IsoSeq | 7.71 | 8.11 |

**Table S9: Supplementary read alignment rates with minimap2 and pbmm2 across PacBio datasets.**

| Parameter | pbmm2 | minimap2 | Definition |
|---|---|---|---|
| Splice annotation (`--junc-bed`) | not supported | transcriptome BED | known junctions given to the aligner |
| Junction bonus (`--junc-bonus`) | 0 | 9 | score added at an annotated splice site |
| SW matrix cap (`--cap-sw-mat`) | $10^8$ | 0 (off) | matrix size above which alignment is skipped |
| Repeated-matches trimming | on | off | query overlap allowed between alignments |
| Secondary alignments | none | 5 | alternate mappings reported |
| All other parameters | identical | | $-A$, $-B$, $-o$, $-O$, $-e$, $-E$, $-z$, $-Z$, $-r$, $-g$, $-G$, $-k$, $-w$, $-C$ |

**Table S10: Parameter differences between `pbmm2 --preset ISOSEQ` and `minimap2 -x splice:hq`.** Alignment parameter differences that may contribute to the observed variant-calling performance differences between pbmm2 and minimap2.

#### 1.5 LongBench datasets

| Factor | F1 | $\log_{10}$(TP + 1) |
|---|---|---|
| Variant type (SNV/INDEL) | 0.786 | 0.966 |
| Variant caller | 0.657 | 0.573 |
| Coverage cutoff | 0.608 | 0.879 |
| Chemistry | 0.482 | 0.919 |
| Cell line | 0.070 | 0.143 |
| Full model $R^2$ | 0.902 | 0.982 |

**Table S11: Partial $R^2$ of each factor for F1 score and number of true positives.** Linear models F1 $\sim$ caller + chemistry + variant type + coverage cutoff + cell line (and the same for $\log_{10}$(TP + 1)), fitted on PASS calls of all 300 caller × sample × variant type × coverage combinations. Partial $R^2$ = ($SSE_{reduced}$ − $SSE_{full}$) / $SSE_{reduced}$, where the reduced model drops that factor. INDEL results of SNV-only callers (longcallR, longcallR-nn, NanoTS) and of isoLASER outside Mas-Seq are excluded, as in the main figures.

| Type | Variant Caller | 1 | 5 | 10 | 30 | 50 | 100 |
|---|---|---|---|---|---|---|---|
| SNV | Clair3-RNA | 0.43 | −0.26 | −0.43 | −0.77 | −0.77 | −0.66 |
| | DeepVariant | 0.43 | 0.43 | 0.43 | 0.54 | 0.54 | 0.54 |
| | isoLASER | 0.43 | 0.43 | 0.54 | 0.54 | 0.54 | 0.54 |
| | longcallR | 0.43 | 0.20 | 0.20 | 0.20 | 0.03 | 0.20 |
| | longcallR-nn | 0.43 | 0.43 | 0.20 | 0.20 | 0.20 | 0.20 |
| | NanoTS | 0.43 | 0.43 | 0.54 | 0.37 | 0.14 | −0.03 |
| INDEL | Clair3-RNA | 0.43 | 0.71 | 0.77 | 0.89 | 0.89 | 0.89 |
| | DeepVariant | 0.49 | 0.94 | 1.00 | 1.00 | 1.00 | 0.94 |

**Table S12: Spearman correlation between (1 − error rate) and F1 score.** Correlations are computed per variant caller, variant type, and coverage cutoff, across all samples and cell lines (H211, H526) and sequencing platforms/chemistries (MasSeq, cDNAxR10, dRNA004) ($n = 6$ per row). GATK is not benchmarked as an RNA-seq variant caller in LongBench (used only to build the WGS ground truth) and is omitted. longcallR, longcallR-nn and NanoTS call SNVs only and are omitted from the INDEL block. isoLASER INDEL calls are restricted to PacBio (MasSeq) samples ($n = 2$), below the minimum needed for a Spearman test, and are omitted.

| Type | Variant Caller | 1 | 5 | 10 | 30 | 50 | 100 |
|---|---|---|---|---|---|---|---|
| SNV | Clair3-RNA | 1.00 | 1.00 | 1.00 | 1.00 | 1.00 | 1.00 |
| | DeepVariant | 1.00 | 1.00 | 1.00 | 1.00 | 1.00 | 1.00 |
| | isoLASER | 0.94 | 0.94 | 0.94 | 0.94 | 0.94 | 0.94 |
| | longcallR | 1.00 | 1.00 | 1.00 | 1.00 | 1.00 | 1.00 |
| | longcallR-nn | 1.00 | 1.00 | 1.00 | 1.00 | 1.00 | 1.00 |
| | NanoTS | 1.00 | 1.00 | 1.00 | 1.00 | 1.00 | 1.00 |
| INDEL | Clair3-RNA | 1.00 | 1.00 | 1.00 | 1.00 | 1.00 | 1.00 |
| | DeepVariant | 1.00 | 1.00 | 1.00 | 1.00 | 1.00 | 1.00 |

**Table S13: Spearman correlation between callable bases and the number of true positives.** Correlations are computed per variant caller, variant type, and coverage cutoff, across all samples and cell lines (H211, H526) and sequencing platforms/chemistries (MasSeq, cDNAxR10, dRNA004) ($n = 6$ per row). Callable bases are the exonic bases on chr1–22 with coverage $\geq n$, i.e. the evaluation region. longcallR, longcallR-nn and NanoTS call SNVs only and are omitted from the INDEL block. isoLASER INDEL calls are restricted to PacBio (MasSeq) samples ($n = 2$), below the minimum needed for a Spearman test, and are omitted.

| Dataset | Coverage | Clair3-RNA | DeepVariant | isoLASER | longcallR | longcallR-nn | NanoTS |
|---|---|---|---|---|---|---|---|
| H211-MasSeq | 1 | 0.756 [0.752, 0.760] | 0.781 [0.776, 0.785] | 0.590 [0.586, 0.595] | 0.783 [0.779, 0.788] | 0.728 [0.723, 0.732] | 0.790 [0.785, 0.794] |
| | 5 | 0.837 [0.832, 0.842] | 0.865 [0.860, 0.870] | 0.712 [0.707, 0.717] | 0.868 [0.863, 0.873] | 0.847 [0.842, 0.852] | 0.873 [0.867, 0.877] |
| | 10 | 0.844 [0.838, 0.849] | 0.877 [0.871, 0.883] | 0.772 [0.767, 0.778] | 0.882 [0.876, 0.887] | 0.870 [0.865, 0.876] | 0.883 [0.878, 0.889] |
| | 30 | 0.844 [0.837, 0.850] | 0.886 [0.879, 0.893] | 0.809 [0.802, 0.816] | 0.892 [0.885, 0.898] | 0.876 [0.870, 0.883] | 0.890 [0.884, 0.897] |
| | 50 | 0.847 [0.839, 0.854] | 0.891 [0.883, 0.898] | 0.820 [0.813, 0.828] | 0.896 [0.888, 0.903] | 0.879 [0.872, 0.886] | 0.894 [0.887, 0.902] |
| | 100 | 0.856 [0.847, 0.864] | 0.898 [0.889, 0.906] | 0.833 [0.825, 0.842] | 0.904 [0.895, 0.913] | 0.886 [0.878, 0.895] | 0.904 [0.895, 0.913] |
| H211-cDNAxR10 | 1 | 0.586 [0.580, 0.592] | 0.514 [0.509, 0.519] | 0.275 [0.270, 0.280] | 0.551 [0.545, 0.556] | 0.441 [0.435, 0.446] | 0.620 [0.614, 0.625] |
| | 5 | 0.784 [0.775, 0.791] | 0.645 [0.638, 0.652] | 0.465 [0.457, 0.474] | 0.744 [0.736, 0.752] | 0.678 [0.670, 0.686] | 0.795 [0.787, 0.802] |
| | 10 | 0.827 [0.818, 0.837] | 0.690 [0.681, 0.699] | 0.582 [0.572, 0.592] | 0.802 [0.792, 0.811] | 0.761 [0.752, 0.771] | 0.847 [0.838, 0.856] |
| | 30 | 0.847 [0.834, 0.860] | 0.740 [0.728, 0.752] | 0.663 [0.650, 0.676] | 0.834 [0.822, 0.847] | 0.798 [0.786, 0.811] | 0.882 [0.870, 0.895] |
| | 50 | 0.855 [0.840, 0.870] | 0.731 [0.716, 0.745] | 0.693 [0.678, 0.708] | 0.841 [0.827, 0.856] | 0.804 [0.789, 0.819] | 0.891 [0.876, 0.905] |
| | 100 | 0.853 [0.834, 0.871] | 0.689 [0.671, 0.707] | 0.706 [0.688, 0.724] | 0.842 [0.824, 0.860] | 0.802 [0.783, 0.820] | 0.894 [0.876, 0.912] |
| H211-dRNA004 | 1 | 0.700 [0.694, 0.705] | 0.662 [0.656, 0.667] | 0.401 [0.396, 0.406] | 0.726 [0.721, 0.731] | 0.626 [0.621, 0.632] | 0.726 [0.721, 0.731] |
| | 5 | 0.846 [0.840, 0.853] | 0.786 [0.779, 0.792] | 0.552 [0.545, 0.559] | 0.849 [0.842, 0.855] | 0.809 [0.802, 0.815] | 0.840 [0.834, 0.846] |
| | 10 | 0.867 [0.860, 0.874] | 0.798 [0.791, 0.805] | 0.636 [0.629, 0.644] | 0.869 [0.862, 0.877] | 0.855 [0.848, 0.863] | 0.857 [0.850, 0.864] |
| | 30 | 0.886 [0.878, 0.895] | 0.789 [0.780, 0.798] | 0.710 [0.701, 0.719] | 0.888 [0.879, 0.896] | 0.873 [0.864, 0.882] | 0.880 [0.871, 0.888] |
| | 50 | 0.897 [0.887, 0.907] | 0.777 [0.767, 0.787] | 0.734 [0.724, 0.744] | 0.897 [0.887, 0.907] | 0.878 [0.868, 0.888] | 0.899 [0.890, 0.909] |
| | 100 | 0.904 [0.892, 0.916] | 0.730 [0.717, 0.742] | 0.754 [0.742, 0.766] | 0.903 [0.890, 0.914] | 0.874 [0.862, 0.886] | 0.908 [0.896, 0.920] |
| H526-MasSeq | 1 | 0.723 [0.719, 0.727] | 0.750 [0.746, 0.754] | 0.550 [0.546, 0.555] | 0.746 [0.742, 0.750] | 0.706 [0.702, 0.711] | 0.759 [0.755, 0.763] |
| | 5 | 0.808 [0.803, 0.813] | 0.842 [0.837, 0.846] | 0.662 [0.657, 0.667] | 0.837 [0.832, 0.841] | 0.825 [0.820, 0.830] | 0.849 [0.844, 0.853] |
| | 10 | 0.817 [0.812, 0.822] | 0.858 [0.852, 0.863] | 0.715 [0.710, 0.720] | 0.853 [0.848, 0.859] | 0.849 [0.844, 0.854] | 0.864 [0.858, 0.869] |
| | 30 | 0.822 [0.816, 0.828] | 0.873 [0.866, 0.879] | 0.759 [0.753, 0.765] | 0.870 [0.864, 0.877] | 0.861 [0.855, 0.867] | 0.879 [0.873, 0.885] |
| | 50 | 0.823 [0.816, 0.829] | 0.876 [0.869, 0.883] | 0.772 [0.765, 0.778] | 0.873 [0.867, 0.880] | 0.863 [0.856, 0.869] | 0.884 [0.877, 0.891] |
| | 100 | 0.829 [0.821, 0.836] | 0.884 [0.876, 0.892] | 0.791 [0.783, 0.798] | 0.881 [0.873, 0.889] | 0.870 [0.862, 0.877] | 0.893 [0.885, 0.900] |
| H526-cDNAxR10 | 1 | 0.557 [0.551, 0.562] | 0.471 [0.466, 0.475] | 0.238 [0.234, 0.243] | 0.525 [0.519, 0.530] | 0.410 [0.405, 0.415] | 0.595 [0.590, 0.600] |
| | 5 | 0.749 [0.741, 0.756] | 0.591 [0.585, 0.597] | 0.402 [0.395, 0.409] | 0.713 [0.706, 0.721] | 0.628 [0.620, 0.635] | 0.771 [0.764, 0.778] |
| | 10 | 0.787 [0.778, 0.796] | 0.637 [0.630, 0.645] | 0.499 [0.490, 0.508] | 0.769 [0.761, 0.778] | 0.707 [0.699, 0.716] | 0.821 [0.812, 0.829] |
| | 30 | 0.810 [0.798, 0.822] | 0.695 [0.684, 0.706] | 0.563 [0.551, 0.575] | 0.812 [0.800, 0.823] | 0.756 [0.744, 0.768] | 0.866 [0.854, 0.877] |
| | 50 | 0.811 [0.797, 0.824] | 0.681 [0.668, 0.694] | 0.575 [0.562, 0.589] | 0.815 [0.801, 0.828] | 0.757 [0.744, 0.771] | 0.870 [0.856, 0.883] |
| | 100 | 0.803 [0.786, 0.820] | 0.639 [0.622, 0.655] | 0.582 [0.565, 0.598] | 0.813 [0.797, 0.830] | 0.757 [0.740, 0.774] | 0.868 [0.851, 0.885] |
| H526-dRNA004 | 1 | 0.677 [0.672, 0.681] | 0.608 [0.604, 0.613] | 0.362 [0.358, 0.367] | 0.704 [0.700, 0.709] | 0.618 [0.613, 0.623] | 0.701 [0.696, 0.706] |
| | 5 | 0.810 [0.804, 0.815] | 0.719 [0.713, 0.724] | 0.484 [0.478, 0.489] | 0.823 [0.817, 0.829] | 0.780 [0.775, 0.786] | 0.811 [0.806, 0.817] |
| | 10 | 0.830 [0.824, 0.836] | 0.730 [0.724, 0.736] | 0.548 [0.541, 0.554] | 0.843 [0.836, 0.849] | 0.822 [0.815, 0.828] | 0.830 [0.824, 0.836] |
| | 30 | 0.849 [0.842, 0.856] | 0.724 [0.717, 0.732] | 0.596 [0.589, 0.604] | 0.862 [0.855, 0.870] | 0.833 [0.825, 0.840] | 0.854 [0.847, 0.861] |
| | 50 | 0.858 [0.850, 0.867] | 0.710 [0.702, 0.718] | 0.615 [0.606, 0.623] | 0.869 [0.861, 0.877] | 0.835 [0.826, 0.843] | 0.867 [0.859, 0.875] |
| | 100 | 0.866 [0.857, 0.876] | 0.672 [0.663, 0.682] | 0.629 [0.620, 0.639] | 0.877 [0.868, 0.887] | 0.828 [0.818, 0.838] | 0.882 [0.872, 0.891] |

**Table S14: SNV F1 score with 95% credible intervals, by sample and coverage cutoff.** F1 = 2*TP / (TRUTH.TOTAL + QUERY.TOTAL); 95% credible intervals computed per sample and coverage cutoff via a Jeffreys-style Beta(TP + 0.5, N - TP + 0.5) posterior on the underlying proportion, doubled to the F1 scale. Unlike the pooled summary table, these intervals are per-sample and therefore wider.

| Dataset | Coverage | Clair3-RNA | DeepVariant | isoLASER |
|---|---|---|---|---|
| H211-MasSeq | 1 | 0.524 [0.515, 0.535] | 0.567 [0.557, 0.577] | 0.190 [0.182, 0.197] |
| | 5 | 0.604 [0.594, 0.617] | 0.635 [0.624, 0.647] | 0.241 [0.232, 0.251] |
| | 10 | 0.617 [0.606, 0.631] | 0.651 [0.638, 0.664] | 0.278 [0.267, 0.289] |
| | 30 | 0.630 [0.617, 0.647] | 0.671 [0.655, 0.686] | 0.332 [0.319, 0.347] |
| | 50 | 0.631 [0.616, 0.650] | 0.673 [0.656, 0.690] | 0.358 [0.343, 0.374] |
| | 100 | 0.634 [0.616, 0.655] | 0.678 [0.657, 0.697] | 0.385 [0.367, 0.405] |
| H211-cDNAxR10 | 1 | 0.317 [0.306, 0.329] | 0.315 [0.305, 0.327] | – |
| | 5 | 0.481 [0.464, 0.498] | 0.465 [0.448, 0.482] | – |
| | 10 | 0.514 [0.494, 0.535] | 0.508 [0.488, 0.529] | – |
| | 30 | 0.531 [0.503, 0.559] | 0.530 [0.502, 0.560] | – |
| | 50 | 0.540 [0.508, 0.573] | 0.525 [0.491, 0.559] | – |
| | 100 | 0.553 [0.512, 0.594] | 0.506 [0.463, 0.549] | – |
| H211-dRNA004 | 1 | 0.262 [0.252, 0.273] | 0.278 [0.269, 0.288] | – |
| | 5 | 0.387 [0.372, 0.403] | 0.386 [0.372, 0.401] | – |
| | 10 | 0.405 [0.387, 0.423] | 0.409 [0.392, 0.426] | – |
| | 30 | 0.387 [0.364, 0.411] | 0.382 [0.360, 0.404] | – |
| | 50 | 0.380 [0.354, 0.408] | 0.358 [0.334, 0.383] | – |
| | 100 | 0.375 [0.342, 0.409] | 0.308 [0.280, 0.335] | – |
| H526-MasSeq | 1 | 0.509 [0.500, 0.520] | 0.563 [0.554, 0.574] | 0.202 [0.194, 0.210] |
| | 5 | 0.586 [0.577, 0.599] | 0.634 [0.622, 0.645] | 0.253 [0.244, 0.263] |
| | 10 | 0.602 [0.591, 0.615] | 0.651 [0.639, 0.663] | 0.287 [0.277, 0.299] |
| | 30 | 0.616 [0.604, 0.632] | 0.672 [0.657, 0.686] | 0.336 [0.324, 0.350] |
| | 50 | 0.622 [0.608, 0.639] | 0.679 [0.662, 0.694] | 0.358 [0.344, 0.374] |
| | 100 | 0.631 [0.614, 0.650] | 0.687 [0.668, 0.705] | 0.384 [0.367, 0.402] |
| H526-cDNAxR10 | 1 | 0.312 [0.301, 0.323] | 0.322 [0.311, 0.332] | – |
| | 5 | 0.470 [0.454, 0.487] | 0.472 [0.455, 0.487] | – |
| | 10 | 0.501 [0.483, 0.521] | 0.515 [0.494, 0.533] | – |
| | 30 | 0.528 [0.503, 0.555] | 0.559 [0.531, 0.586] | – |
| | 50 | 0.538 [0.508, 0.569] | 0.561 [0.527, 0.592] | – |
| | 100 | 0.541 [0.503, 0.581] | 0.534 [0.492, 0.575] | – |
| H526-dRNA004 | 1 | 0.372 [0.362, 0.383] | 0.337 [0.327, 0.345] | – |
| | 5 | 0.465 [0.452, 0.478] | 0.399 [0.388, 0.409] | – |
| | 10 | 0.482 [0.469, 0.496] | 0.409 [0.396, 0.420] | – |
| | 30 | 0.491 [0.475, 0.508] | 0.408 [0.394, 0.422] | – |
| | 50 | 0.493 [0.476, 0.512] | 0.406 [0.391, 0.421] | – |
| | 100 | 0.495 [0.475, 0.516] | 0.390 [0.373, 0.408] | – |

**Table S15: INDEL F1 score with 95% credible intervals, by sample and coverage cutoff.** F1 = 2*TP / (TRUTH.TOTAL + QUERY.TOTAL); 95% credible intervals computed per sample and coverage cutoff via a Jeffreys-style Beta(TP + 0.5, N - TP + 0.5) posterior on the underlying proportion, doubled to the F1 scale. longcallR and longcallR-nn produce no usable INDEL calls and are omitted. isoLASER INDEL calls are only usable on the PacBio platform (MasSeq) and are left blank (–) elsewhere.

### 2 Supplementary Figures

#### 2.1 Overall variant calling performance

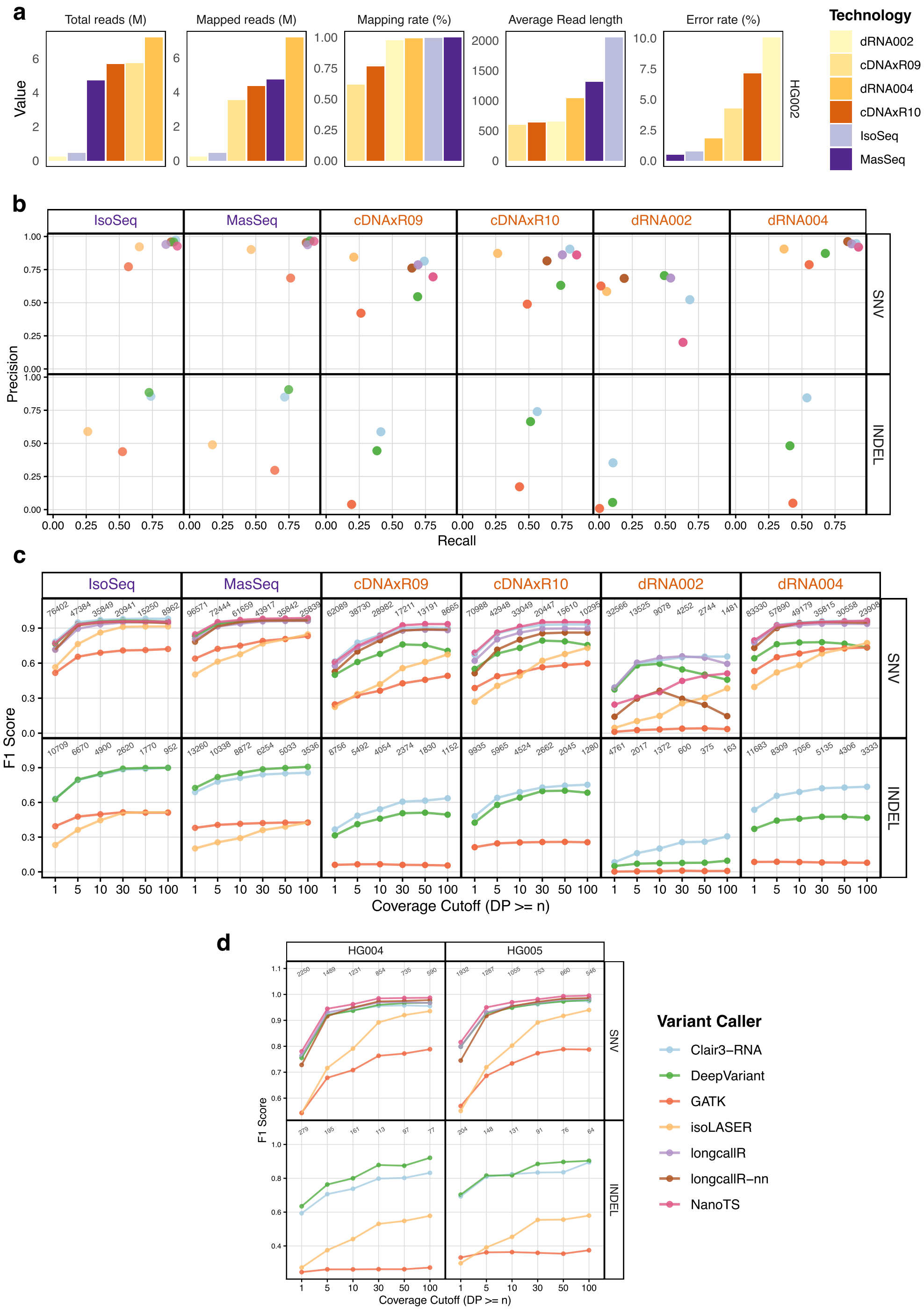

**Fig. S1: Data quality and variant-calling performance for HG002 and chromosome 20-only datasets.** (**a**) Data quality metrics for the HG002 datasets, colored by library preparation kit and stratified by metric and sequencing chemistry. Total sequences and mapped reads are shown in millions. (**b**) Precision and recall of each variant caller at 5× coverage, stratified by variant type (SNVs and indels) and library preparation kit. (**c**) F1 scores for SNV and indel calling across different minimum read-depth thresholds. (**d**) F1 scores for SNV and indel calling in chromosome 20-only HG004 and HG005 Mas-Seq datasets. All variant calls were generated from minimap2 alignments without BAM transformation. Panels **b**–**d** share the same color legend.

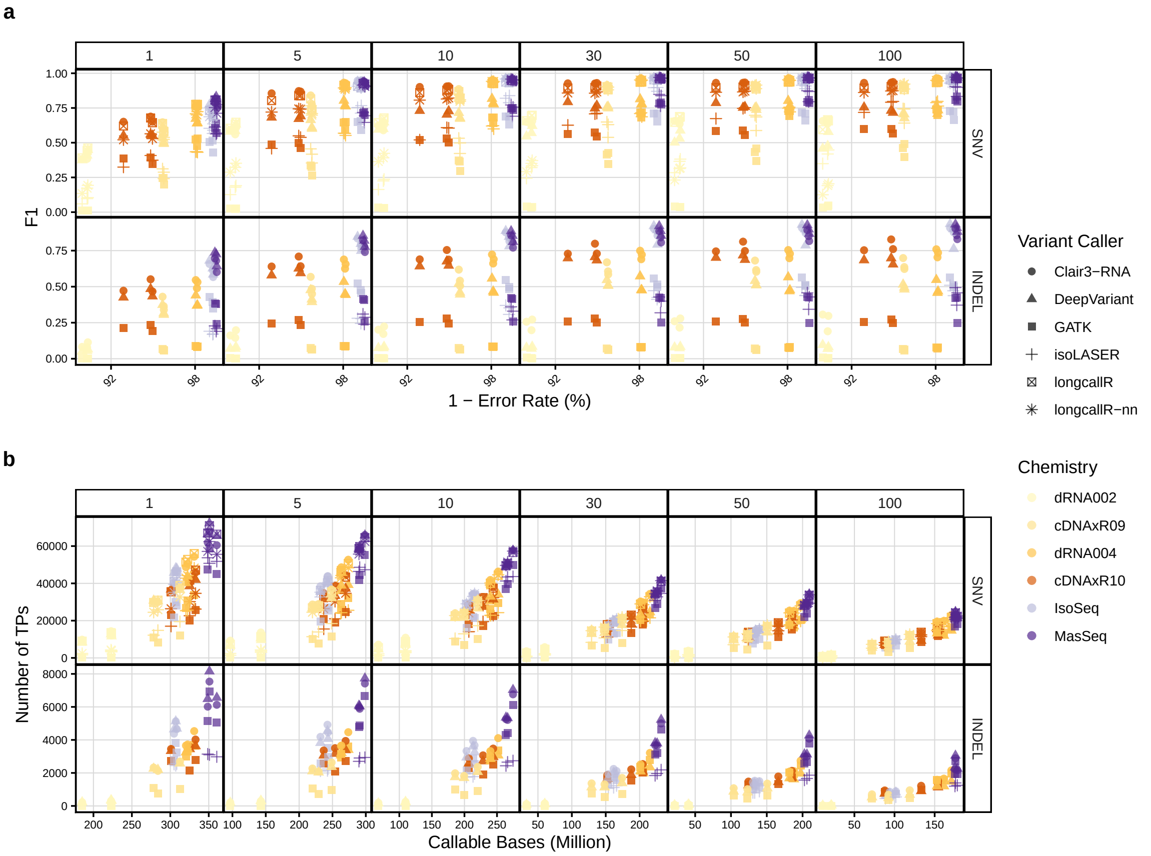

**Fig. S2: Impact of sequencing characteristics on variant calling performance in the GIAB datasets.** (**a**) Alignment-derived error rate versus F1 scores for each variant caller. (**b**) Number of callable bases versus the number of TP variant calls returned by each variant caller. Rows = SNVs and INDELs.

**a**

Standard evaluation region

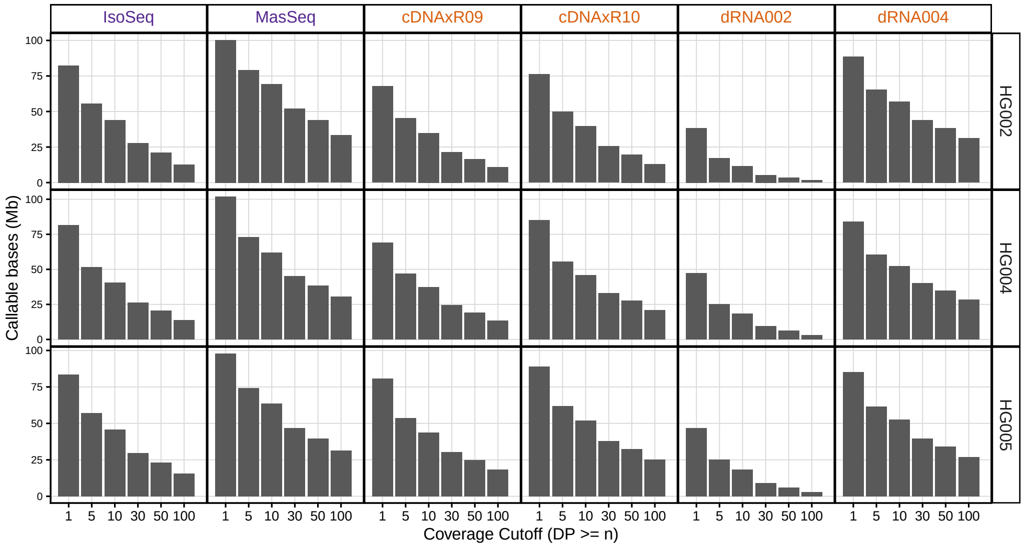

**b**

Common region (standard region with isoLASER effective depth)

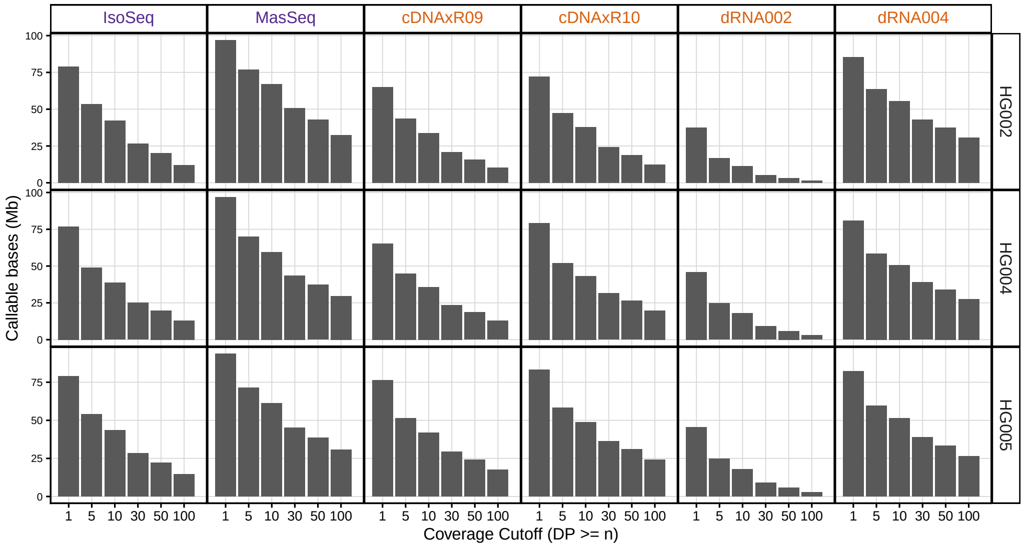

**Fig. S3: Number of callable bases in each GIAB dataset at different minimum read-depth thresholds.** (**a**) Callable regions are defined as GENCODE-annotated exonic regions that are covered by mapped lrRNA-seq reads at the corresponding minimum read-depth threshold and intersect the GIAB high-confidence regions. (**b**) The same as panel **a**, but intersected with isoLASER-preprocessed reads.

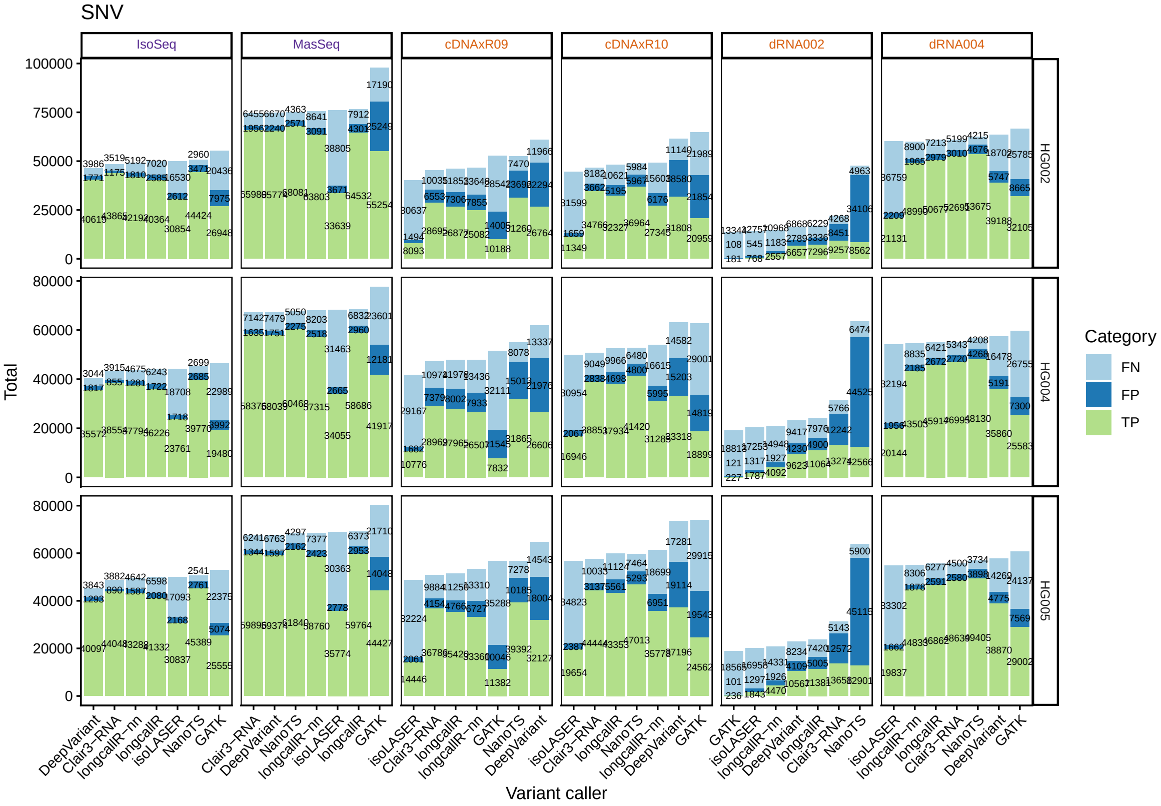

**Fig. S4: The number of FP, FN, and TP SNV calls by different variant callers at cutoff 5.** Annotated numbers from the top to the bottom indicate the number of FN, FP and TP.

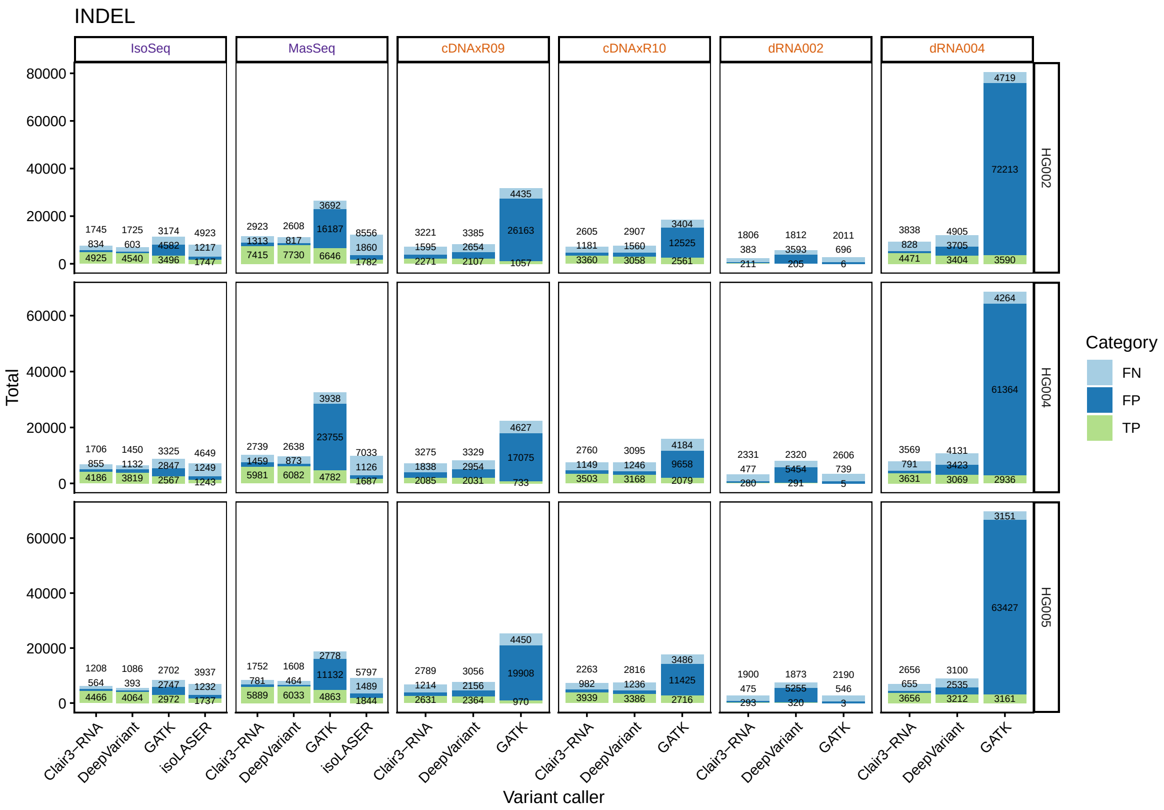

**Fig. S5: The number of FP, FN, and TP indel calls by different variant callers at cutoff 5.** Annotated numbers from the top to the bottom indicate the number of FN, FP and TP.

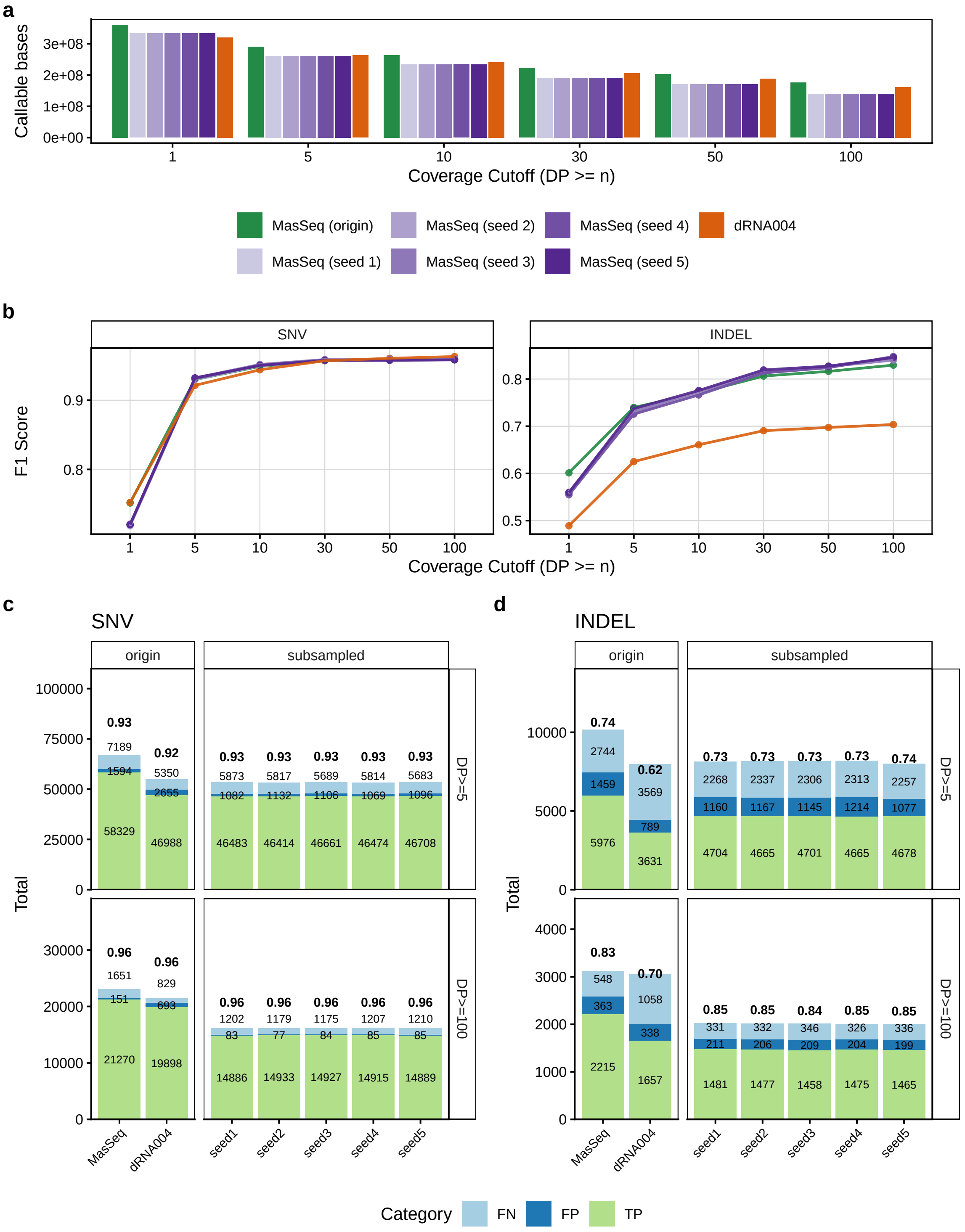

**Fig. S6: Subsampling analysis of variant-calling performance using HG004 datasets.** (**a**) The number of callable bases in the original and subsampled Mas-Seq datasets (using five random seeds) and the original dRNA004 dataset for the HG004 cell line. (**b**) Variant-calling performance measured by F1 score for the original and subsampled Mas-Seq datasets and the original dRNA004 dataset using Clair3-RNA. (**c, d**) The numbers of TP, FP, and FN calls for (**c**) SNVs and (**d**) indels in the original and subsampled Mas-Seq datasets and the original dRNA004 dataset at coverage cutoffs of 5 and 100, annotated with F1 scores, the number of FN, FP and TP. Clair3-RNA was run with the default parameters.

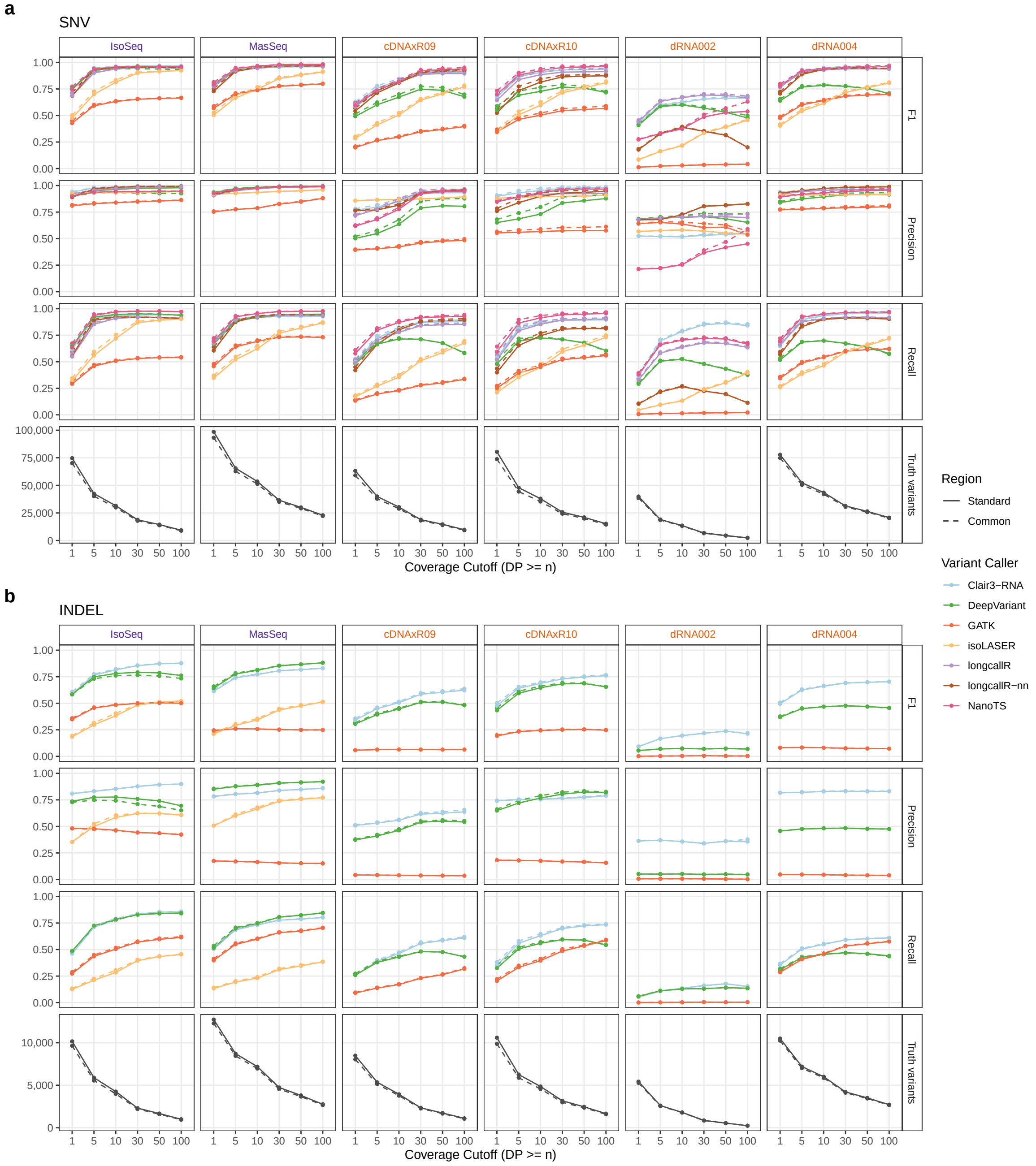

**Fig. S7: Variant-calling performance on regions covered after isoLASER preprocessing.** Variant calling performance for SNVs (**a**) and indels (**b**) evaluated on the common regions of the genome-aligned reads and the isoLASER-preprocessed reads. Truth variants indicate the number of true variants in ground truth set in the benchmarking region.

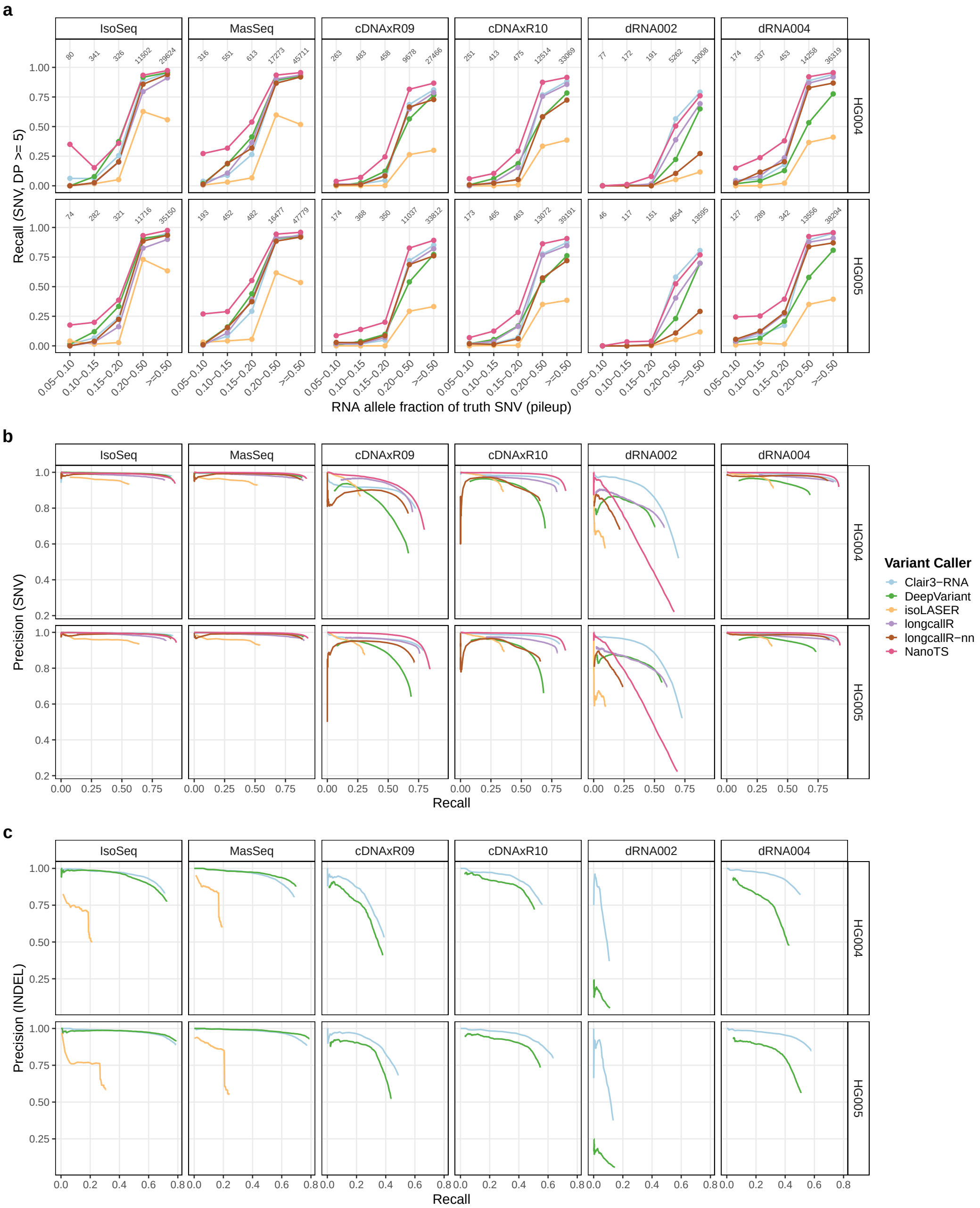

**Fig. S8: Robustness of caller performance to allele-fraction and quality-score thresholds.** (**a**) SNV recall stratified by the RNA allele fraction of each truth SNV, measured by pileup of the genome-aligned reads (MAPQ $\geq 20$); numbers indicate the truth SNVs per bin. (**b**, **c**) Precision–recall curves for SNVs (**b**) and indels (**c**) obtained by sweeping each caller's QUAL score over its PASS calls, so curves extend only to thresholds stricter than each caller's default filter; RNA-editing-labelled calls are therefore excluded as in the main analysis. The irregular curves of longcallR-nn reflect its QUAL distribution: for PacBio and ONT cDNA data, 50–82% of its SNV calls have QUAL 16–20 and at most 4% exceed 25, and these high-QUAL calls have lower precision than the main group, so its curves dip at stringent thresholds, most visibly for ONT cDNA. Indel curves are shown for Clair3-RNA, DeepVariant and isoLASER (PacBio data only). Rows = cell lines, columns = library preparation kits; minimap2 alignments, coverage $\geq 5$.

#### 2.2 pbmm2 vs. minimap2

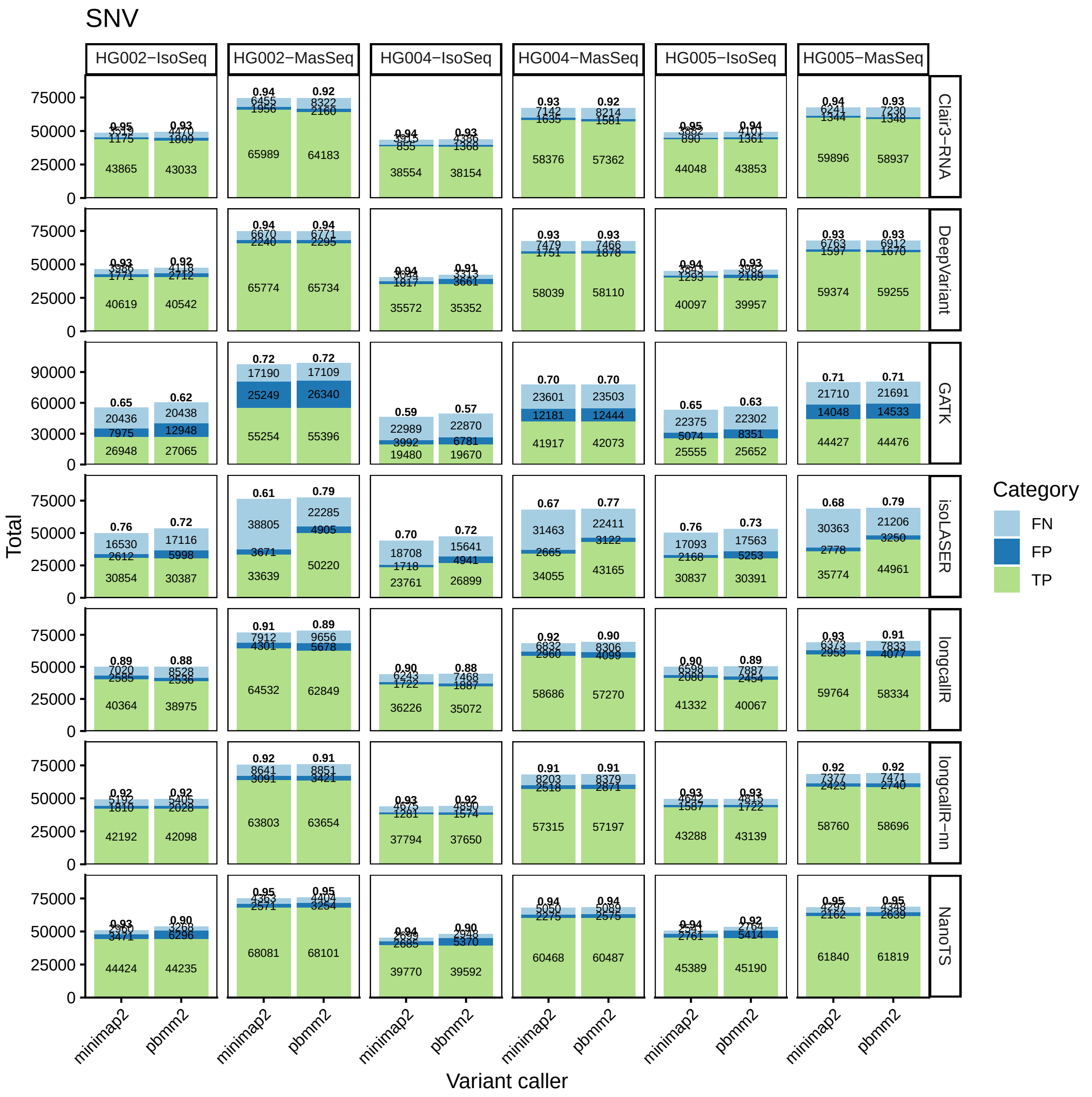

**Fig. S9: The number of FP, FN, and TP SNV calls by pbmm2 and minimap2 at cutoff 5 in PacBio datasets.** Annotated numbers from the top to the bottom indicate F1 score and the number of FN, FP and TP.

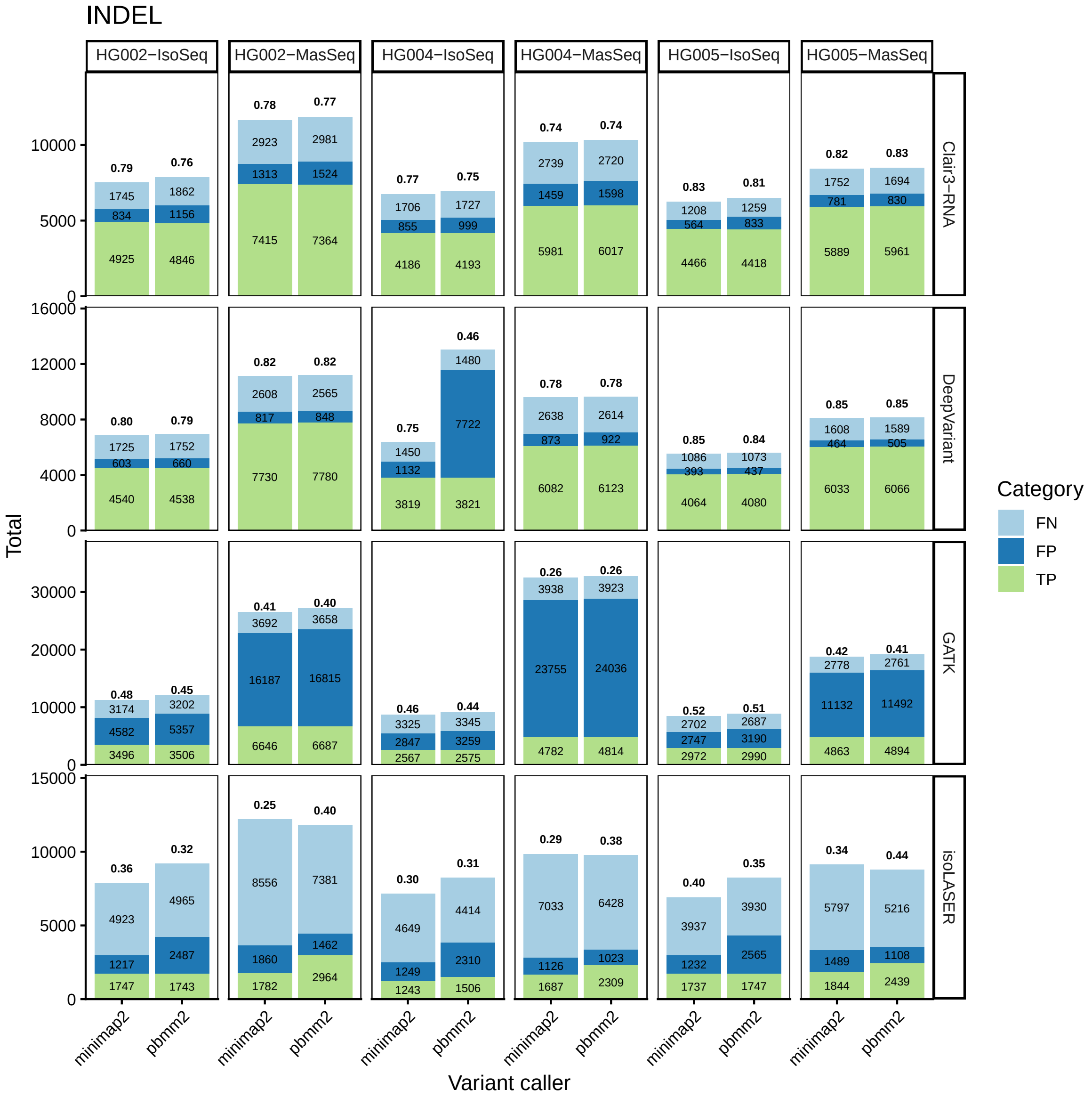

**Fig. S10: The number of FP, FN, and TP indel calls by pbmm2 and minimap2 at cutoff 5 in PacBio datasets.** Annotated numbers from the top to the bottom indicate F1 score and the number of FN, FP and TP.

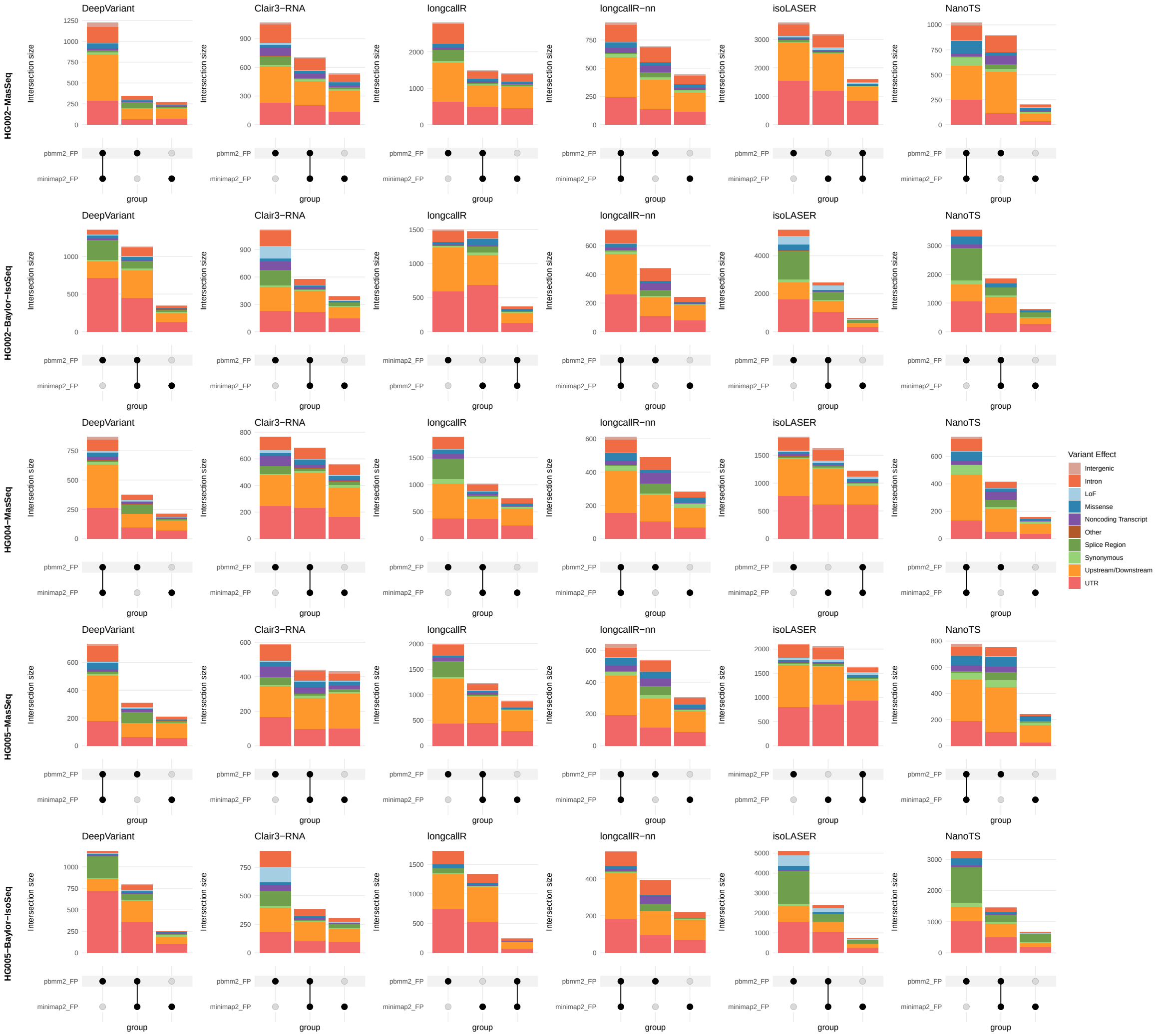

**Fig. S11: UpSet plots illustrating the genomic context of aligner-specific and shared FP calls at coverage cutoff of 5x.**

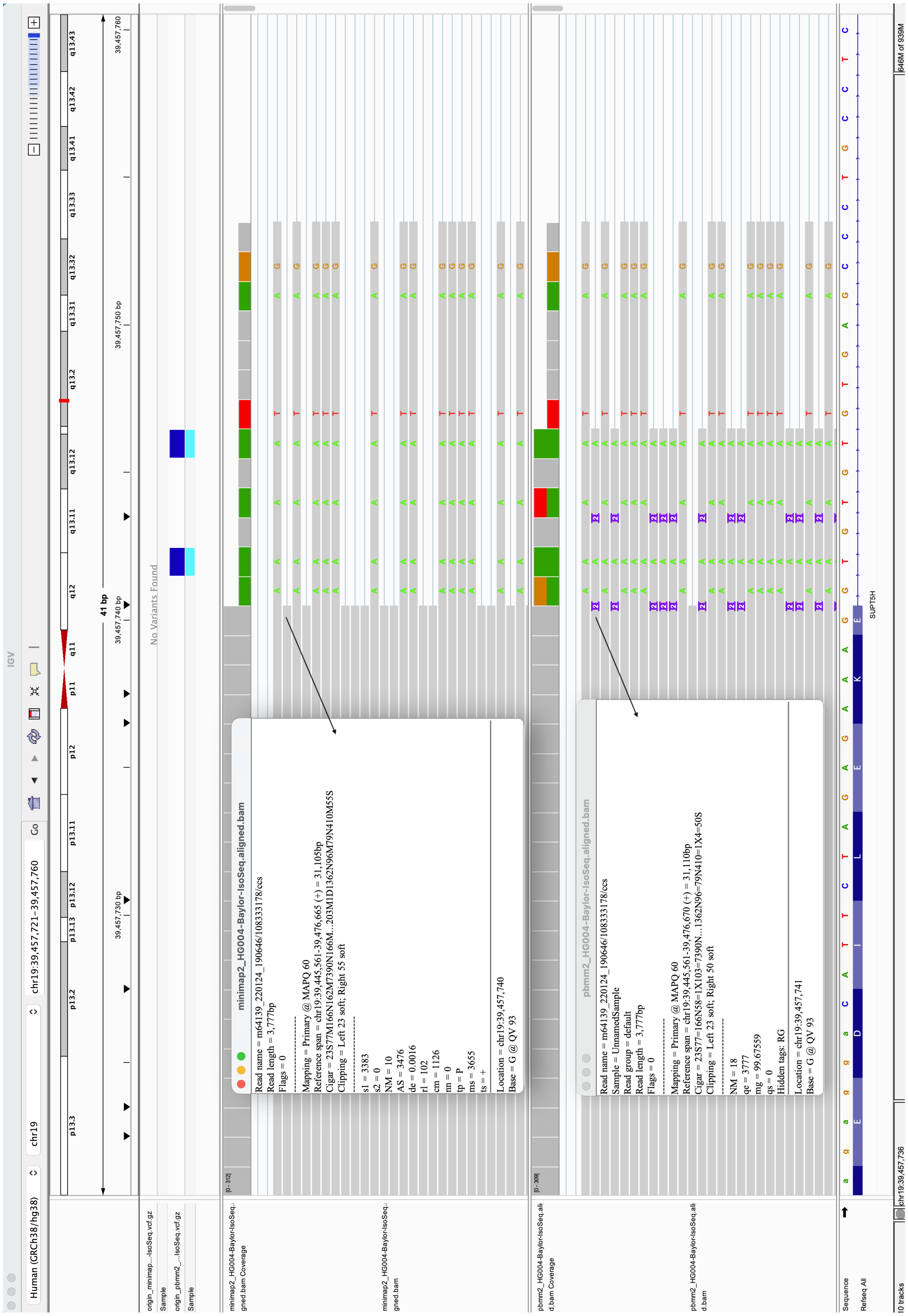

**Fig. S12: IGV screenshot of read alignments generated by pbmm2 and minimap2.** Alignments at a splice-region pbmm2-specific FP SNV (chr19:39,457,741_G/A) detected by longcallR. The window displays detailed information for a representative pbmm2 and minimap2 read.

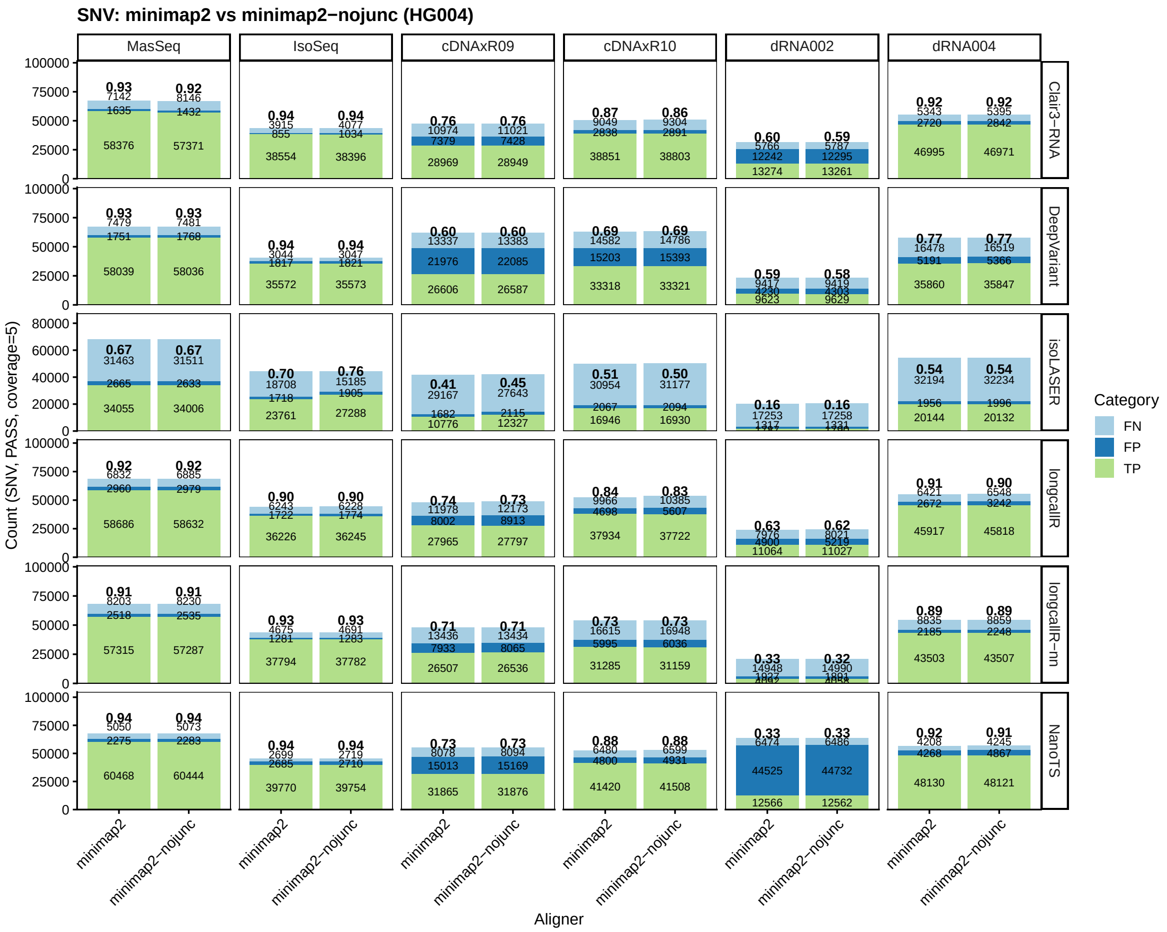

**Fig. S13: The number of FPs, FNs, and TPs SNV calls by minimap2 with and without the transcriptome annotation file as input at cutoff 5 in HG004 datasets.** Annotated numbers from the top to the bottom indicate F1 score and the number of FN, FP and TP. minimap2-nojunc means minimap2 alignment without transcriptome annotation BED file passing for `--junc-bed`

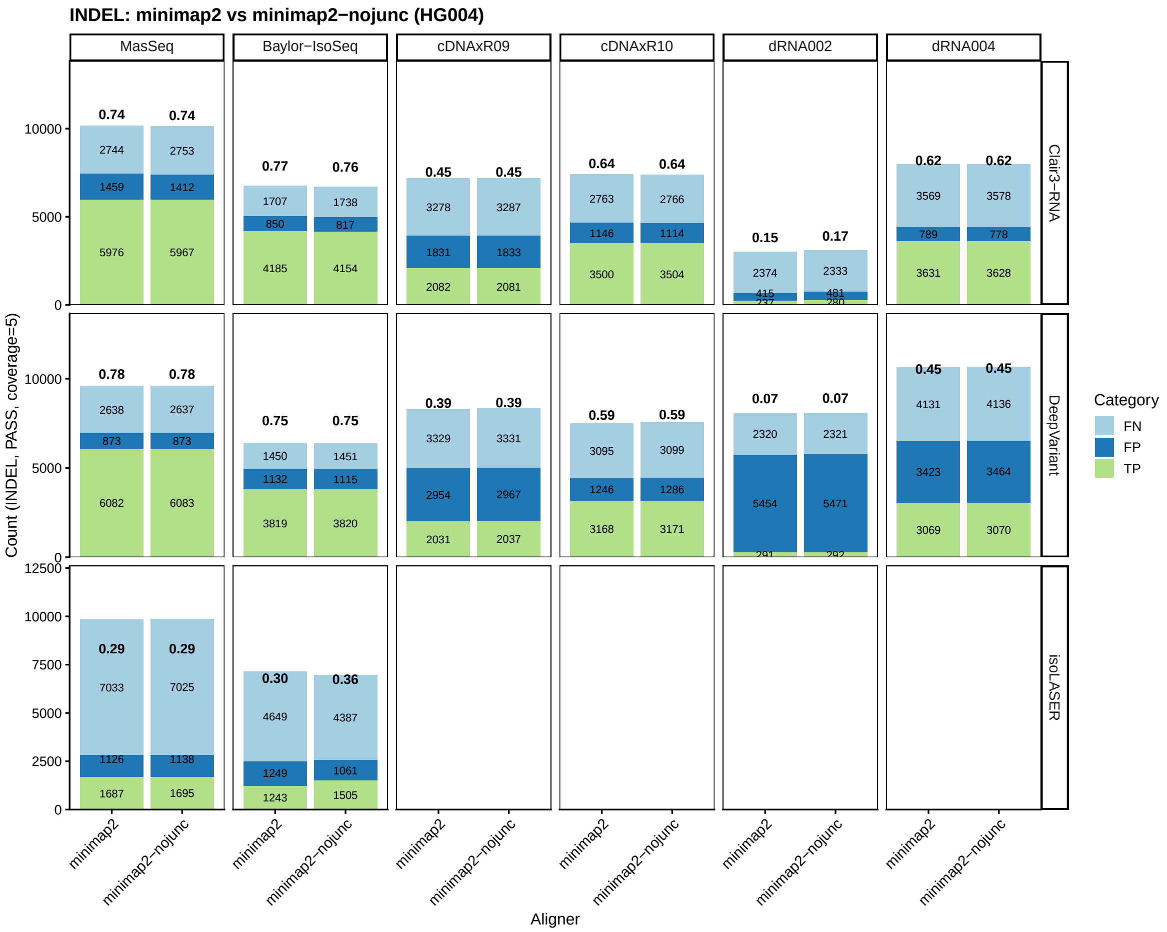

**Fig. S14: The number of FPs, FNs, and TPs indel calls by minimap2 with and without the transcriptome annotation file as input at cutoff 5 in HG004 datasets.** Annotated numbers from the top to the bottom indicate F1 score and the number of FN, FP and TP. minimap2-nojunc means minimap2 alignment without transcriptome annotation BED file passing for `--junc-bed`

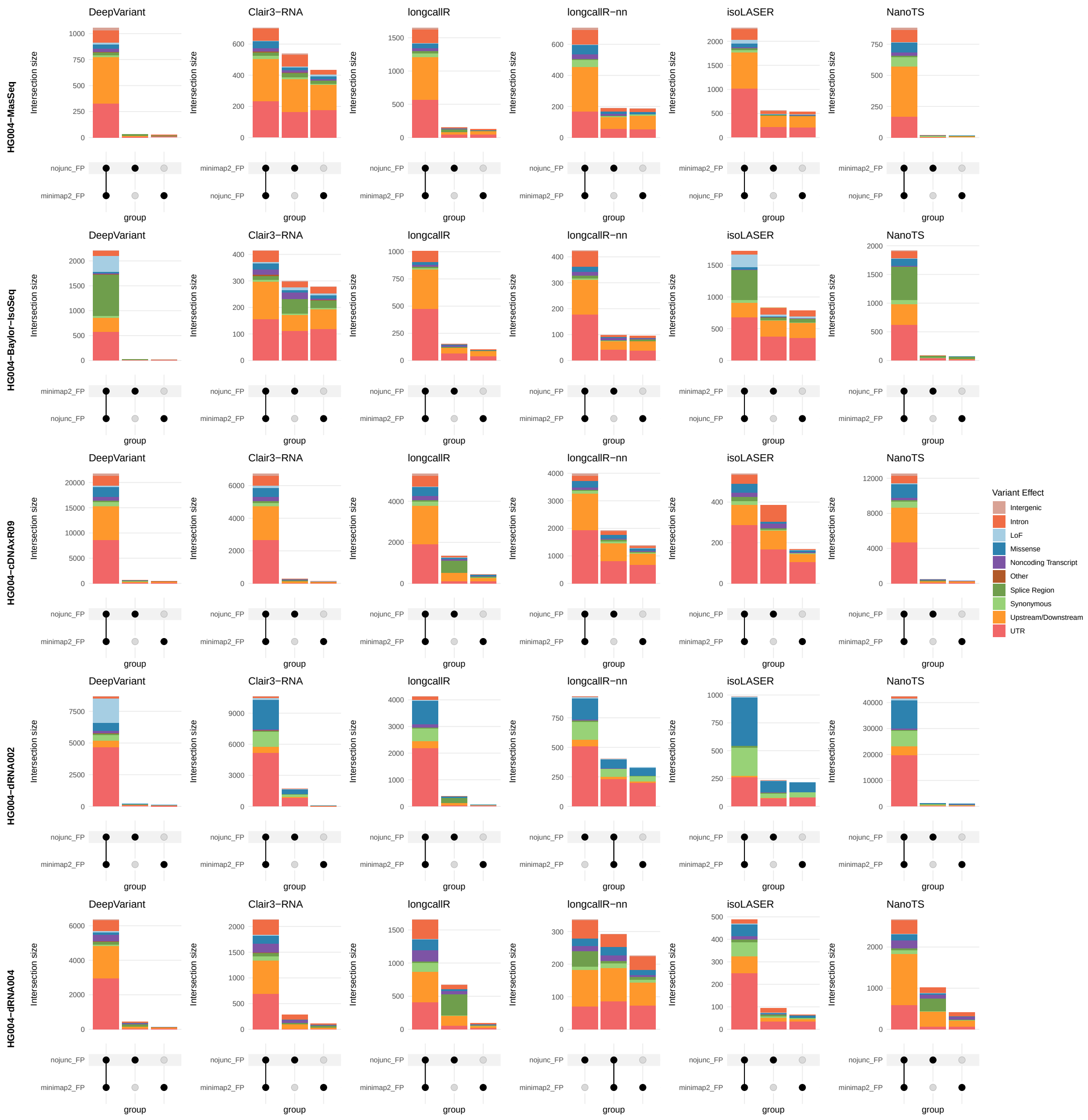

**Fig. S15: UpSet plots illustrating the genomic context of aligner-specific and shared FP calls at cutoff 5x.** nojunc means minimap2 alignment without transcriptome annotation BED file passing for `--junc-bed`

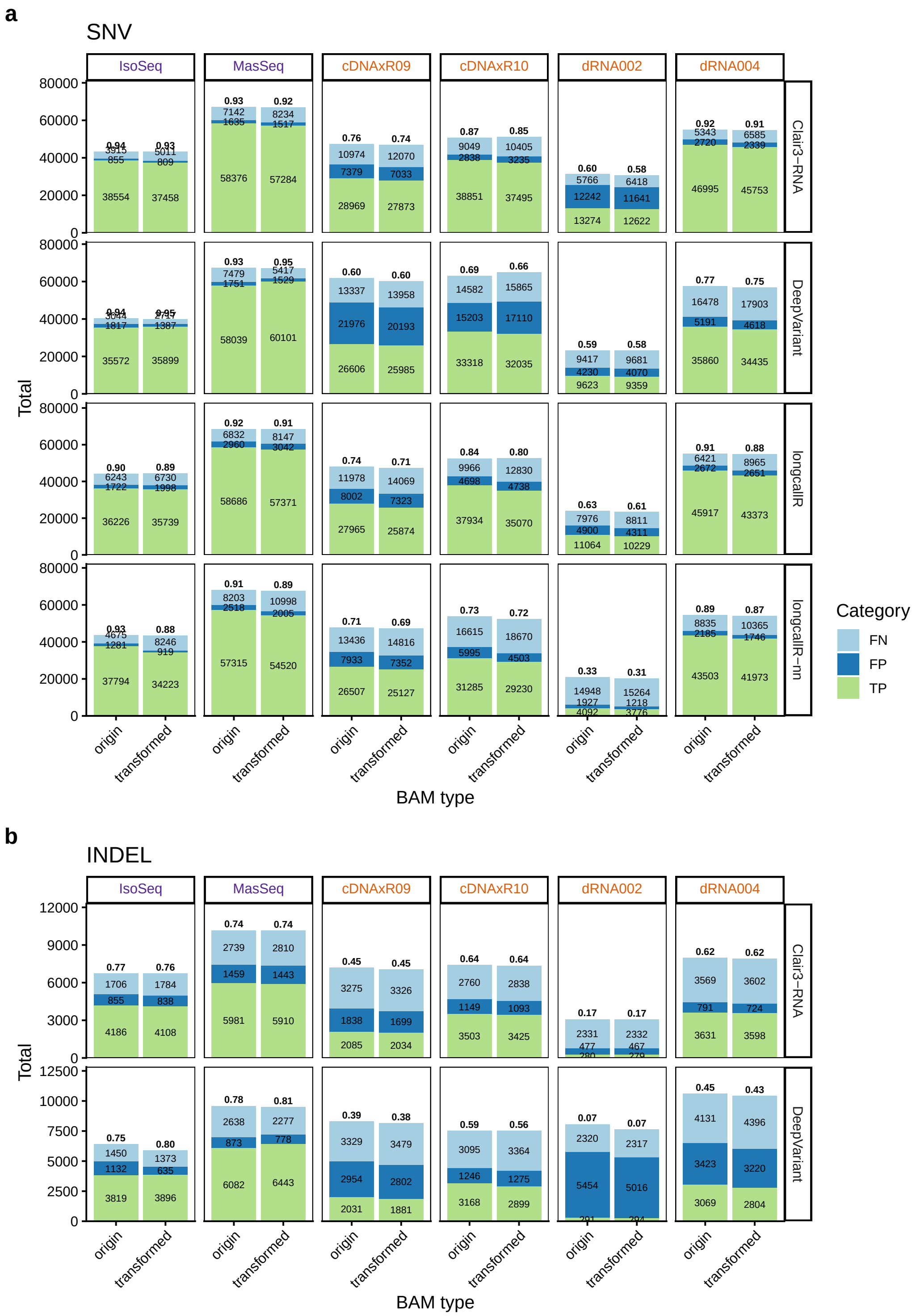

**Fig. S16: The number of FP, FN, and TP calls from raw genome-aligned and transformed reads at cutoff 5 in HG004 datasets.** Annotated numbers from the top to the bottom indicate F1 score and the number of FN, FP and TP for (**a**) SNVs and (**b**) indels.

#### 2.3 RNA editing site overlap

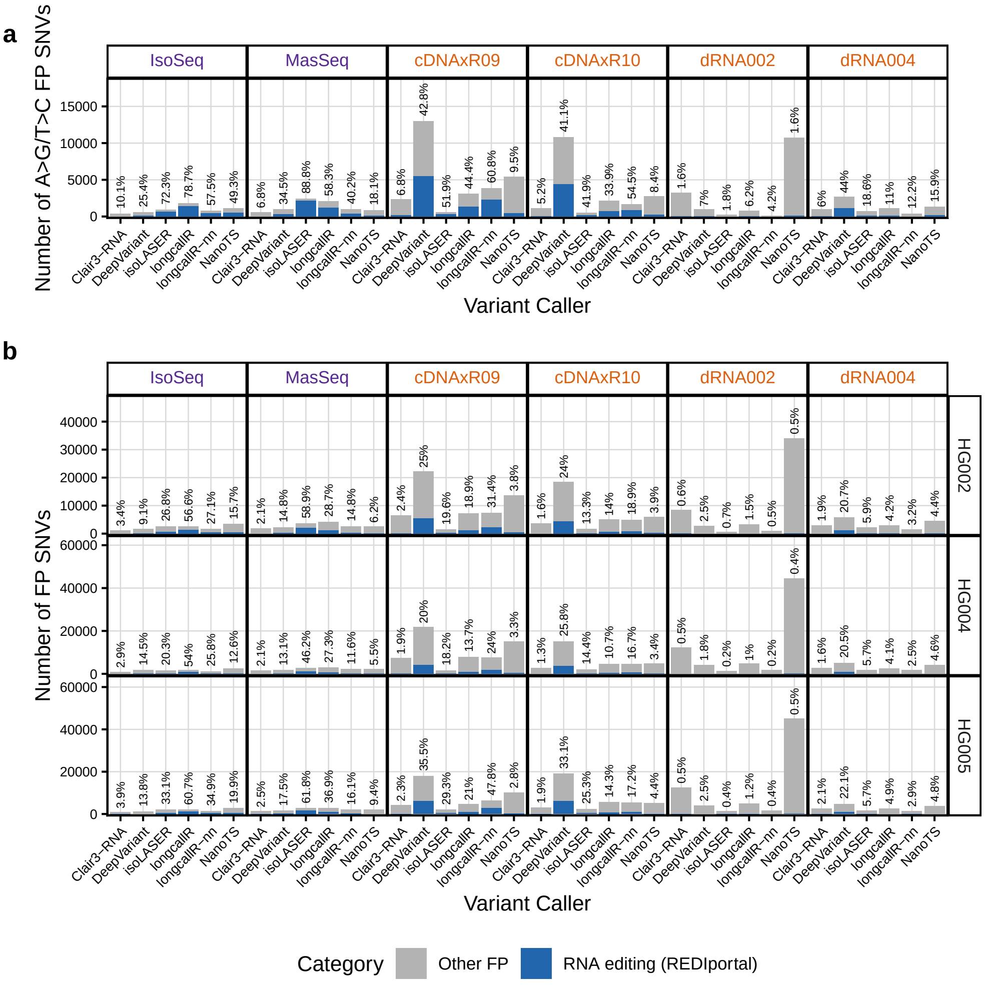

**Fig. S17: Proportion of RNA editing sites in FP SNVs.** (**a**) Number of A-to-G/T-to-C FP SNVs overlapping REDIportal (blue) or not (grey) for HG002 datasets. Percentages of REDIportal-overlapping SNVs are shown above each bar. (**b**) Number of all FP SNVs overlapping REDIportal (blue) or not (grey). Percentages of REDIportal-overlapping SNVs are shown above each bar.

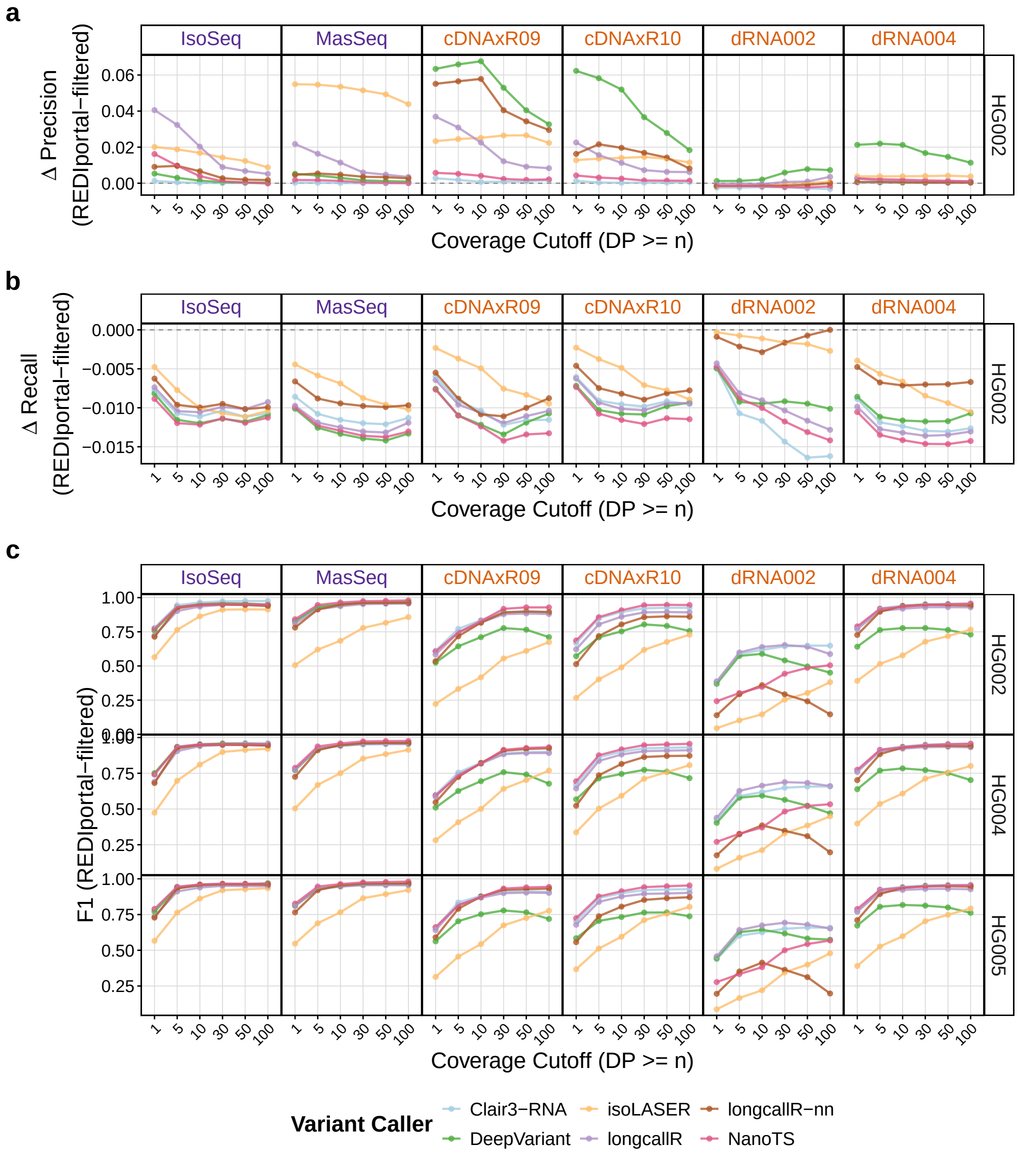

**Fig. S18: Proportion of RNA editing sites in FP SNVs.** (**a, b**) Change in precision (**a**) and recall (**b**) after excluding REDIportal-overlapping SNVs for HG002 datasets. (**c**) F1 scores after excluding REDIportal-overlapping SNVs.

#### 2.4 Genome-context-stratified regions

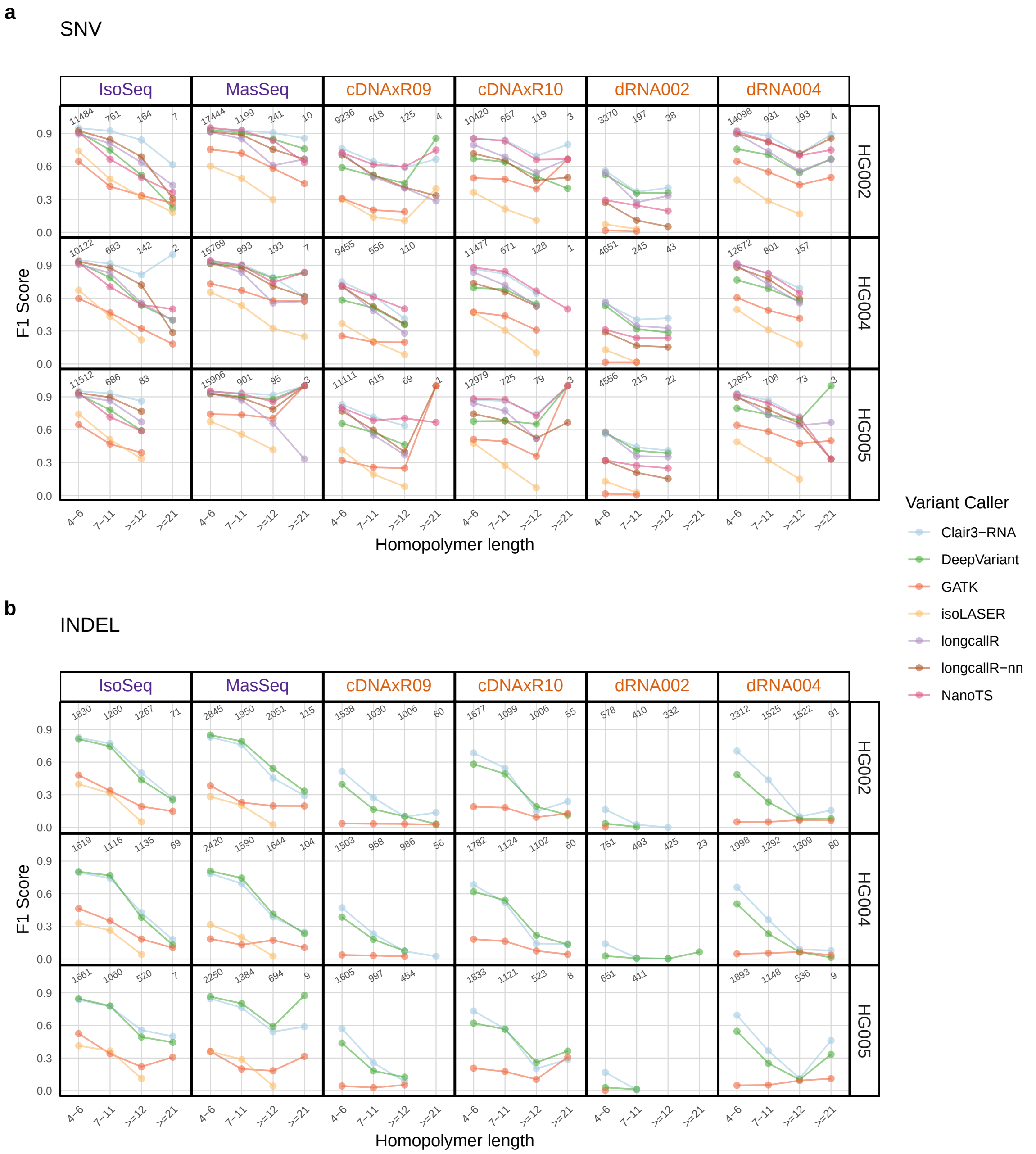

**Fig. S19: Variant calling performance based on F1 scores across different homopolymer lengths.** F1 scores for SNV (**a**) and INDEL (**b**) calling within homopolymer regions. Rows = cell lines, columns = library preparation kits. The annotated number indicates the number of variants in truth set.

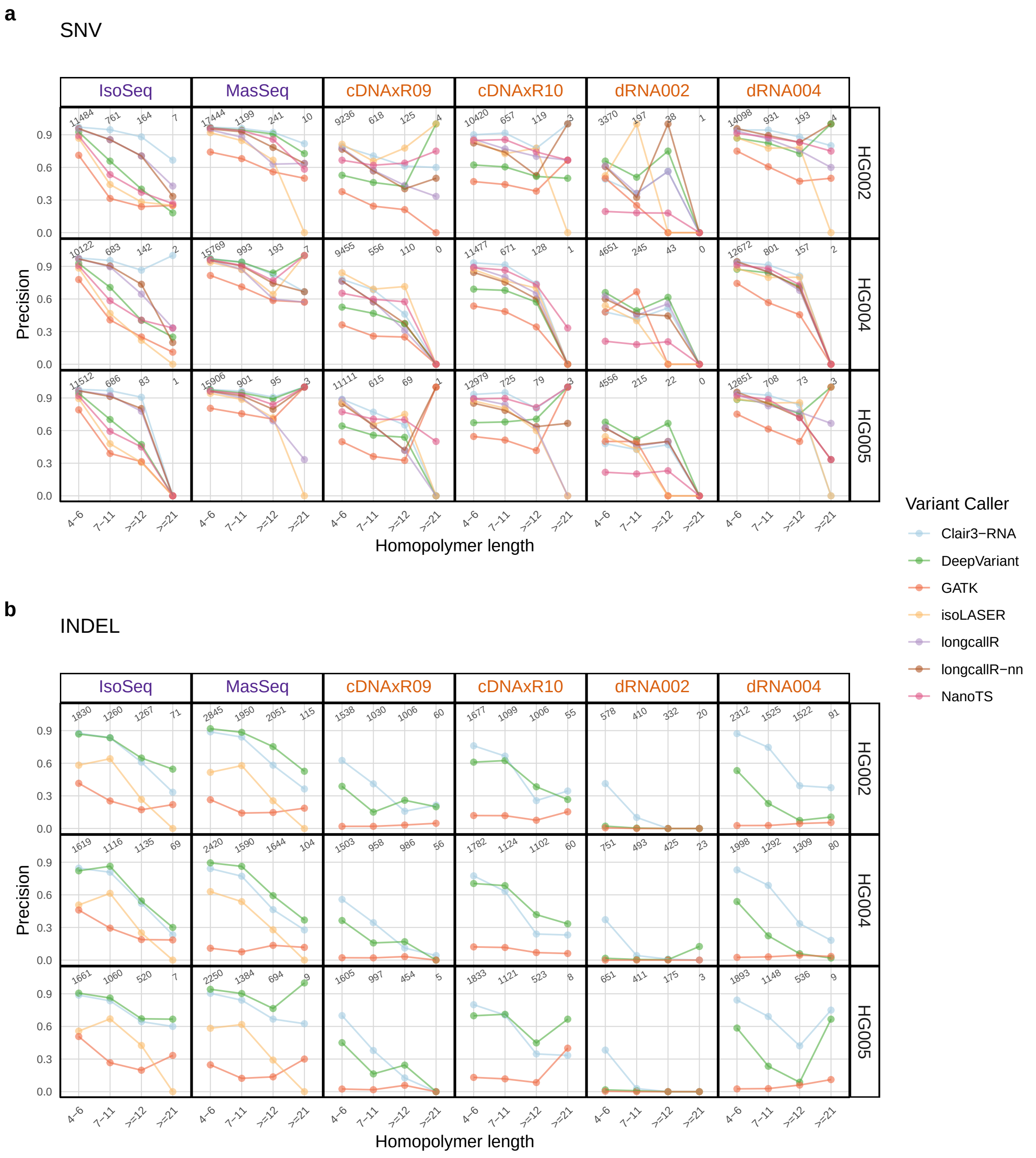

**Fig. S20: Variant calling performance based on precision across different homopolymer lengths.** Precision for SNV (**a**) and INDEL (**b**) calling within homopolymer regions. Rows = cell lines, columns = library preparation kits.

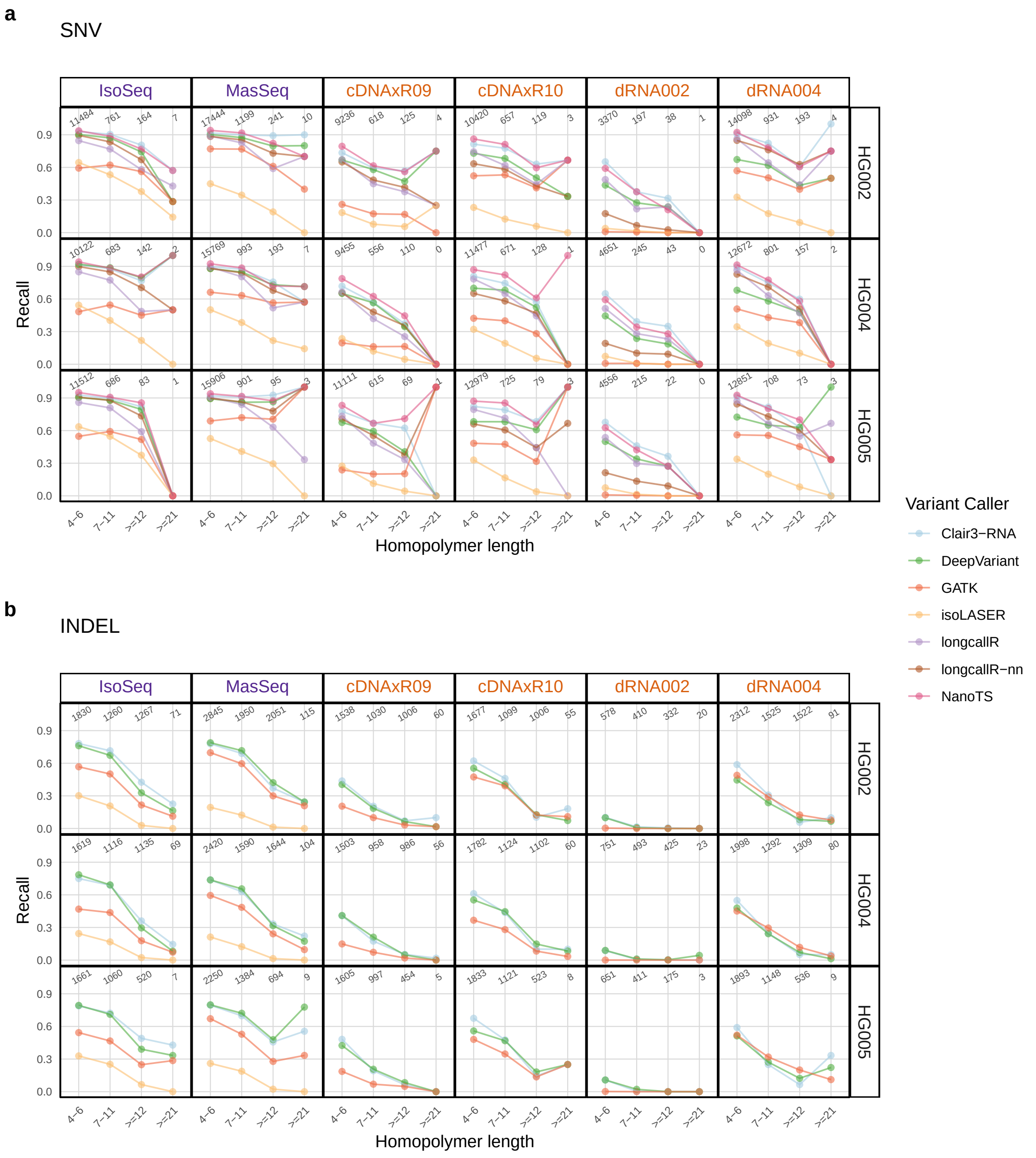

**Fig. S21: Variant calling performance based on recall across different homopolymer lengths.** Recall for SNV (**a**) and INDEL (**b**) calling within homopolymer regions. Rows = cell lines, columns = library preparation kits.

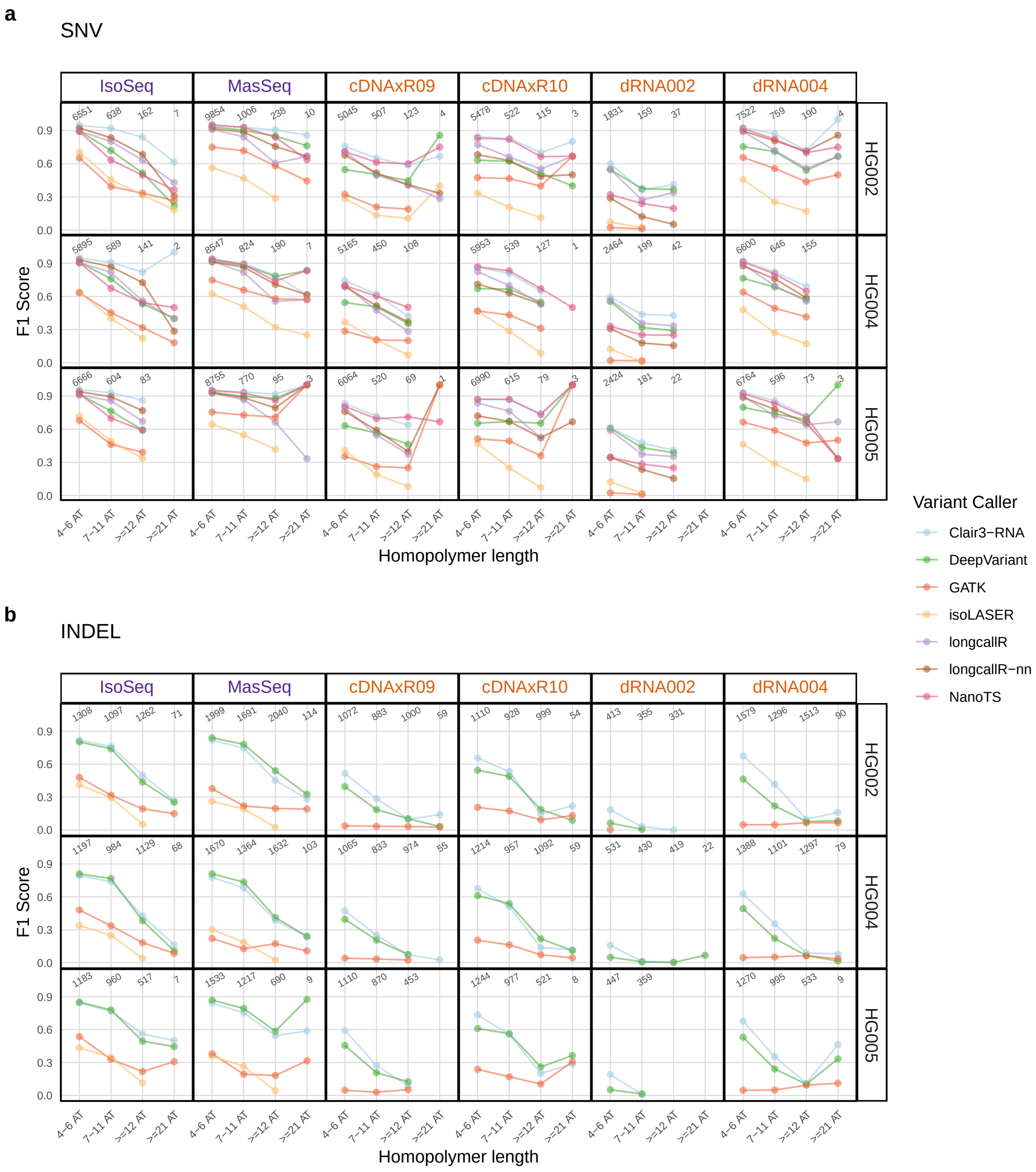

**Fig. S22: Variant calling performance based on F1 scores across different homopolymer lengths (AT specific).** F1 scores for SNV (**a**) and INDEL (**b**) calling within homopolymer regions. Rows = cell lines, columns = library preparation kits.

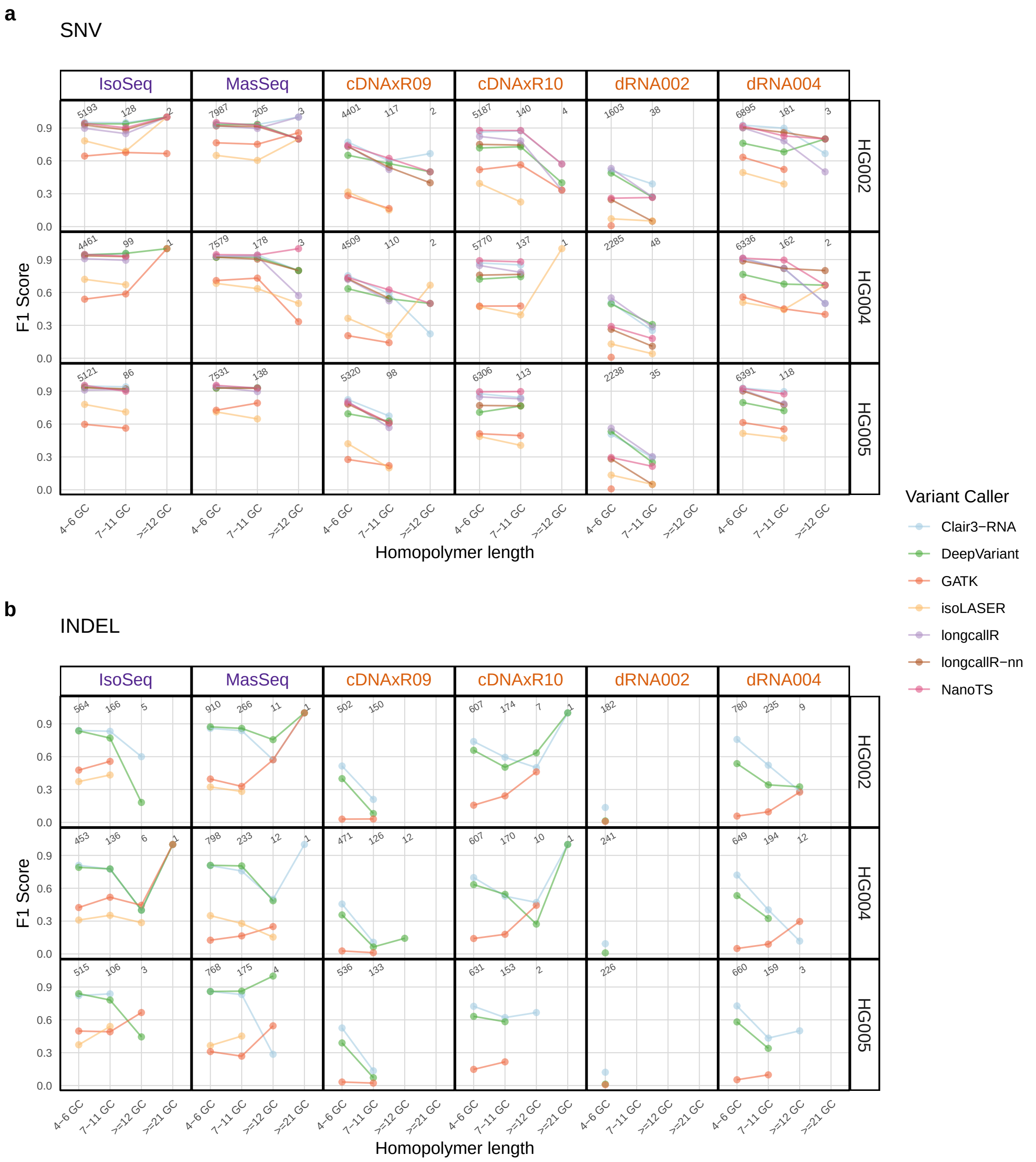

**Fig. S23: Variant calling performance based on F1 scores across different homopolymer lengths (GC specific).** F1 scores for SNV (**a**) and INDEL (**b**) calling within homopolymer regions. Rows = cell lines, columns = library preparation kits.

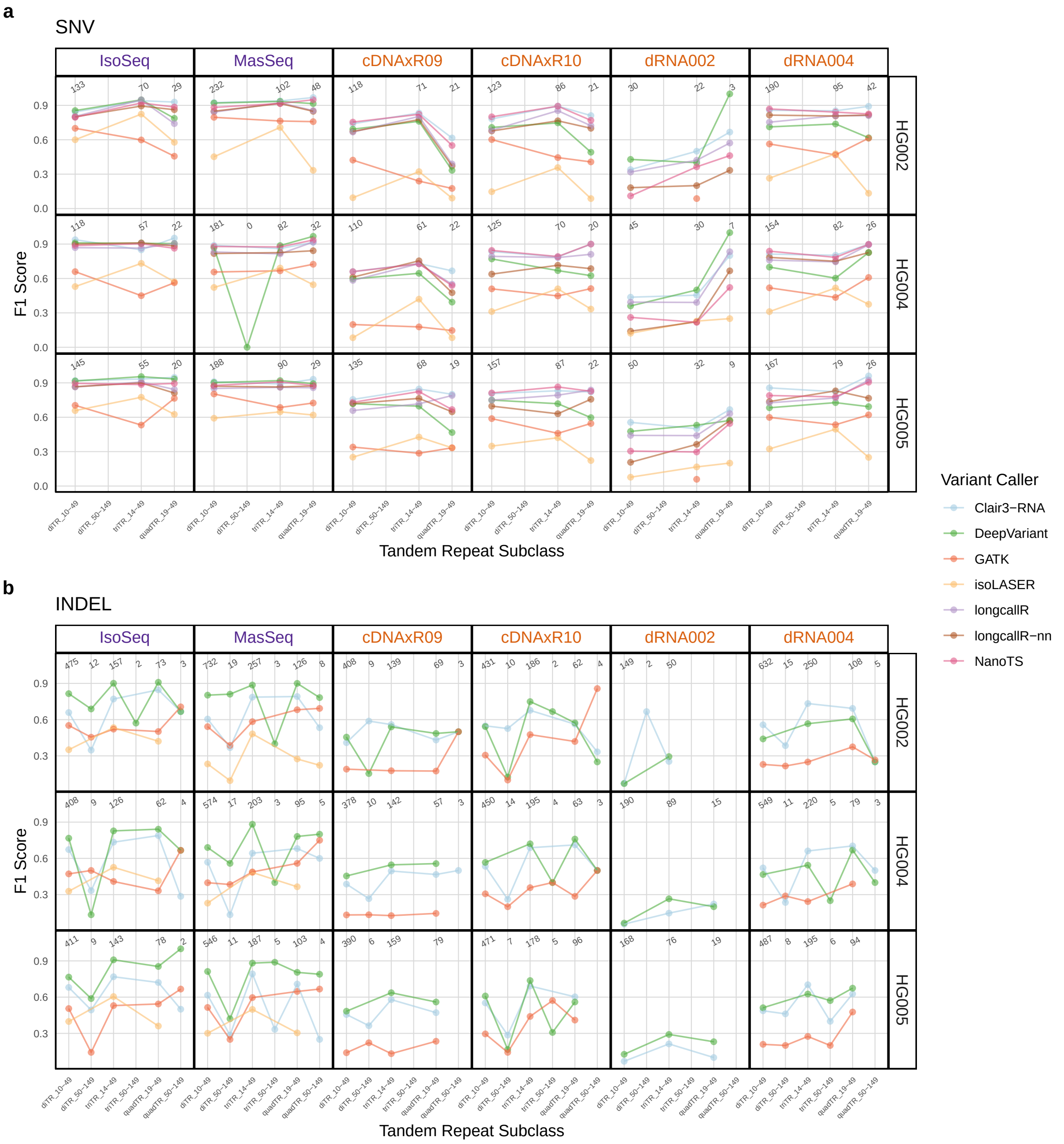

**Fig. S24: Variant calling performance based on F1 scores across short tandem repeats subclasses.** F1 scores for SNV (**a**) and INDEL (**b**) calling within STR subclasses. Rows = cell lines, columns = library preparation kits. For short tandem repeats, diTR, triTR, and quadTR denote di-, tri-, and quad-nucleotide tandem repeats, respectively, and the numeric ranges (e.g., 10–49, 50–149) indicate the total repeat length in base pairs.

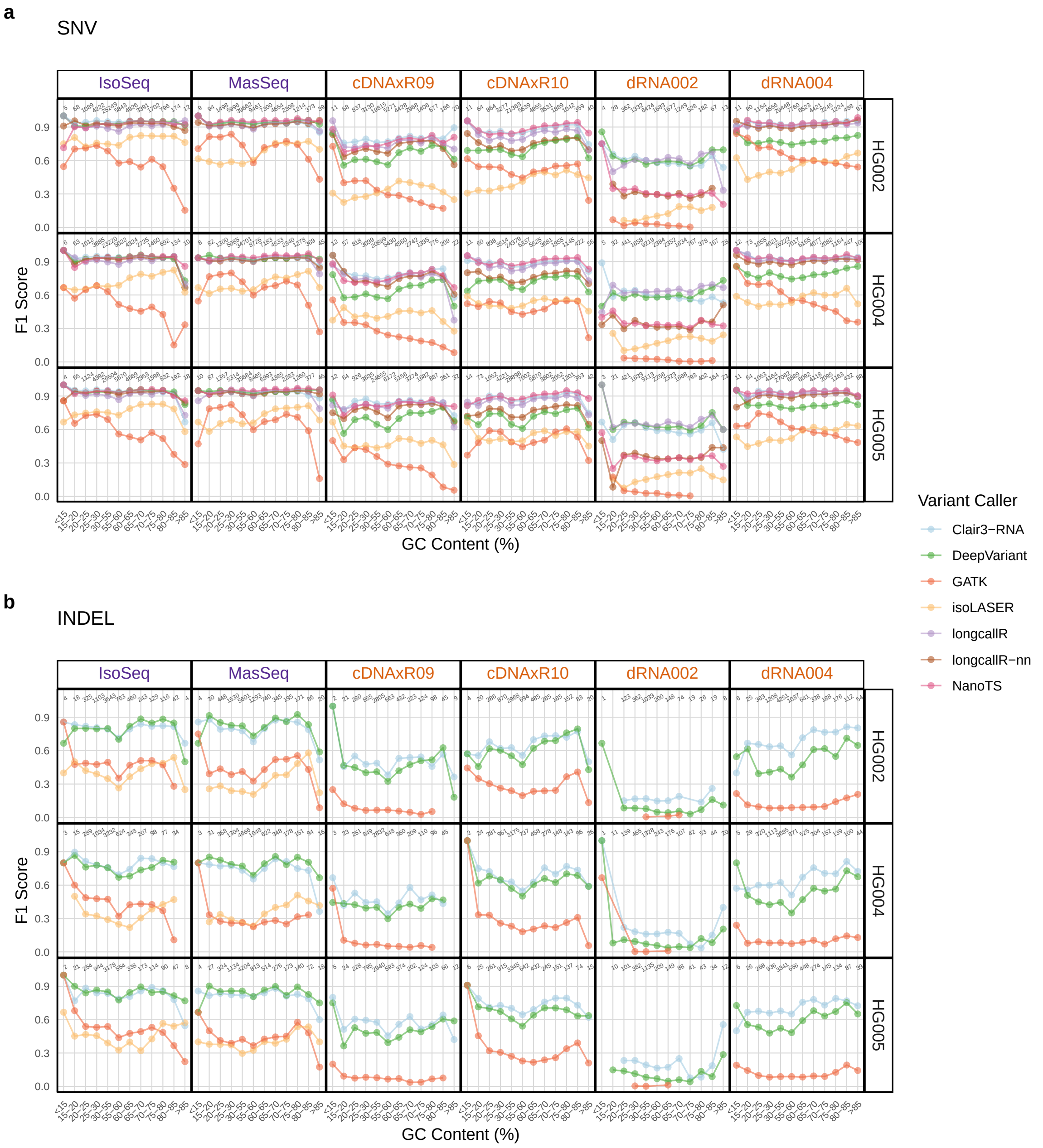

**Fig. S25: Variant calling performance based on F1 scores across different GC contents.** F1 scores for SNV (**a**) and INDEL (**b**) calling across different GC contents. Rows = cell lines, columns = library preparation kits.

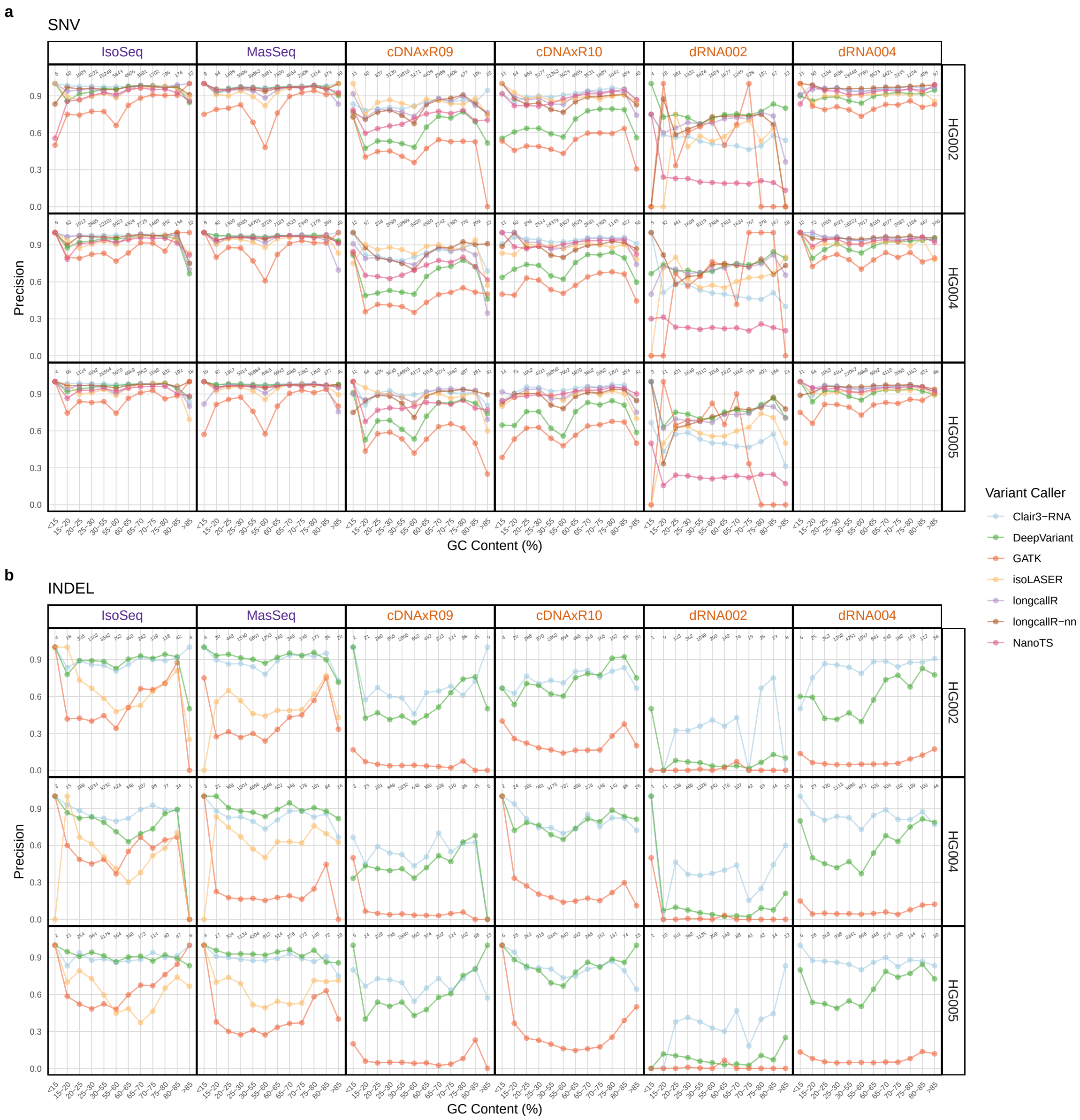

**Fig. S26: Variant calling performance based on precision across different GC contents.** Precision for SNV (**a**) and INDEL (**b**) calling across different GC contents. Rows = cell lines, columns = library preparation kits.

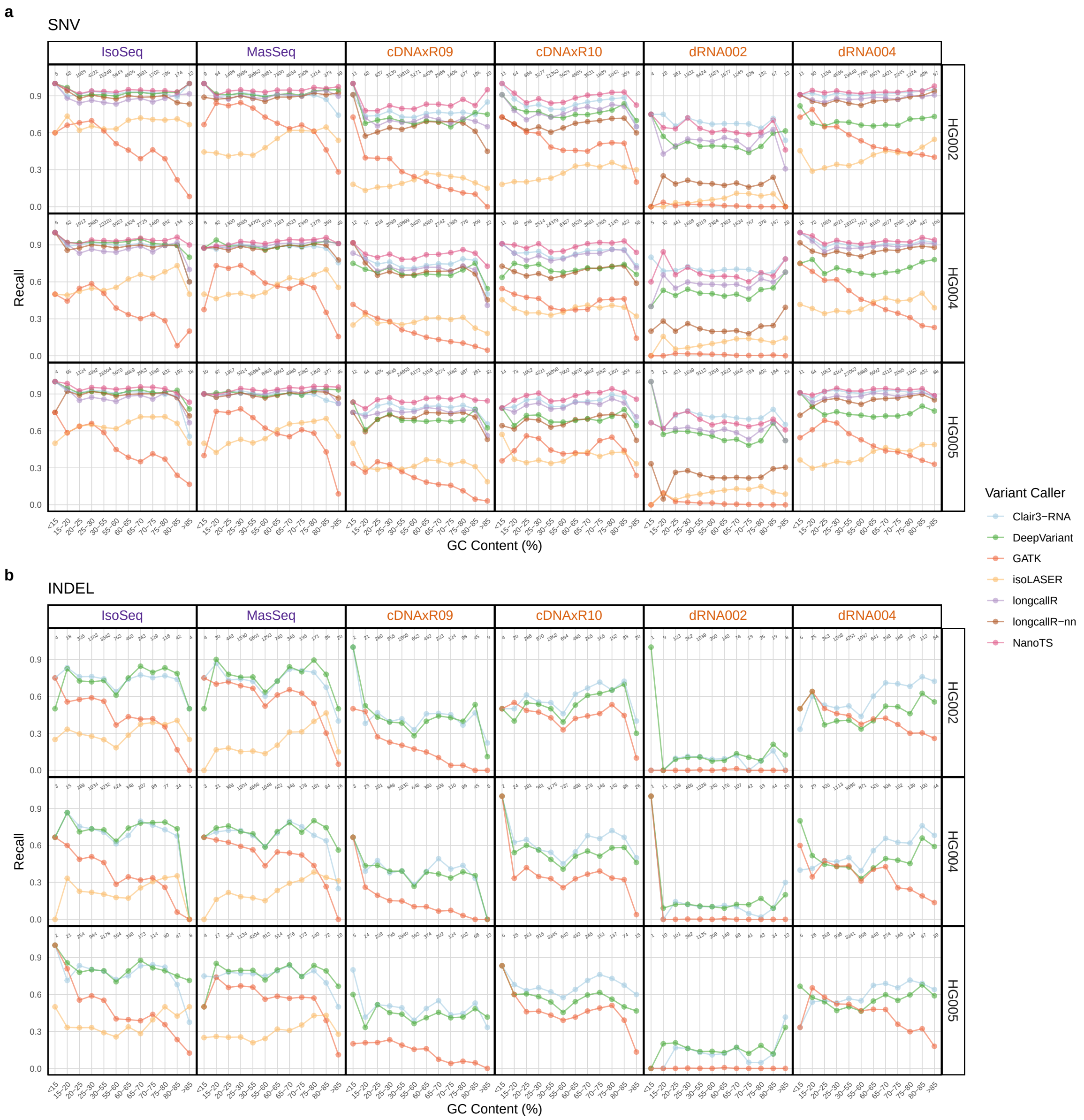

**Fig. S27: Variant calling performance based on recall across different GC contents.** Recall for SNV (**a**) and INDEL (**b**) calling across different GC contents. Rows = cell lines, columns = library preparation kits.

#### 2.5 LongBench datasets

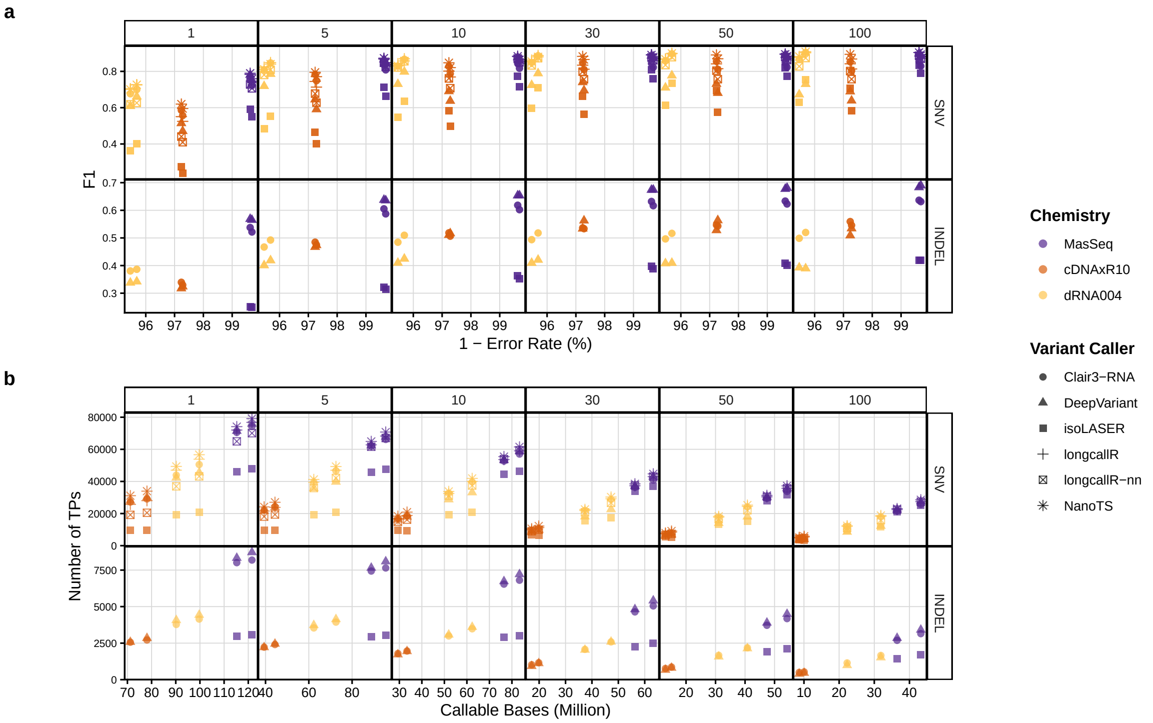

**Fig. S28: Impact of sequencing characteristics on variant calling performance in the LongBench datasets.** (**a**) Alignment-derived error rate versus F1 scores for each variant caller. (**b**) Number of callable bases versus the number of TP variant calls returned by each variant caller. Rows = SNVs and INDELs.

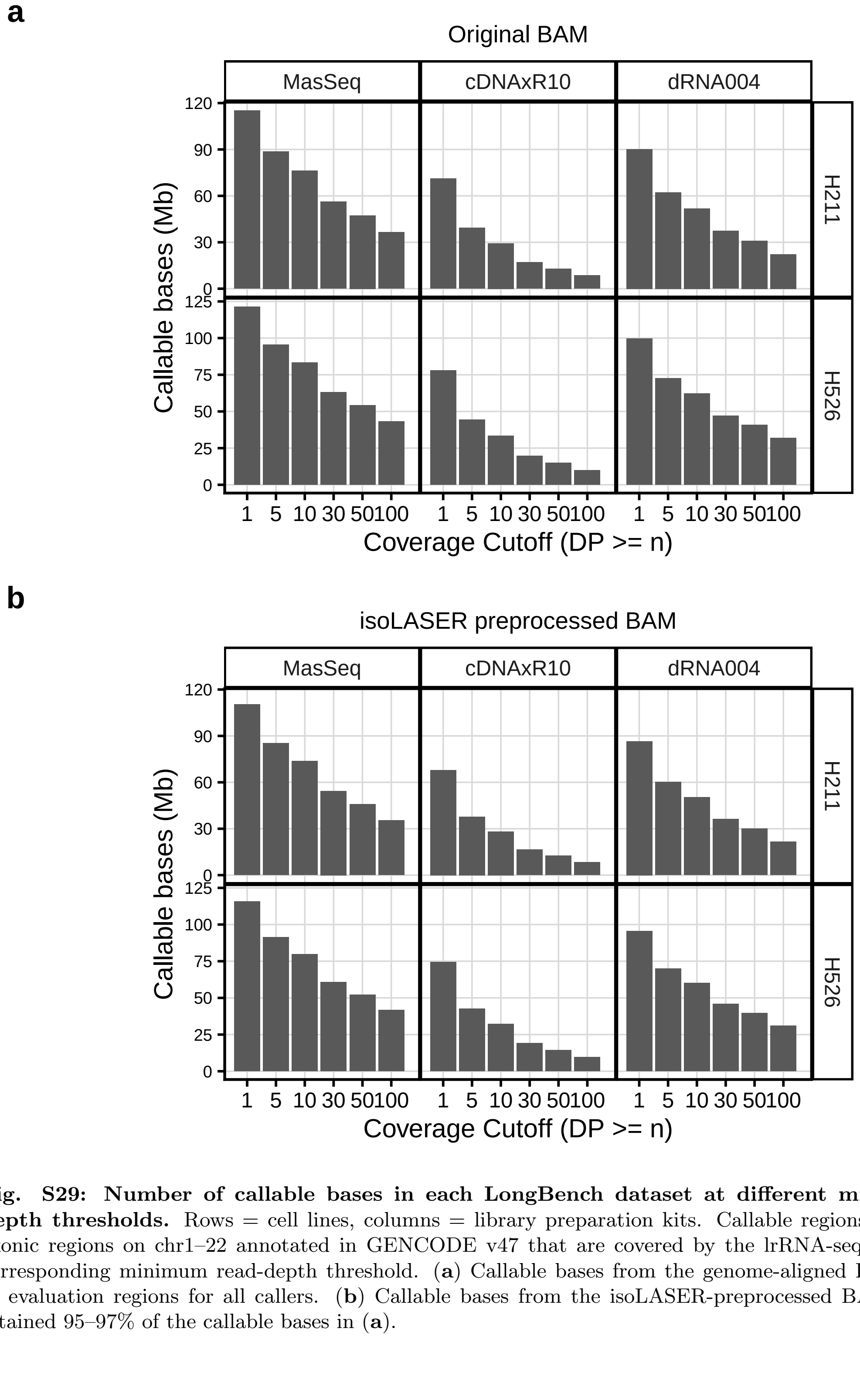

**Fig. S29: Number of callable bases in each LongBench dataset at different minimum read-depth thresholds.** Rows = cell lines, columns = library preparation kits. Callable regions are defined as exonic regions on chr1–22 annotated in GENCODE v47 that are covered by the lrRNA-seq dataset at the corresponding minimum read-depth threshold. (**a**) Callable bases from the genome-aligned BAM files, used as evaluation regions for all callers. (**b**) Callable bases from the isoLASER-preprocessed BAM files, which retained 95–97% of the callable bases in (**a**).

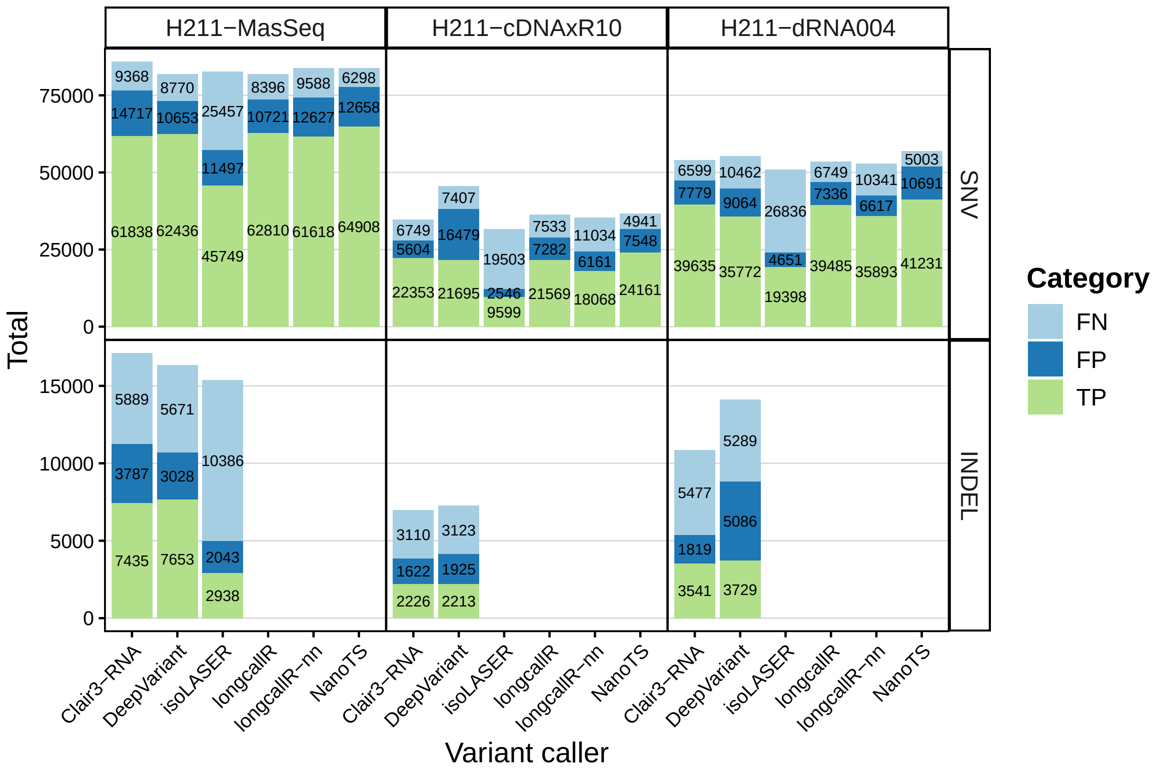

**Fig. S30: Number of TP, FN, and FP calls for each variant caller in the H211 datasets.**

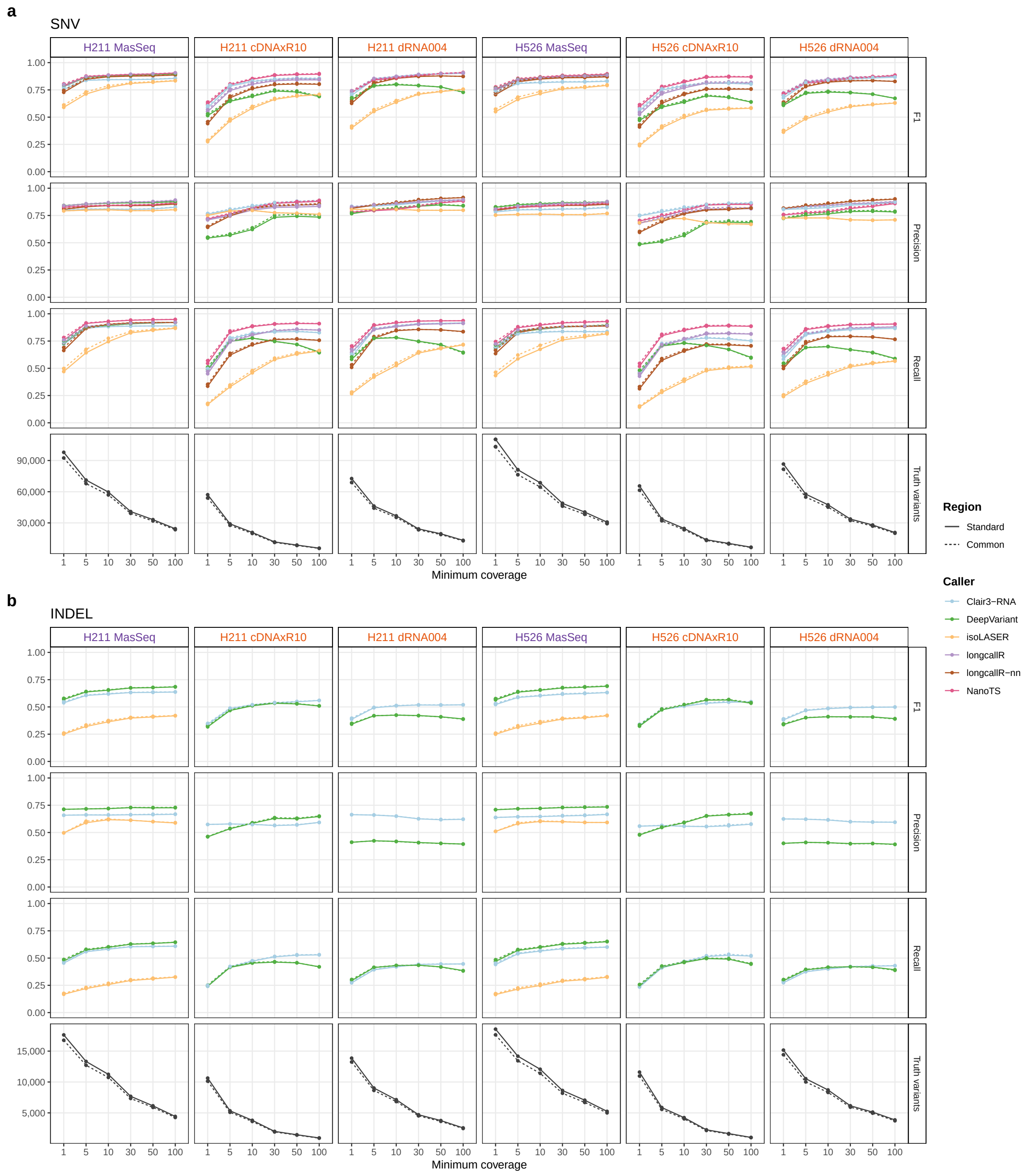

**Fig. S31: Variant-calling performance on regions covered after isoLASER preprocessing in LongBench datasets.** Variant calling performance for SNVs (**a**) and indels (**b**) evaluated on the common regions of the genome-aligned reads and the isoLASER-preprocessed reads. Truth variants indicate the number of true variants in ground truth set in the benchmarking region.

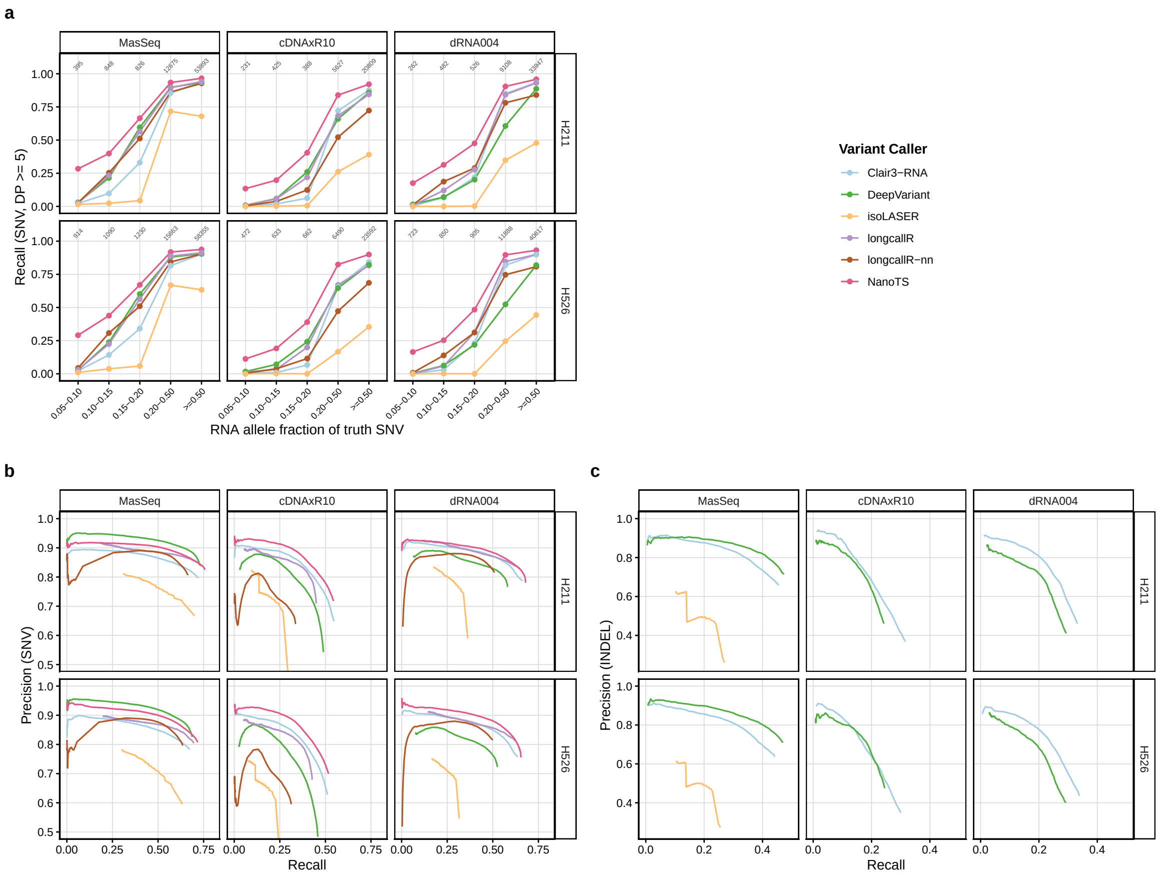

**Fig. S32: Robustness of caller performance to allele-fraction and quality-score thresholds on LongBench data.** (**a**) SNV recall stratified by the RNA allele fraction of each truth SNV at coverage $\geq 5$; numbers indicate the truth SNVs per bin. (**b**, **c**) Precision–recall curves for SNVs (**b**) and indels (**c**). In contrast to Supplementary Fig. X, curves were computed on all callable regions (coverage $\geq 1$), the confidence score of isoLASER was GQ, and calls failing only the score threshold (LowQual) were included in addition to PASS calls, so curves extend below the default thresholds where callers report LowQual calls (not longcallR-nn and NanoTS); thresholds retaining fewer than 100 calls are not shown. Curves of DeepVariant, longcallR and isoLASER start at non-zero recall because their scores are capped. The irregular curves of longcallR-nn reflect its narrow QUAL distribution (Supplementary Fig. X). Indel curves are shown for Clair3-RNA, DeepVariant and isoLASER (Mas-Seq only).

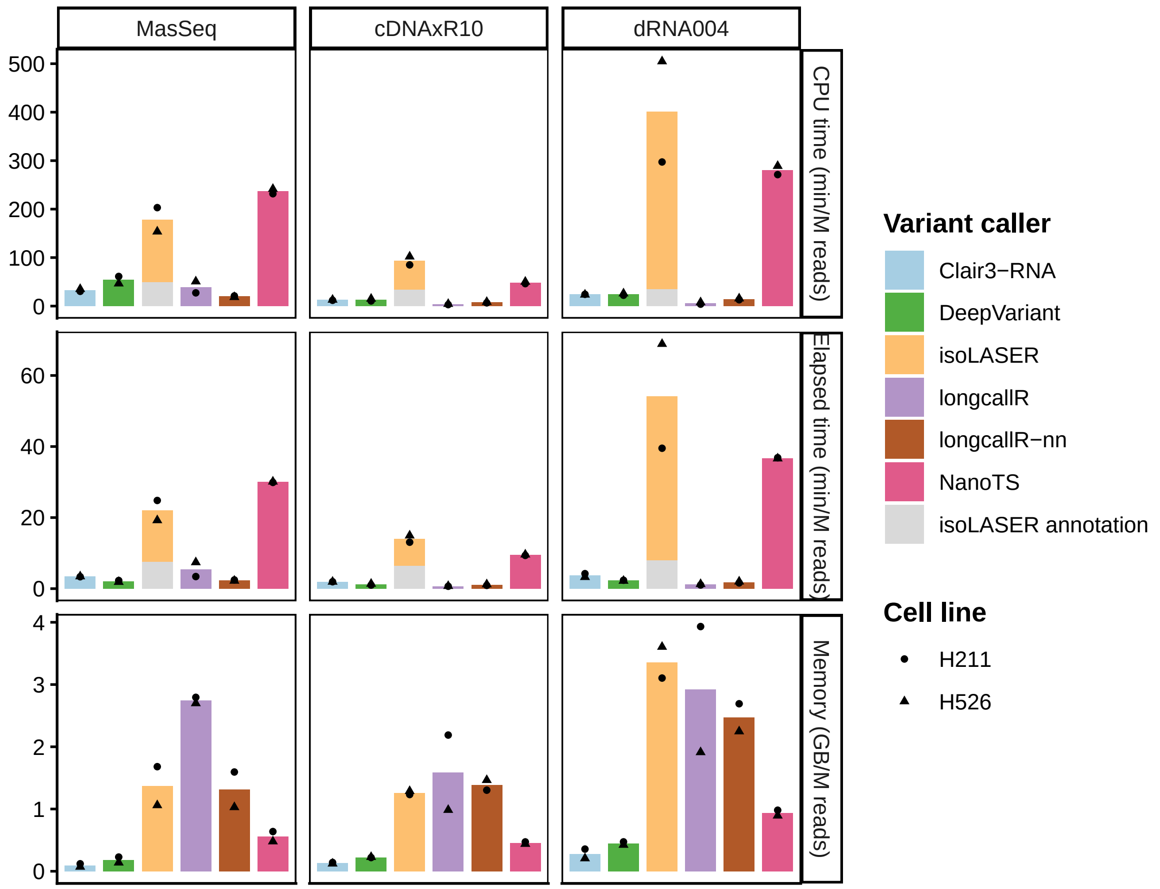

**Fig. S33: Computational costs per million reads on LongBench datasets.** Runtime, CPU time, and peak memory usage normalized per million reads, stratified by chemistries.
